# An in vivo chemical genetic approach for targeted glycoproteome analysis in *Drosophila melanogaster*

**DOI:** 10.64898/2026.09.15.751681

**Authors:** Sophie D. Schmidt, Cyrille Alexandre, Lucia Di Vagno, Liping Zhang, Helen Flynn, Joachim Kurth, Steve Murray, Sasha Ruiz Herrera, Hannah Shaw, Ganka Bineva-Todd, Ömür Y. Tastan, Mark Skehel, Nadine L. Samara, Kelly G. Ten Hagen, Jean-Paul Vincent, Benjamin Schumann

## Abstract

Glycans impact every aspect of physiology. Despite the rapid increase in relevance and translational potential, there remains a lack of tools for studying glycosylation *in vivo*. Here, we present FlyMOE, a chemical-genetic platform for targeted glycoprotein profiling in *Drosophila melanogaster*. Transgenic flies are equipped with biosynthetic capabilities to install bioorthogonally-tagged monosaccharides into the cellular glycoproteome. The genetic tractability of *Drosophila* allows for versatile application based on developmental stages, genetic drivers and glycosyltransferase enzymes. We employ FlyMOE to create an overview of the glycoproteome of *Drosophila* tissues and embryos based on mass spectrometry glycoproteomics, and investigate the substrates of individual members of the PGANT glycosyltransferase family *in vivo*. FlyMOE will allow the detailed investigation of the glycoproteome in *Drosophila* and similarly genetically modifiable *in vivo* model systems.

## Introduction

Glycosylation is one of the most abundant post-translational modifications and glycans play vital roles in protein function. Dysregulation of glycosylation is associated with a wide range of diseases ^1–3^. Unlike proteins, glycans are secondary gene products that are made by the complex interplay of over 200 glycosyltransferases (GTs) ^4,5^. Manipulation of GT gene expression in cells and animals has brought substantial insights into the intricacies of glycosylation in health and disease ^6–9^. While powerful, for instance in understanding the phenotypes upon loss of GT expression, such reductive methods rarely reveal the arsenal of glycoproteins that is made by cells, tissues or entire organisms. Furthermore, it is still challenging to correlate expression profiles of individual GTs with their protein substrates. Metabolic oligosaccharide engineering (MOE) has been used to address these shortcomings. Monosaccharides with bioorthogonal modifications are fed in a membrane-permeable form and, following biosynthetic activation via salvage pathways, incorporated into the cellular glycoproteome (Fig. 1a) ^10–13^. Bioorthogonal tags are then used to attach biotin or fluorophores for visualization and mass spectrometry (MS). A number of MOE reagents are analogues of *N*-acetylgalactosamine (GalNAc) ^10,14^. These compounds are generated by replacing the acetamide with bioorthogonal acylamide groups, such as azide-containing GalNAz and alkyne-containing GalNAlk (Fig. 1b). Their use in living cells relies on the biosynthetic conversion to the corresponding nucleotide-sugars. In mammals, the salvage pathway for GalNAc and some of its analogues consists of the kinase GALK2 and the pyrophosphorylase AGX1 that transform GalNAc to GalNAc-1-phosphate and then to UDP-GalNAc, respectively. Both enzymes have paralogues in mammals with different substrate specificities - GALK1 accepts galactose instead of GalNAc, and AGX1 is thought to bear specificity for *N*-acetylglucosamine (GlcNAc)-1-phosphate, although this notion has been contested ^15^. Once biosynthesized, UDP-GalNAc can be used as a substrate, for instance by a family of GalNAc-transferases (GalNAc-Ts) to initiate O-GalNAc glycosylation ^16,17^. Alternatively, UDP-GalNAc can be interconverted to UDP-GlcNAc by UDP-galactose-4-epimerase GALE (Fig. 1a) ^18,19^. GlcNAc is then incorporated into a range of glycan types including Asn(N)-linked glycans, extended Ser/Thr(O)-GalNAc and intracellular O-GlcNAc glycans. MOE has been heavily used to study glycosylation in cell lines^10,20,21^.

**Figure 1:**
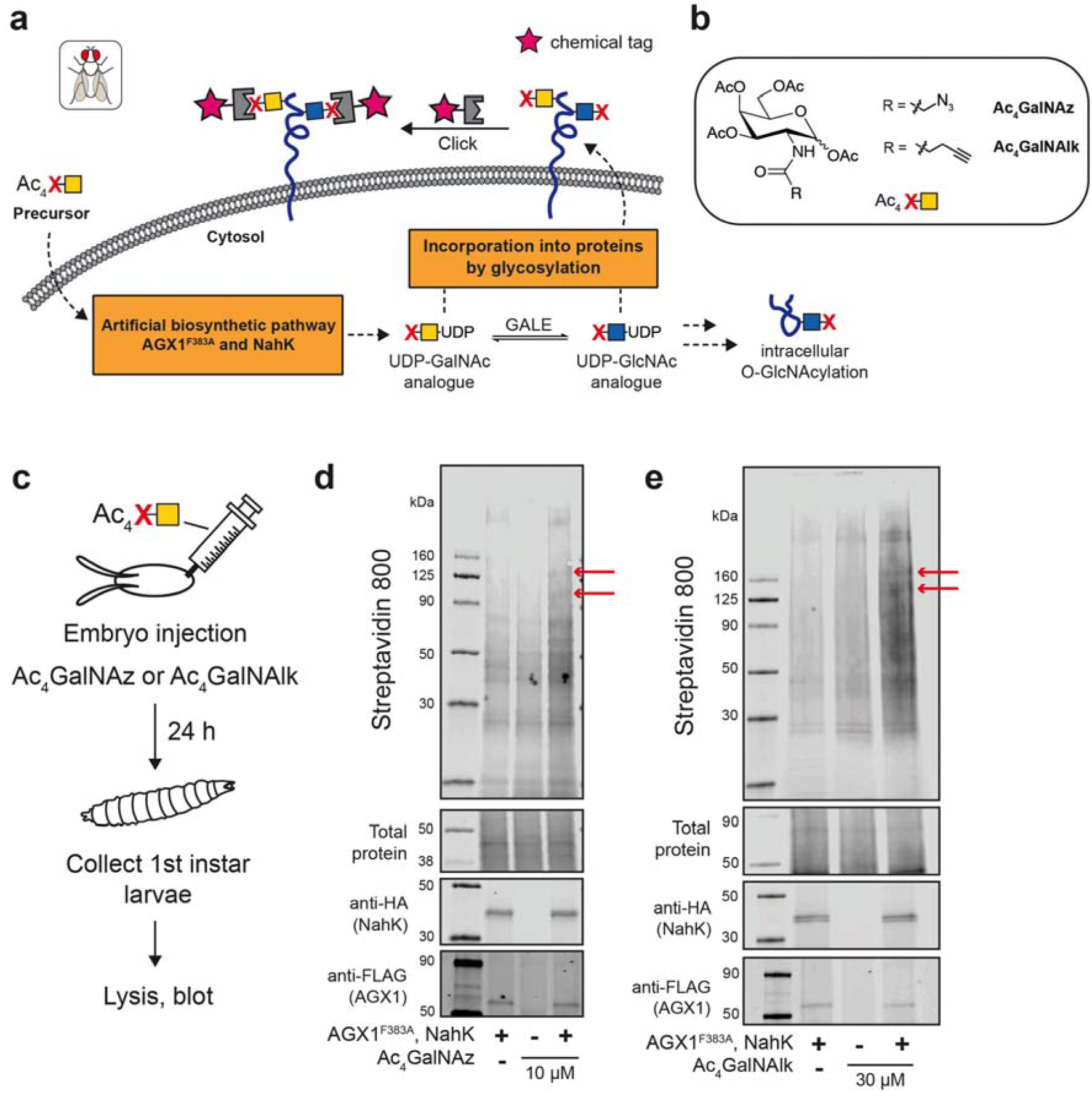
An artificial biosynthetic pathway allows for incorporation of chemically modified sugars Ac_4_GalNAz and Ac_4_GalNAlk by injection of embryos. a) Overview of FlyMOE. A chemically modified GalNAc analogue is delivered as a membrane-permeable peracetylated sugar. The modified sugar is sequentially activated to the GalNAc-1-P analogue and then the UDP-GalNAc analogue by the enzymes of the artificial biosynthetic pathway, AGX1^F383A^ and NahK. The nucleotide sugar can then be interconverted to the UDP-GlcNAc analogue by the epimerase GalE. Analogues of GalNAc and GlcNAc can be incorporated glycoproteins. The functional group (azide or alkyne) is then used in the CuAAC reaction to add a chemical tag (pink star) to label glycoproteins. b) Chemical structure of Ac_4_GalNAz and Ac_4_GalNAlk. c) Scheme of embryo injection with modified sugars. Transgenic embryos expressing NahK-HA and AGX1^F383A^-FLAG ubiquitously were injected 2 h after egg laying with either 10 µM Ac_4_GalNAz or 30 µM Ac_4_GalNAlk and left to develop into first instar larvae for 24 h. Surviving larvae were collected, lysed and treated with biotin under CuAAC conditions for analysis by streptavidin blot. d) Chemical tagging of the glycoproteome of embryos injected with Ac_4_GalNAz. Results are one representative out of two independent replicates. Red arrows denote new bands in lysates from Ubi-MOE larvae. e) Chemical tagging of the glycoproteome of embryos injected with Ac_4_GalNAlk. Results are one representative out of two independent replicates. Red arrows denote new bands in lysates from Ubi-MOE larvae.

The GalNAc salvage pathway in *Drosophila* is less well understood than in mammals. Flies have one orthologue of GALK1/2 (*Galk*) that utilizes galactose as a substrate, without proven activity for GalNAc ^22^. Similarly, the substrate specificity of the single *Drosophila* AGX1/2 orthologue Mummy for the possible substrates GalNAc-1-phosphate and GlcNAc-1-phosphate is ill-defined ^23,24^. A preferential source of UDP-GalNAc in *Drosophila* is therefore thought to be the epimerisation of UDP-GlcNAc by Gale ^22^.

Despite the immense potential to address questions in glycobiology, MOE has been sparsely used in *Drosophila*. Vocadlo and colleagues employed caged, per-acetylated GalNAz (Ac_4_GalNAz) added to food to tag glycans in larvae, suggesting that *Drosophila* metabolism is somewhat receptive towards GalNAc analogues ^25–27^. Chen and colleagues fed flies with non-acetylated GalNAz to tag O-GlcNAc proteins in adult heads, and changed their approach subsequently towards a post-lysis tagging protocol ^28^. We and others have found that *N*-acyl modifications hamper nucleotide-sugar biosynthesis in mammalian cells: While GalNAz is generally converted to UDP-GalNAz by GALK2/AGX1, even the smallest known alkyne derivative GalNAlk is not efficiently converted ^29,30^. For both analogues, conversion to the UDP-GalNAc analogues was substantially boosted by establishing an artificial salvage pathway ^29–31^. A bacterial kinase NahK and a mutant AGX1 with an enlarged active site (AGX1^F383A^) sequentially transform GalNAc analogues to the GalNAc-1-phosphate and UDP-GalNAc analogues, respectively. Since both enzymes can be encoded on the same gene cassette, artificial biosynthesis allows for efficient, genetically engineered MOE. We applied the technology as Bio-Orthogonal Cell-specific Tagging of Glycoproteins (BOCTAG) for GalNAc analogues, while Chen and colleagues used GlcNAc analogues in mice ^32,33^.

Beyond cell specificity, methods in chemical genetics have allowed the development of bioorthogonal tools for individual GT enzymes. In a glycosyltransferase bump-and-hole approach, we have engineered human GalNAc transferases to accept the alkyne-tagged UDP-GalNAc analogue UDP-GalN6yne ^34,35^. Informed by structural data, bulky Leu and Ile residues were replaced with Ala residues in the active sites of various GalNAc-T family members. The additional space accommodated the extended alkyne chain in UDP-GalN6yne, delivering an enzyme-specific bioorthogonal reporter strategy. Since UDP-GalN6yne can be biosynthesized in cells using NahK and AGX1^F383A^, a genetic strategy was devised in which all three enzymes were co-expressed. Feeding per-acetylated Ac_4_GalN6yne to these cells allowed for profiling of GalNAc-T-specific glycoprotein substrates ^34,36–38^. Similar to the human GalNAc-T isoenzymes, their structurally homologous *Drosophila* counterparts called PGANTs are involved in specific processes in physiology. For instance, PGANT3 and PGANT35A are highly expressed in imaginal discs of *Drosophila* larvae and differentially involved in mediating cell adhesion in the developing wing and epithelial tube formation, respectively ^39–44^. We have recently reported that PGANT9A and 9B are splice forms that are differentially expressed across larval tissues and exhibit distinct preferences for synthetic peptide and glycopeptide substrates ^45,46^. We reasoned that a direct reporter strategy for the activity of PGANT enzymes *in vivo* would be highly beneficial to further uncover the biology of O-GalNAc glycosylation in different tissues and developmental stages.

We present FlyMOE as a modular chemical-genetic platform to uncover *Drosophila* glycoproteins in a stage-, driver- and enzyme-specific fashion. Our approach was underpinned by the availability of genetic tools for straightforward genetic engineering and recent progress in developing chemical precision tools. We show that artificial biosynthesis can be installed with both ubiquitous and driver-specific expression to boost incorporation of bioorthogonal GalNAc analogues in *Drosophila*. We establish delivery modes for MOE reagents in fly embryo and larvae, and profile glycoproteomes through enrichment and mass spectrometry. Finally, we bump-and-hole engineer *Drosophila* PGANT isoenzymes to tag glycosyltransferase-specific substrates *in vivo* as an enhancement of FlyMOE. Our work establishes a programmable platform to investigate the *in vivo* glycobiology of *Drosophila* as a genetically tractable model system.

## Results

### Artificial biosynthesis for programmable chemical glycoproteome tagging in *Drosophila melanogaster*

To implement FlyMOE *in vivo*, we first generated transgenic flies expressing NahK and AGX1^F383A^ separated by a P2A peptide ubiquitously and constitutively using a previously designed “Ubi” promoter (Ubi-MOE) ^47^. We first confirmed that our artificial biosynthetic pathway does not substantially alter the *Drosophila* glycome, as blots with the lectins Concanavalin A (ConA), Soybean agglutinin (SBA), *Dolichos biflorus* agglutinin (DBA), *Vicia villosa* lectin (VVL), *Maackia amurensis* lectin II (MALII), *Sambucus nigra* lectin (SNA) and Aleuria aurantia lectin (AAL) showed little or no changes between larval lysates (Supplementary Fig. 1). We injected Ubi-MOE *Drosophila* embryos with Ac_4_GalNAz or Ac_4_GalNAlk (Fig. 1b, c). DMSO was used as a vehicle-control, and embryos from non-transformed sibling controls (w^11^^18^ “WT” strain) were injected with the same MOE reagents as further controls. Embryos were left to develop into first instar larvae for a day before collection and lysis. CuAAC with biotin probes revealed an approximate 1.5-2-fold increase in streptavidin signal over controls when NahK and AGX1^F383A^ were expressed (Fig. 1d, e). Furthermore, new bands at higher molecular weight (∼100-130 kDa) were visible in lysates from Ubi-MOE larvae, indicating successful incorporation of GalNAz and GalNAlk into the glycoproteome. Artificial biosynthesis was the basis for genetically programmable, chemical glycoprotein tagging.

We next expanded the scope of compound delivery and genetic tractability in FlyMOE. Compound feeding was employed as an alternative monosaccharide delivery method Fig. 2) ^25,26^. We placed transgenic, adult Ubi-MOE flies on media containing Ac_4_GalNAz and allowed them to lay eggs overnight (Fig. 2a). Following development into third instar wandering larvae on these media, lysates were then collected from all tissues except fat body, and CuAAC with biotin was conducted. Increasing the concentration of Ac_4_GalNAz in media led to a gradual increase in signal of Ubi-MOE lysates over WT lysates (approximate 1.3-fold increase for 100 µM, 1.8-fold for 300 µM and 2.5-fold for 500 µM) (Fig. 2c). However, slight toxicity was observed at 500 µM Ac_4_GalNAz with fewer adults emerging after pupation. We therefore chose 300 µM Ac_4_GalNAz for any future experiments. To test whether GalNAz was distributed across the whole larva, we analysed uptake and incorporation into different larval tissues (Fig. 2d-g). We found a significant increase in streptavidin signal separately in isolated gut and CNS/imaginal disc tissue from Ac_4_GalNAz-fed Ubi-MOE larvae over controls (Fig. 2g). New bands for chemically tagged glycoproteins at ∼90-150 kDa were observed in Ubi-MOE lysates. Finally, we tested the inducible GAL4/UAS system with FlyMOE (Fig. 2h). The *tubulin*-GAL4 (*tub*GAL4) driver was used to induce expression of NahK and AGX1^F383A^ constitutively across all tissues, mimicking the Ubi driver. A 1.6-1.9-fold increase in signal was observed over controls with new bands appearing at ∼100 kDa (Fig. 2h). Taken together, our data show that FlyMOE provides genetically programmable incorporation of GalNAz with a choice of compound delivery and incorporation workflows.

**Figure 2:**
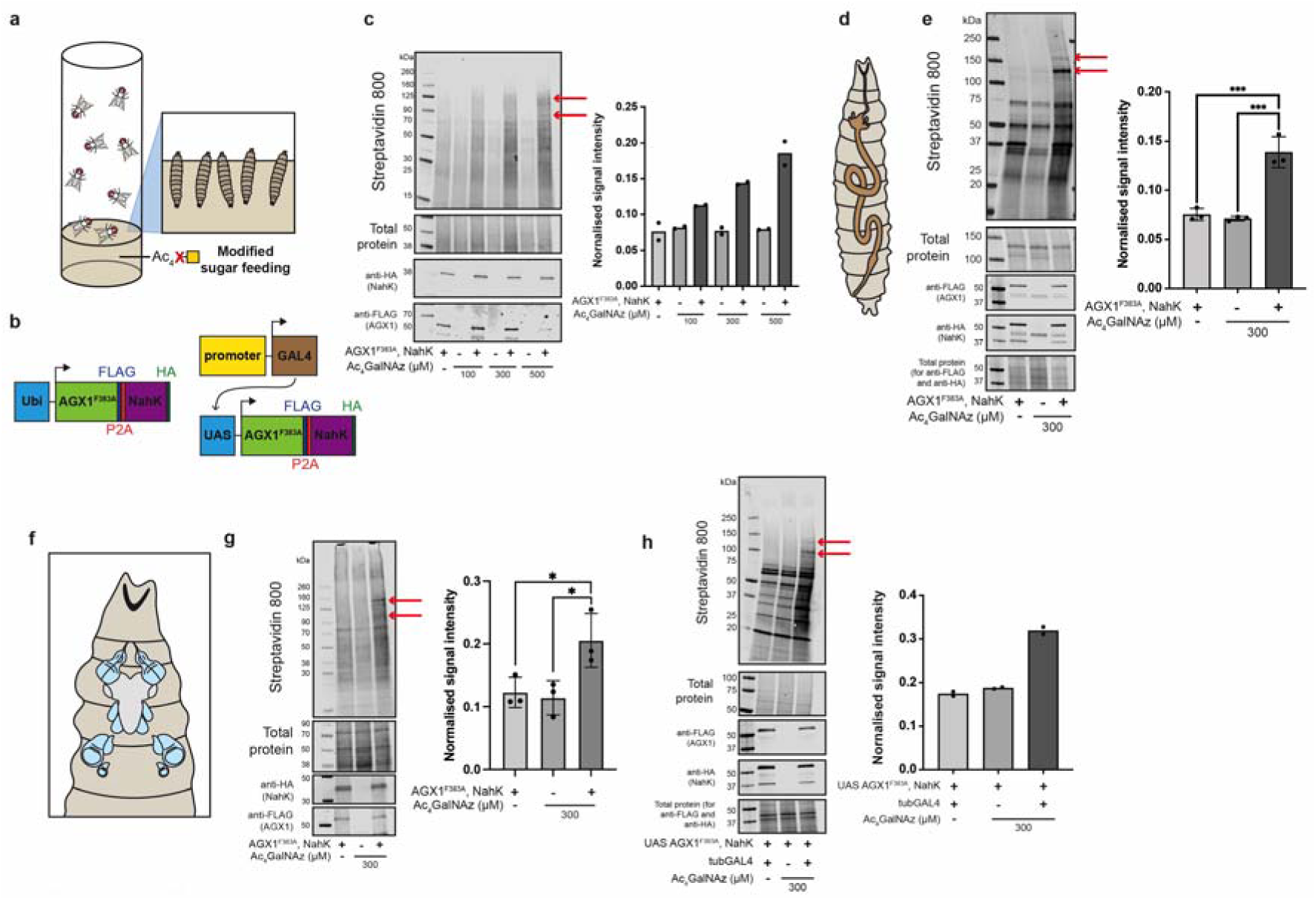
Incorporation of Ac_4_GalNAz into the glycoproteome of larval tissues by feeding. **a)** Transgenics expressing the enzymes of the artificial biosynthetic pathway were placed on food containing Ac_4_GalNAz. Flies were allowed to lay eggs and resulting third instar wandering larvae were collected and tissues were lysed for subsequent CuAAC reaction with biotin and analysis by streptavidin blot. **b)** Transgenics were generated either expressing AGX1^F383A^ and NahK ubiquitously throughout development and across tissues (Ubi driver, left) or using the GAL4/UAS driver system for tissue- or cell-specific expression with a GAL4 promoter (UAS driver, right). **c)** Chemical tagging of the glycoproteome of all tissues except the fat body of wandering larvae (Ubi) fed with Ac_4_GalNAz (100, 300 or 500 µM) (left) and quantification of normalised signal intensity of each lane (right). Results are one representative of two independent replicates. **d)** Scheme of the gut of *Drosophila* larvae. **e)** Chemical tagging of the glycoproteome of the gut of wandering larvae (Ubi) fed with 300 µM Ac_4_GalNAz (left) and quantification of normalised signal intensity of each lane (right). Data are individual data points and means ± SD of n = 3 independent replicates. Asterisks represent statistical significance: ns not significant; * *P* < 0.05; ** *P* < 0.01; *** *P* < 0.001. Results are one representative out of three independent replicates. **f)** *Drosophila* imaginal discs and central nervous system (CNS). **g)** Chemical tagging of the glycoproteome of the imaginal discs and CNS of wandering larvae (Ubi) fed with 300 µM Ac_4_GalNAz (left) and quantification of normalised signal intensity of each lane (right). Data are individual data points and means ± SD of n = 3 independent replicates. Asterisks represent statistical significance: ns not significant; * *P* < 0.05. **h)** Chemical tagging of the glycoproteome of all tissues except the fat body of wandering larvae (*tub*GAL4>UAS) fed with 300 µM Ac_4_GalNAz (left) and quantification of normalised signal intensity of each lane (right). The *tubulin* promoter drives the expression of AGX1^F383A^ and NahK in a constitutive manner. Results are one representative out of two independent replicates. Red arrows in all blots denote new bands in lysates from FlyMOE larvae.

### FlyMOE provides an overview of the glycoproteome of wing disc and embryos

We employed FlyMOE to provide insight into the *Drosophila* glycoproteome on different developmental stages. We employed our previously developed chemical glycoproteomics workflow in which glycoprotein samples are derivatized with acid-cleavable diphenyl disiloxane (DADPS)-containing biotin alkyne under CuAAC conditions. Enrichment on neutravidin beads allows for on-bead LysC digest to obtain a peptide fraction ready for analysis by data-dependent acquisition LC-MS/MS. Glycopeptides can be cleaved off beads using formic acid and further characterized. Chemical glycoproteome tagging in embryos was achieved by placing Ubi-MOE adults in cages on Ac_4_GalNAz-containing media and harvesting the resulting embryos. The peptide fraction revealed 17 proteins to be significantly enriched (Fig. 3b, Supplementary Table 1). These hits include intracellular nucleoporins Nup54, Nup58, Nup62, Nup98-96 and Nup214 that are known to be heavily O-GlcNAcylated ^48^, as well as the secreted, developmentally-regulated hemomucin (Hmu) glycoprotein that is O-GalNAc-glycosylated ^49^. Previously unannotated glycoproteins significantly enriched in our embryo dataset are Calmodulin (Cam), Cabeza (Caz), Lingerer (Lig), Tumbleweed (Tum), Microtubule-associated protein 205 (Map205) and Zipper (Zip). Because of known biosynthetic interconversions after Ac_4_GalNAz feeding, we expect these proteins to carry abundant glycan types such as O-GalNAc, O-GlcNAc and Asn(N)-linked glycans.

**Figure 3:**
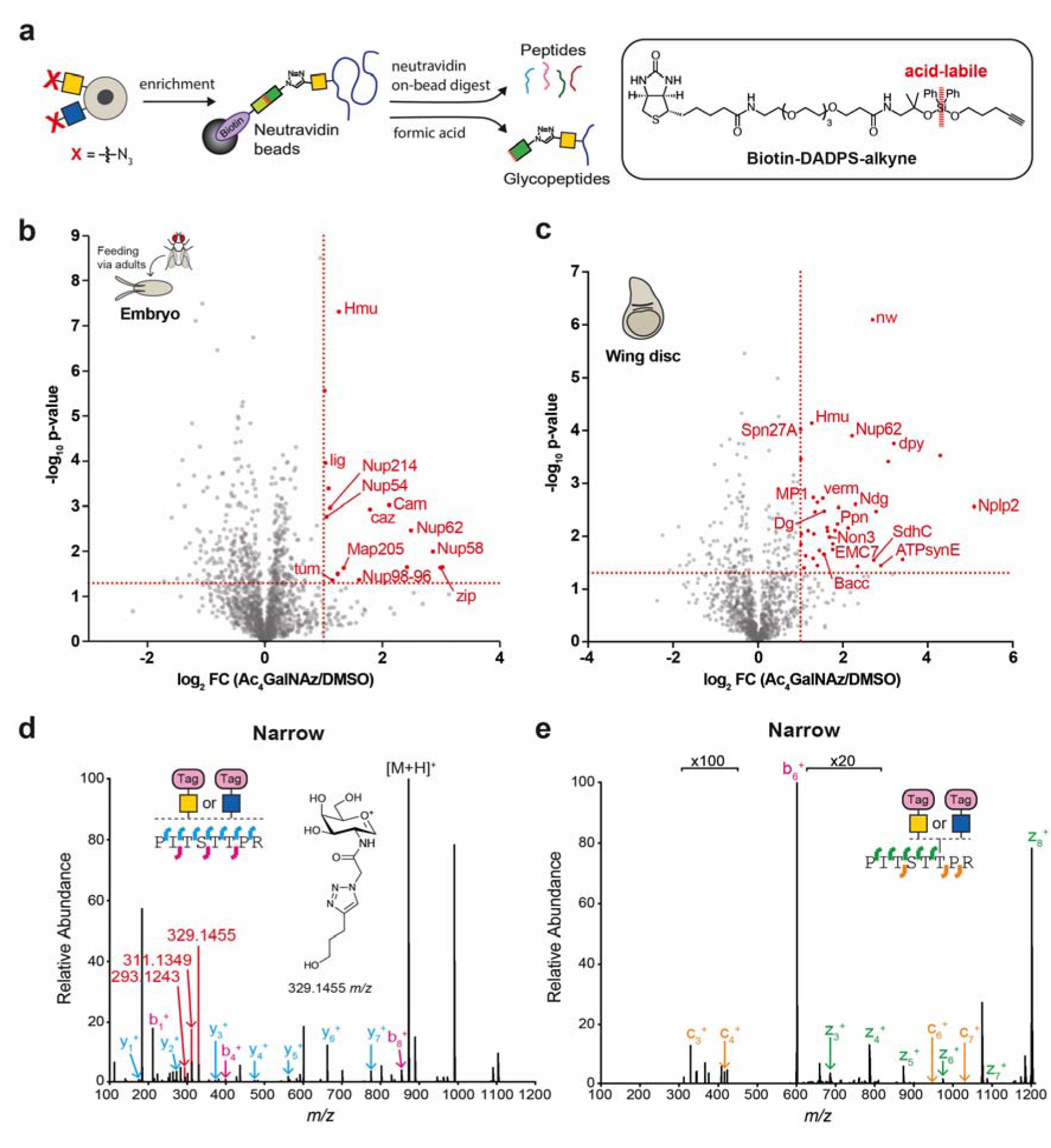
The glycoproteome of embryos and wing disc using FlyMOE. **a)** Lysates from different developmental stages of Ubi-MOE flies fed with Ac_4_GalNAz were subjected to CuAAC with biotin-DADPS-alkyne and enriched on neutravidin beads. On-bead digestion gives rise to a peptide fraction and glycopeptides were released by formic acid treatment. **b)** Proteome of Ubi-MOE embryos fed via adults. Results are from three biological replicates. **c)** Proteome of wing disc of third instar wandering Ubi-MOE larvae fed with 300 µM Ac_4_GalNAz or DMSO. Results are from two independent replicates. Results are shown as a volcano plot with a cutoff of 2-fold enrichment and p-value of <0.05. Enriched peptides were determined using two-sided two-sample Welch’s T-test and Benjamini-Hochberg to calculate adjusted p-values. For clarity, only enriched proteins with assigned gene names are shown; enriched CG-designated genes can be found in Supplementary Tables 1 and 2. For wing disc proteomics, ribosomal proteins are also not shown for clarity. **d)** HCD spectrum of Narrow (Nw) glycopeptide from wing disc glycoproteomics. Trigger ions (red) correspond to GalNAz (structure shown) or GlcNAz reacted with biotin-DADPS-alkyne, hydrolyzed through aqueous formic acid treatment and liberated oxonium ion (329.1455 *m/z*), with sequential loss of water (311.1349 *m/z* and 293.1243 *m/z*). Results are one from representative out two independent replicates. **e)** ETD spectrum of a Nw glycopeptide from wing disc glycoproteomics. Results are one representative out two independent replicates.

We next subjected wing discs from third instar wandering Ubi-MOE larvae fed with Ac_4_GalNAz or DMSO to our MS-glycoproteomics workflow (Fig. 3c, Supplementary Table 2). The peptide fraction revealed both known glycoproteins, including Nup62, Nidogen (Ndg), Papilin (Ppn), Hmu and Vermiform (Verm), and previously unannotated glycoproteins such as neuropeptide-like precursor 2 (Nplp2), ER membrane protein complex subunit 7 (EMC7), Dumpy (Dpy) and Narrow (Nw) to be significantly enriched in lysates from Ubi-MOE wing discs. These data suggest that a range of glycosylation types can be enriched through FlyMOE, and allow characterization of their associated glycoproteins. We then released the glycopeptide fraction associated with wing disc samples, and employed a tandem mass spectrometry approach to characterize peptide glycosylation. Higher Collision Dissociation (HCD) was used to observe the presence of the chemical modification on glycopeptides. A peptide from the extracellular matrix protein Narrow was found to be modified with a chemically modified HexNAc analogue (Fig. 3d). We then used Electron Transfer Dissociation (ETD) to mildly fragment the peptide backbone while leaving the attached glycan intact ^50^. Thr201 was unambiguously found to carry the HexNAc analogue on Nw (Fig. 3e). Nw is involved in the regulation of the size of the adult wing and footpad hairs ^51,52^. Although it is not possible from our data to conclude which HexNAc isomer is found on Nw, we note that Thr201 is part of a Thr/Pro-rich peptide that is reminiscent of mucin-type O-GalNAc glycopeptides ^53^. We conclude that FlyMOE allows mass spectrometry profiling of glycoproteins as well as individual glycosylation sites in *Drosophila*.

### FlyMOE to investigate glycosyltransferase specificity in *Drosophila melanogaster*

Understanding the intricacies of the glycoproteome requires precision methods to profile the products of individual glycosylation enzymes. We turned our attention to the family of PGANT glycosyltransferases that prime O-GalNAc glycosylation in *Drosophila*. We have previously employed the BH principle in engineering the human PGANT orthologues (the GalNAc-Ts) to accept a chemically modified UDP-GalNAc analogue termed UDP-GalN6yne (Fig. 4e). This approach was underpinned by artificial biosynthesis using NahK and AGX1^F383A^ to generate UDP-GalN6yne in cells. We reasoned that FlyMOE would provide a modular platform to visualize the *in vivo* glycoprotein substrates of individual PGANTs (Fig. 4a). Through sequence alignments with the human orthologues and overlays of our own crystal structures (Fig. 4b and Supplementary Fig. 2), we reasoned that replacing the bulky gatekeeper residues Ile and Leu in the active site of PGANTs to smaller Ala residues would accommodate the additional ‘bumped’ side chain in UDP-GalN6yne ^34,38^.

**Figure 4:**
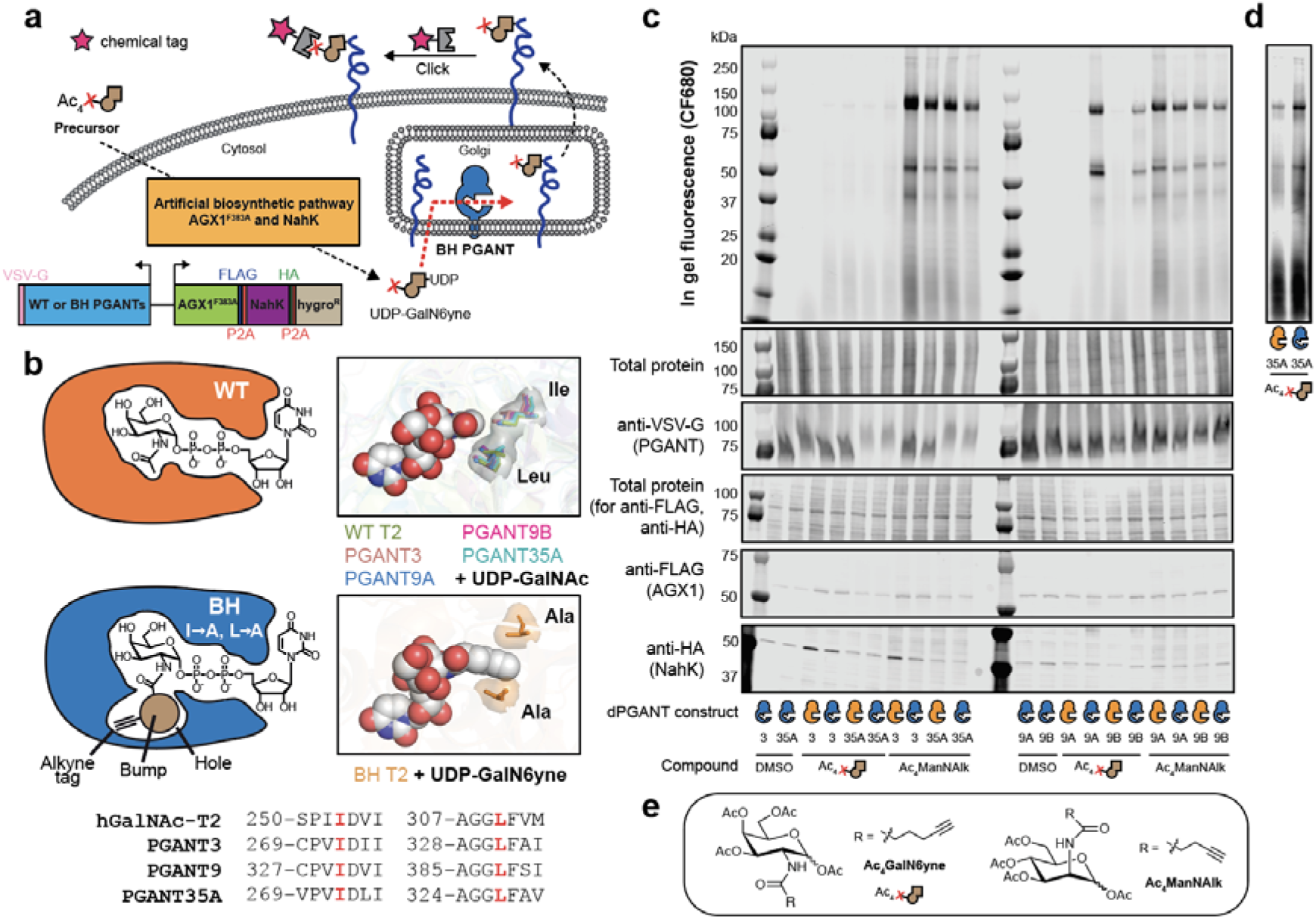
BH engineering on *Drosophila* PGANTs allows for detection of activity of individual glycosyltransferases. **a)** Overview of BH engineering. Ac_4_GalN6yne is delivered to cells and activated to UDP-GalN6yne using the artificial biosynthetic pathway made up of AGX1^F383A^ and NahK. UDP-GalN6yne is then used as a sugar donor for the BH engineered PGANT to glycosylate protein substrates. A chemical tag (pink star) can be used to detect protein substrates. **b)** Wild-type (WT) and Bump-and-Hole (BH) enzymes bound to UDP-GalNAc and UDP-GalN6yne. Crystal structure of WT GalNAc-T2 bound to UDP-GalNAc (PDB 4D0T), PGANT9A (PDB 6E4Q), PGANT9B (PDB 6E4R) and AlphaFold structure of PGANT3 (AF-Q9Y117-F1) and PGANT35A (AF-Q8MVS5-F1). Gatekeeper Ile residues are conserved between GalNAc-T2 and PGANT3, 9 and 35A. Crystal structure of BH GalNAc-T2 bound to UDP-GalN6yne (PDB 6NQT). Gatekeeper residues are mutated to Ala to accommodate the additional bump on UDP-GalN6yne. **c)** Cell surface labelling of human K-562 cells expressing *Drosophila* PGANTs (WT or BH). Cells were fed either DMSO (negative control), 2 µM Ac_4_GalN6yne or 10 µM Ac_4_ManNAlk (positive control). Lysates were treated with PNGase F prior to in-gel fluorescence to cleave N-glycans. Results are one representative out of three independent replicates. **d)** High intensity cut-out of WT and BH PGANT35A fed with 2 µM Ac_4_GalN6yne. **e)** Chemical structures of Ac_4_GalN6yne and Ac_4_ManNAlk.

Our workflow of engineering human GalNAc-Ts is contingent on detecting chemically modified glycoproteins on human K-562 leukemia cells as a straightforward cellular assay ^29,32,54^. To assess the fidelity of engineered PGANTs, we first used the same cell line as a heterologous host for *Drosophila* O-glycosylation. K-562 cells stably co-expressing *Drosophila* BH- or WT-PGANT isoenzymes 3, 9A, 9B or 35A along with NahK and AGX1^F383A^ were fed overnight with DMSO, Ac_4_GalN6yne or the sialic acid precursor Ac_4_ManNAlk as a positive control (Fig. 4e). CuAAC with the fluorophore CF-680 picolyl azide allowed visualization of cell surface glycoproteins by in-gel fluorescence (Fig. 4c). Out of the four isoenzymes chosen, BH variants of PGANT9A and 9B installed alkyne tags on K-562 cells with a substantial increase in fluorescence intensity over the corresponding WT enzymes (Fig. 4c). A characteristic banding pattern of cell surface mucin-type glycoproteins was seen in these samples ^32,34,38^. Expression of BH-PGANT35A led to a slight increase in fluorescence intensity over WT-PGANT35A in two out of three independent replicates (Fig. 4c, d, Supplementary Fig. 3). We note that expression levels of PGANT35A was generally lower than of PGANT9A and 9B. We did not see any visible increase of fluorescence signal in cells expressing BH-PGANT3 over WT-PGANT3 or DMSO controls.

Our data indicated that BH-variants of PGANT9A, 9B and, to a lesser extent, 35A, glycosylate proteins in the secretory pathway of living cells. Based on these data, we generated transgenic flies ubiquitously and constitutively expressing WT- or BH-PGANTs, NahK and AGX1^F383A^ as an enzyme-specific application of FlyMOE. Since PGANT9A and 9B are splice isoforms ^45^, we chose the more abundantly expressed PGANT9A to compare *in vivo* activity with PGANT35A. No changes were seen in the binding intensity of the lectin ConA that mainly probes terminal mannosides and glucosides (Supplementary Fig. 1a). Overexpression of WT-PGANT35A and 9A, but not the corresponding BH-PGANTs, induced an increase in binding intensity compared to WT flies (w^1118^ strain) of the lectins SBA, DBA and VVL that probe O-GalNAc glycans (Supplementary Fig. 1b-d). These findings confirm that overexpressed WT-PGANTs are able to use native UDP-GalNAc as a sugar donor and therefore alter glycosylation patterns. In contrast, BH-PGANTs require a bumped sugar to actively glycosylate substrates and thus do not affect the O-GalNAc glycome when overexpressed. There were no changes in binding pattern in any transgenic flies in MALII binding that probes a subset of α(2,3)-linked sialosides (Supplementary Fig. 1e) ^55,56^. We observed a slight change in binding pattern with the α(2,3)-sialic acid-specific lectin SNA when any PGANT was overexpressed (Supplementary Fig. 1g). This finding is in line with observations made in human cells where overexpression of GalNAc-T isoenzymes leads to a change in cellular sialylation (Ref: https://www.nature.com/articles/s41467-022-33854-0). No changes in binding to AAL that probes α(1,3)-fucosylation were observed in any transgenics compared to WT flies (Supplementary Fig. 1f).

To establish optimal metabolic engineering conditions, larvae expressing BH- or WT-PGANT35A were first fed with media containing 30, 100 or 300 µM Ac_4_GalN6yne and lysates were subjected to biotin picolyl azide under CuAAC conditions as stated above. Analysis by streptavidin blot indicated a slight increase in intensity and a specific band pattern >100 kDa when BH-PGANT35A was expressed compared to a WT-PGANT35A control (Supplementary Fig. 4a, c). These differences were more pronounced when an enrichment on neutravidin beads was performed before analysis (Fig. 5a). BH-PGANT35A-specific glycosylation signal increased with increasing concentration of Ac4GalN6yne in media. Since no toxicity of Ac_4_GalN6yne was observed, a 300 µM concentration was used in subsequent experiments. We next tested PGANT9A *in vivo* under these optimised conditions (Fig. 5b, Supplementary Fig. 4b, d). A general increase in streptavidin signal and new bands at ∼130 kDa and ∼90 kDa appeared in larvae expressing BH-over WT-PGANT9A. These data demonstrate that FlyMOE is suitable to chemically tag PGANT-specific protein substrates *in vivo*.

**Figure 5:**
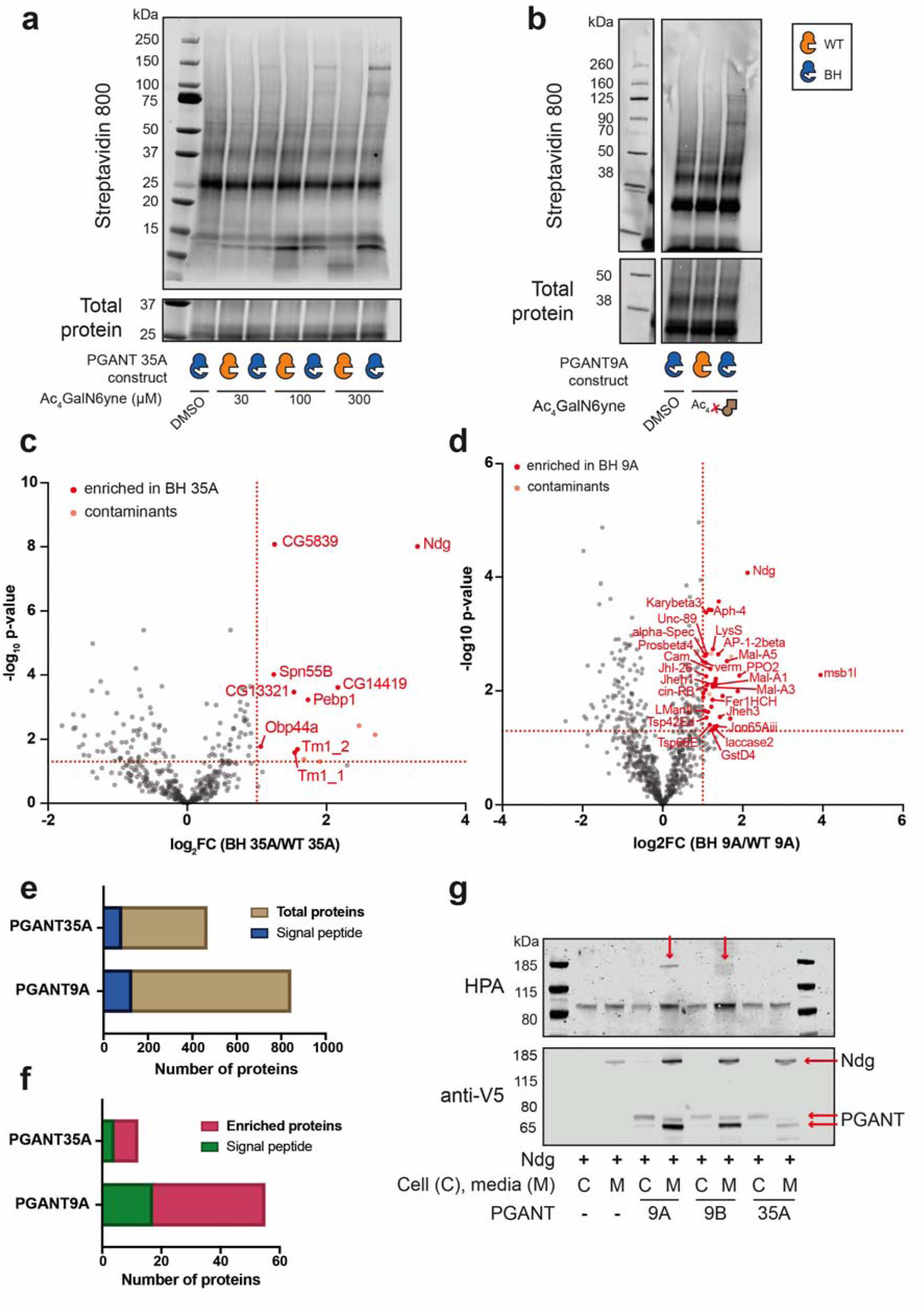
BH engineering allows for detection of protein substrates of PGANT35A and PGANT9A. **a)** Tagging of protein substrates of PGANT35A. Flies were fed with Ac_4_GalN6yne (30, 100, 300 µM) and all tissues except for fat body were collected for lysis and subsequent CuAAC with biotin. Clicked samples were enriched on neutravidin beads before analysis by streptavidin blot. Results are one representative out of three independent replicates. **b)** Tagging of protein substrates of PGANT9A. Flies were fed with 300 µM Ac_4_GalN6yne and all tissues except for fat body were collected for lysis and subsequent CuAAC with biotin. Clicked samples were enriched on neutravidin beads before analysis by streptavidin blot. Results are one representative out of three independent replicates. **c)** Proteomics to identify protein substrates of PGANT35A. Proteins enriched in BH-35A over WT-35A are shown in red. Results are from three independent replicates. **d)** Proteomics to identify protein substrates of PGANT9A. Proteins enriched in BH-9A over WT-9A are shown in red. Results are from three independent replicates. **e)** Proportion of proteins predicted to contain a signal peptide out of total number of proteins from proteomics experiments on whole larvae expressing BH-PGANT35A or 9A, NahK and AGX1^F383A^ fed with Ac_4_GalN6yne. PGANT35A: 82/384 (21.35 %); PGANT9A: 127/716 (17.74 %). **f)** Proportion of proteins predicted to contain a signal peptide out of enriched proteins from proteomics experiments on whole larvae expressing BH-PGANT35A or 9A, NahK and AGX1F383A fed with Ac_4_GalN6yne. PGANT35A: 4/8 (50 %); PGANT9A: 17/38 (44.74 %). Signal peptide predictions were performed using SignalP 6.0 (Teufel et al. 2022). **g)** HPA staining on purified proteins from media and cell lysates from *Drosophila* S2R+ cells co-expressing Ndg-V5 and either PGANT9A-V5, PGANT9B-V5 or PGANT35A-V5. Arrows in HPA blot show detection of terminal GalNAc on Ndg. Arrows in anti-V5 blot denote bands corresponding to Ndg or the PGANTs.

We conducted a chemical MS-proteomics experiment to identify protein substrates of BH-PGANT9A and 35A *in vivo*. We used lysates of whole larvae excluding fat body, installed DADPS-biotin picolyl azide by CuAAC and conducted on-bead digest. Peptides were analysed by quantitative data-independent acquisition (Fig. 5c, d). The software DIA-NN was used to identify and quantify proteins based on a whole-protein fasta file of *Drosophila melanogaster*. Statistical analysis was performed using Perseus ^57^. Excluding common contaminants, nine hits were found significantly enriched from larvae expressing BH-PGANT35A and fed with Ac_4_GalN6yne, of which three are products of poorly characterised computed genes (CG) (Fig. 5c, Supplementary Table 3). Five glycoproteins were predicted to contain a secretion signal peptide using SignalP 6.0 (DTU Health Tech) ^58^. Among the hits was the known secreted glycoprotein Nidogen (Ndg), an extracellular matrix protein. Remaining secreted proteins Serpin 55B (Spn55B), odorant binding protein 44a (Obp44A), CG14419 and CG5839, although these have not yet been annotated as glycoproteins. A total of 39 proteins were significantly enriched from larvae expressing BH-PGANT9A over WT, of which 17 are predicted to contain a secretion signal peptide. Of enriched protein hits, 14 are CG products (Fig. 5d, Supplementary Table 4). Hits contained the known glycoproteins Ndg and Verm. Notably, Ndg was identified as a substrate of PGANT9A, and therefore the only glycoprotein that was enriched after glycosylation by both PGANT35A and 9A. Ndg is a key component of the basement membrane that protects Collagen IV from degradation ^59–61^. Mammalian Ndg homologues (NID1/2 in humans and Nid1/2 in mice) are O-GalNAc-glycosylated, while no glycans have been annotated in the fly orthologue as of yet. To validate Ndg as a PGANT substrate, we co-expressed epitope tagged versions of WT-PGANT9A, 9B or 35A with a full-length Ndg construct in *Drosophila* S2R+ cells (Fig. 5g) ^62^. *Helix pomatia* agglutinin (HPA) blotting showed a band for terminal O-GalNAc glycosylation of Ndg when PGANT9A and 9B were expressed (Fig. 5g). Glycosylation of Ndg by PGANT35A was not detectable under our experimental conditions, possibly because expression of PGANT35A is lower than PGANT9A and 9B, and glycosylation of Ndg may be below detectable conditions.

## Discussion

The *Drosophila* model has led to some of the most important discoveries in physiology, enabled by a myriad of genetic constructs and strategies. We reasoned that a genetically enabled strategy for chemical tracing of glycosylation was a particular unmet need. Despite the long history of fly models for glycosylation, the design rules for both artificial biosynthesis of nucleotide-sugars and GT-specific reporter tools have been available for less than a decade ^29,34,63^. With advancements in sugar incorporation, the programmability of FlyMOE will allow combination with an extensive library of tissue- and cell-specific promoters. The GAL4/UAS system thus offers a platform to employ FlyMOE in profiling cell-type-specific glycosylation *in vivo*. Key to this advancement was the evaluation of administration techniques for MOE reagents. In addition to feeding larvae ^25,26^, we achieved metabolic glycan tagging of embryos either via injection or via feeding their mothers during oogenesis. FlyMOE is thus validated to profile the glycoproteomes of different tissues and developmental stages.

Our work establishes the first unbiased, chemically-informed glycoproteome datasets *in vivo* by mass spectrometry. We identified a range of proteins with various glycan types through the use of the promiscuous MOE reagent Ac_4_GalNAz. We and others have developed reagents that probe individual glycan types, for instance through their resistance to the epimerase GALE ^31,38,64,65^. Since artificial biosynthesis by NahK/AGX1^F383A^ is an enabling technology to these approaches, we expect FlyMOE to be compatible with alternative analogues of GlcNAc and GalNAc.

Our MS-glycoproteomics data allowed us to site-localise a modified glycan within a mucin-like peptide sequence on Nw. Annotation was aided by the modified sugar introducing a known mass shift in glycopeptides ^50,66,67^. FlyMOE thus lays a foundation to identify novel glycoproteins *in vivo* and study various glycosylation types.

Understanding the *in vivo* biology of O-GalNAc glycans is a longstanding question in *Drosophila* ^39,41,42,45,68–74^. The bump-and-hole approach initiated for human GalNAc-T isoenzymes was applicable to the *Drosophila* PGANTs based on sequence and structural homologies ^32,34,66^. With artificial biosynthesis underpinning UDP-GalN6yne biosynthesis, we suggest BH-PGANTs as an optional module in the FlyMOE platform. Implementation of this module identified Ndg to be a potential substrate for PGANT35A and PGANT9A. We reason that O-GalNAc glycosylation may regulate the stabilising function of Ndg within the basement membrane and influence its interaction with its binding partners. Ndg is conserved between *Drosophila* and humans where it is implicated in cancer progression, neurological disorders and congenital malformations ^75–77^. Our data suggest that *Drosophila* Ndg may be an adequate model system to study the role of human NID1/2 glycosylation. Since the gatekeeper residues are conserved in all *Drosophila* PGANTs, we suggest that that BH engineering is applicable, in principle, to other isoenzymes to shed light on their activity *in vivo* ^2,16,78^.

## Acknowledgements

We thank Iain Wilson, Erhard Hohenester, James MacRae and Alex Gould for their input. We thank Tania Auchynikava for help with data analysis. We thank the Francis Crick Institute Cell Sciences Platform for valuable help. This work was supported by the Francis Crick Institute (to B. S. and J.-P. V.), which receives its core funding from Cancer Research U.K. (CC2127 and FC001204), the U.K. Medical Research Council (CC2127 and FC001204), and Wellcome Trust (CC2127 and FC001204). This work was supported by UK Research and Innovation (UKRI) under the U.K. government’s Horizon Europe funding guarantee (grant number UKRI3602 to B.S.). We thank the Biotechnology and Biological Sciences Research Council (BB/V008439/1, BB/V014862/1 and APP23633 to B.S.). S. D. S. was supported by a Crick-HEI doctoral studentship between the Francis Crick Institute and the Department of Chemistry at Imperial College London. This work was supported in part by the Intramural Research Program of the NIDCR, National Institutes of Health (NIH) (1-ZIA-DE000754-03 to N.L.S. and Z01-DE000713 to K.G.T.H). The contributions of the NIH authors were made as part of their official duties as NIH federal employees, are in compliance with agency policy requirements, and are considered Works of the United States Government, However, the findings and conclusions presented in this paper are those of the authors and do not necessarily reflect the views of the NIH or the U. S. Department of Health and Humnan Sciences.

## Data availability

Proteomics and glycoproteomics data will be uploaded to ProteomeXchange through the MassIVE server. The data supporting the findings of this study are available within the paper and its Supplementary Information. Any additional raw data files are available from the corresponding authors upon reasonable request.

## Conflicts of interest statement

The authors declare no conflicts of interest.

## Notes

### Competing Interest Statement

The authors have declared no competing interest.

