## Supplementary Information for "An in vivo chemical genetic approach for targeted glycoproteome analysis in *Drosophila melanogaster*"

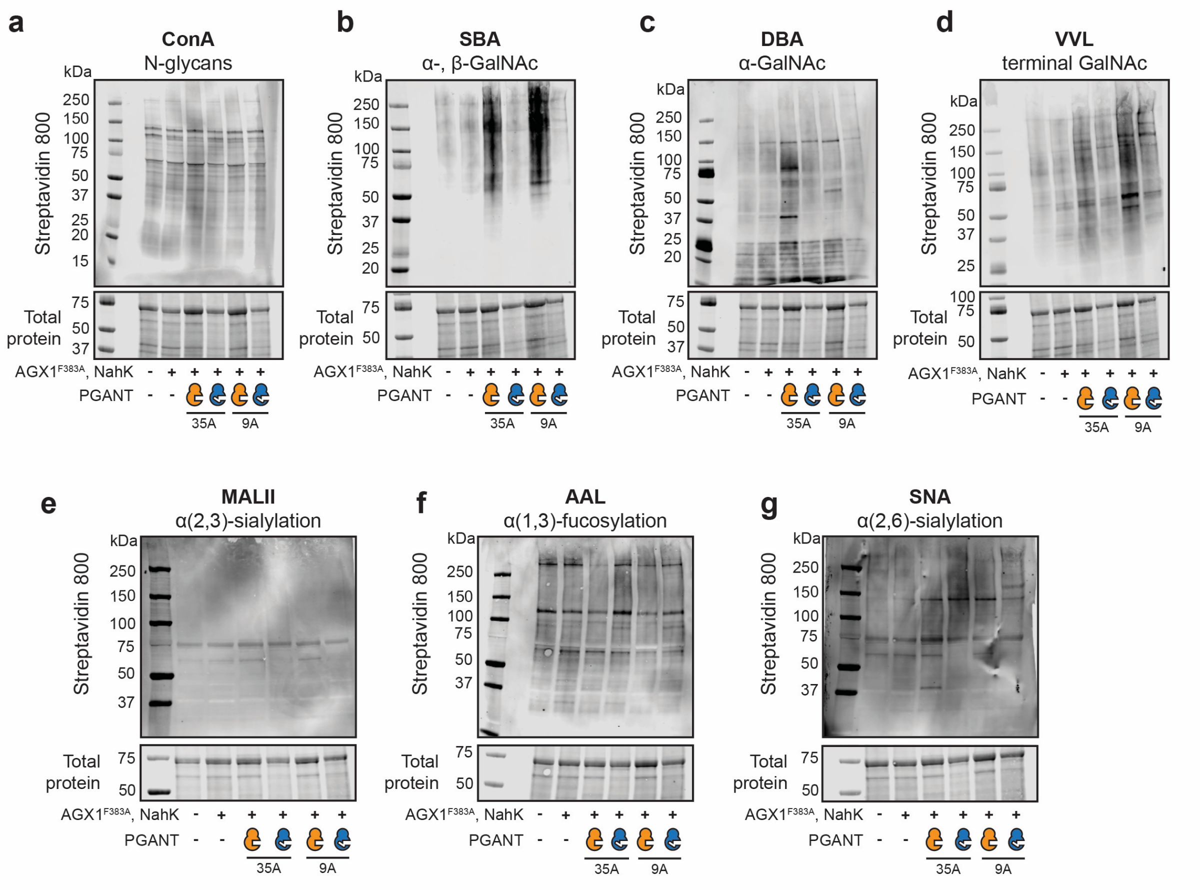


**Supplementary Fig. 1: Lectin blots on transgenic *Drosophila* to assess changes in the glycome.** Whole larval lysates (without fat body) of Ubi-MOE and Ubi-PGANT transgenics were collected and lysed for lectin blots. **a)** Concanavalin A (ConA). **b)** Soybean agglutinin (SBA). **c)** Dolichos biflorus agglutinin (DBA). **d)** Vicia villosa lectin (VVL) binding. **e)** Maackia amurensis lectin II (MALII). **f)** Aleuria aurantia lectin (AAL). **g)** Sambucus nigra lectin (SNA). Each blot is one representative out of three independent replicates.

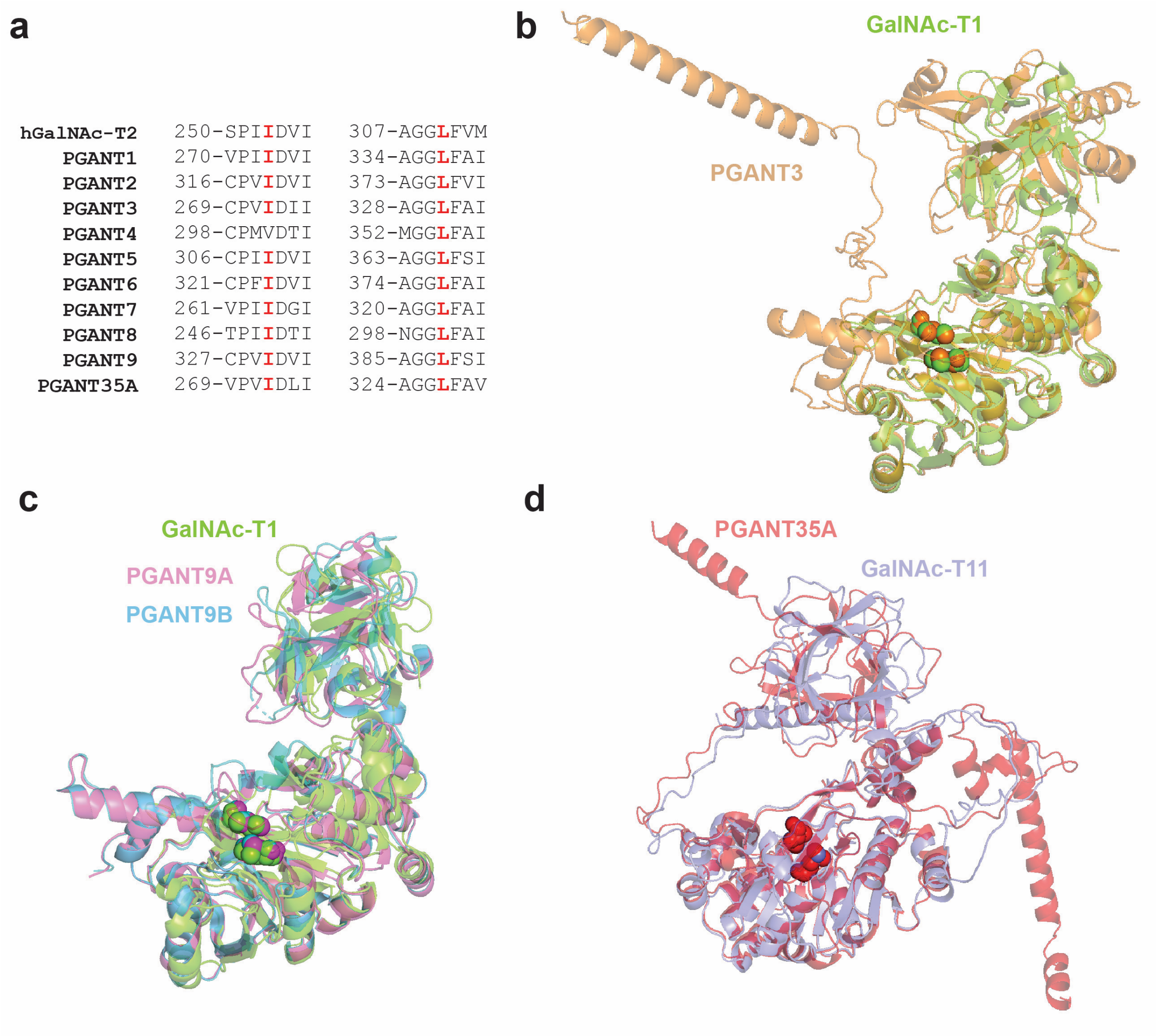


**Supplementary Fig. 2: Sequence alignment of *Drosophila* PGANTs and human GalNAc-T2 and structural alignment of PGANTs with their human orthologues.** **a) b)** Crystal structure of GalNAc-T1 (green, PDB 1XHB) aligned with AlphaFold structure of PGANT3 (orange, AF-Q9Y117-F1). Conserved gatekeeper residues are shown as spheres: Ile238 and Leu295 (GalNAc-T1); Ile272 and Leu331 (PGANT3). **c)** Aligned crystal structures of GalNAc-T1 (green, PDB 1XHB), PGANT9A (pink, PDB 6E4Q) and PGANT9B (cyan, PBD 6E44R). Conserved gatekeeper residues are shown as spheres: Ile238 and Leu295 (GalNAc-T1); Ile331 and Leu388 (PGANT9A and 9B). **d)** Aligned AlphaFold structures of GalNAc-T11 (purple, AF-Q8NCW6-F1) and PGANT35A (red, AF-Q8MVS5-F1). Conserved gatekeeper residues are shown as spheres: Ile274 and Leu329 (GalNAc-T11); Ile272 and Leu327 (PGANT35A).

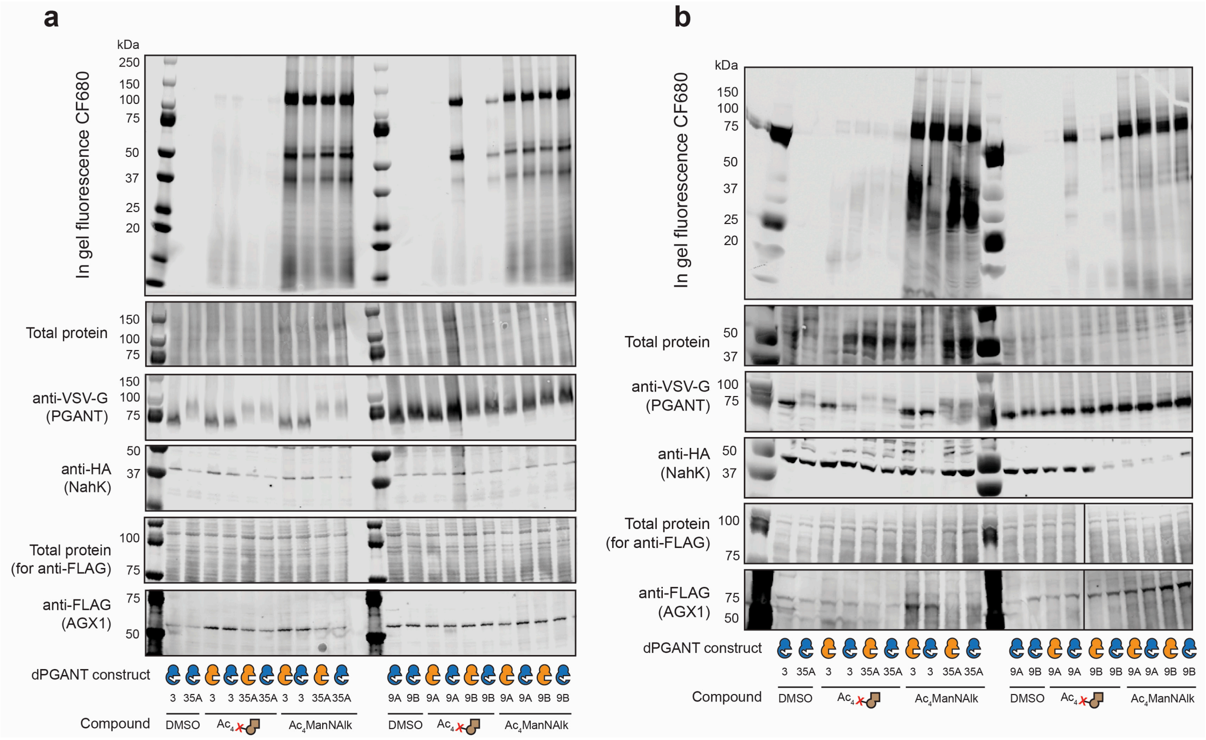


**Supplementary Fig. 3: Independent replicates of K-562 cell surface labelling.** Cell surface labelling of human K-562 cells expressing *Drosophila* PGANTs (WT or BH). Cells were fed either DMSO (negative control), 2 µM $Ac_4$GalN6yne or 10 µM $Ac_4$ManNAlk (positive control). Lysates were treated with PNGase F prior to in-gel fluorescence to cleave N-glycans. **a)** Replicate 1. **b)** Replicate 2.

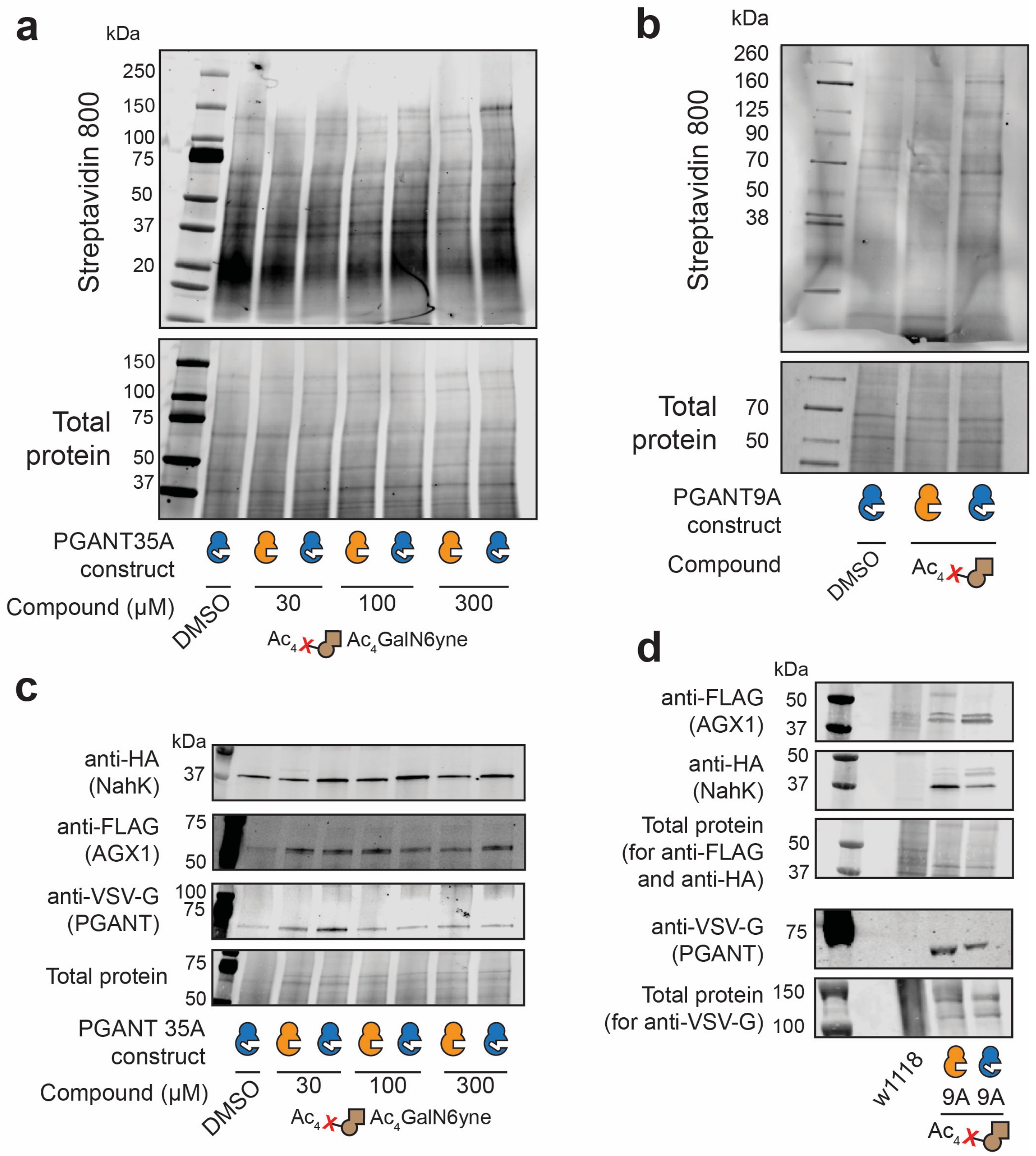


**Supplementary Fig. 4: BH engineering allows for detection of protein substrates of PGANT35A and PGANT9A. a)** Tagging of protein substrates of PGANT35A. Flies were fed with $Ac_4GalN6yne$ (30, 100, 300 μM) and all tissues except for fat body were collected for lysis, then CuAAC with biotin and subsequent streptavidin blot. Results are from one experiment. **b)** Tagging of protein substrates of PGANT9A. Flies were fed with 300 μM $Ac_4GalN6yne$ and all tissues except for fat body were collected for lysis, then CuAAC with biotin and subsequent streptavidin blot. Results are from one experiment. **c)** Western blotting against HA-NahK, FLAG-AGX1$^{F383A}$ and VSV-G-PGANT35A of lysates from whole larvae without fat body fed with $Ac_4GalN6yne$ (30, 100, 300 μM) or DMSO. Results are from one experiment. **d)** Western blotting against

HA-NahK, FLAG-AGX1$^{F383A}$ and VSV-G-PGANT9A of lysates from whole larvae without fat body fed with 300 μM $Ac_4GalN6yne$. $w^{1118}$ strain was used as a WT control. Results are from one experiment.

**Supplementary Table 1: List of enriched proteins from the peptide fraction of embryos fed with $Ac_4GalNAz$.** Function was taken from FlyBase [1]; cellular location was taken from UniProt [2]; annotated glycosylation was taken from GlyGen [3].

| **Gene name** | **Function** | **Cellular location** | **Glycosylation** |
|---|---|---|---|
| CG32795 | - | - | - |
| Zipper (zip) | Microtubule-binding protein involved in cytoskeleton-dependent intracellular transport. | Cell periphery | Unknown |
| Nucleoporin 58kD (Nup58) | Part of the nuclear pore complex | Nucleus | O-GlcNAc |
| Nucleoporin 62kD (Nup62) | Part of the nuclear pore complex | Nucleus | O-GlcNAc |
| CG14834 | - | - | - |
| Calmodulin (Cam) | Calcium-binding messenger protein | Cytoskeleton | Unknown |
| Cabeza (caz) | Chromatin binding protein involved in locomotion, synaptic growth at the neuromuscular junction and eye development. | Nucleus | Unknown |
| Nucleoporin 98-96kD (Nup98-96) | Part of the nuclear pore complex | Nucleus | O-GlcNAc |
| Microtubule-associated protein 205 (Map205) | Microtubule-associated protein 205. Binds and stabilises microtubules. | Cytoskeleton | Unknown |
| Hemomucin (Hmu) | Transmembrane mucin that may be involved in cellular adhesion and the innate immune response. | Endomembrane | N-glycan, O-GalNAc |
| Glycerol-3-phosphate acetyltransferase (Gpat4) | Encodes a de novo synthase of lysophosphatidic acid. | Endomembrane system | Unknown |
| Tumbleweed (tum) | GTPase activating protein for Rho family GTPases involved in Wnt signalling regulation | Cytoskeleton | Unknown |
| Nucleoporin 214kD (Nup214) | Part of the nuclear pore complex | Nucleus | O-GlcNAc |
| FBgn0037150 | - | - | - |
| Nucleoporin 54kD (Nup54) | Part of the nuclear pore complex | Nucleus | O-GlcNAc |

| | | | |
|---|---|---|---|
| Lingerer (lig) | Putative RNA binding protein that forms a complex with the products of orb and orb2 | Nucleus | Unknown |
| Translocase of outer membrane 5 (Tom5) | Predicted to be involved in protein targeting to mitochondrion | Mitochondrion | Unknown |

**Supplementary Table 2: List of enriched proteins from the peptide fraction of wing discs fed with $Ac_4GalNAz$.** Function was taken from FlyBase [1]; cellular location was taken from UniProt [2]; annotated glycosylation was taken from GlyGen [3].

| Gene name | Function | Cellular location | Glycosylation |
|---|---|---|---|
| Neuropeptide-like precursor 2 (Nplp2) | Involved in humoral immune response | Extracellular | Unknown |
| Ecdysone-inducible gene E2 (ImpE2) | Induces morphogenesis of imaginal discs | Extracellular | Predicted N-glycan |
| Ribosomal protein L37-1 (RpL37-1) | Binds to the 23S rRNA. | Cytosol | Unknown |
| Dumpy (dpy) | Encodes an extracellular protein involved in epidermal-cuticle attachment, apposition of wing surfaces and trachea development | Extracellular | Unknown |
| CG3777 | - | - | - |
| ATP synthase, subunit E (ATPsynE) | Predicted to enable protein transmembrane transporter activity. | Membrane | Unknown |
| Age-related lipid regulator (Arlr) | Catalyses RNA cleavage. | Endomembrane | Unknown |
| Succinate dehydrogenase, subunit C (SdhC) | Involved in mitochondrial electron transport and oxidative stress response. | Mitochondrion | Unknown |
| Narrow (nw) | Predicted to enable carbohydrate binding activity. Involved in regulation of wing disc size. | Extracellular space | Unknown |
| Ribosomal protein L36A (RpL37A) | Ribosomal protein | Cytosol | Unknown |

| | | | |
|---|---|---|---|
| Nidogen (Ndg) | Cell adhesion glycoprotein which is widely distributed in basement membranes. Involved in ECM interactions. | Extracellular | N-glycan, O-GalNAc |
| Nucleporin 62kD (Nup62) | Essential component of the nuclear pore complex | Nuclear pore | O-GlcNAc |
| Ribosomal protein L28 (RpL28) | Ribosomal protein | Cytoplasm | Unknown |
| CG2221 | - | - | - |
| Papilin (Ppn) | Encodes a secreted sulfated glycoprotein which is part of basement membranes and provisional extracellular matrices. | Extracellular | N-GlcNAc |
| CG9782 | - | - | - |
| CDGSH iron sulfur domain (Cisd2) | Predicted to enable 2 iron, 2 sulfur cluster binding activity. | Mitochondrion | Unknown |
| ER membrane protein complex subunit 7 (EMC7) | Predicted to enable carbohydrate binding activity. | Membrane | Unknown |
| Novel nucleolar protein 3 (Non3) | Required for normal assembly of the mitotic spindle. | Nucleus | Unknown |
| Ribosomal protein S25 (RpS25) | Ribosomal protein | Cytosol | Unknown |
| Ribosomal protein L26 (RpL26) | Ribosomal protein | Cytosol | Unknown |
| Bacchus (Bacc) | Tyramine-dependent nuclear regulator involved in ethanol sensitivity. | Nucleus | Unknown |
| Dystroglycan (Dg) | Major non-integrin ECM receptor that connects the ECM to the actin cytoskeleton. | Membrane, cell periphery | Unknown |
| Ribosomal protein L36 (RpL36) | Ribosomal protein | Cytosol | unknown |
| Vermiform (verm) | Chitin deacetylase-like protein that is secreted by tracheal cells and accumulates in the embryonic tracheal lumen. | Extracellular | N-GlcNAc |
| ATP synthase, subunit F (ATPsynF) | Mitochondrial membrane ATP synthase | Mitochondrion | Unknown |

| | | | |
|---|---|---|---|
| Ribosomal protein L35 (RpL35) | Ribosomal protein | Cytoplasm | Unknown |
| Ribosomal protein S9 (RpS9) | Ribosomal protein | Cytoplasm | Unknown |
| Ribosomal protein L14 (RpL14) | Ribosomal protein | Cytoplasm | Unknown |
| Sec61β subunit (Sec61β) | Encodes a protein involved in the regulation of autophagy | ER membrane | Unknown |
| Melanization protease 1 (MP1) | Encodes a serine protease which plays an essential role in the melanization immune response. | Extracellular | Predicted N-glycan |
| CG2907 | - | - | - |
| Hemomucin (Hmu) | Transmembrane mucin that may be involved in cellular adhesion and the innate immune response. | Endomembrane | O-GalNAc, predicted N-glycan |
| Eukaryotic translation initiation factor 2 subunit β (eIF2β) | Component of the eIF2 complex that functions in the early steps of protein synthesis. | Cytoplasm | Unknown |
| CG5390 | - | - | - |
| CG13220 | - | - | - |
| Ribosomal protein L18A (RpL18A) | Ribosomal protein | Cytoplasm | Unknown |
| Ribosomal protein L27A (RpL27A) | Ribosomal protein | Cytoplasm | Unknown |
| Ribosomal protein L6 (RpL6) | Ribosomal protein | Cytoplasm | Unknown |
| Serpin 27A (Spn27A) | Encodes a hemolymphatic serpin that negatively regulates a serine protease involved in the melanization cascade. | Secreted | Predicted N-glycan, O-GlcNAc |

**Supplementary Table 3: List of enriched proteins from the peptide fraction of BH-PGANT35A over WT-PGANT35A fed with $Ac_4$GalN6yne.** Function was taken from FlyBase [1]; cellular location was taken from UniProt [2]; annotated glycosylation was taken from GlyGen [3].

| Gene name | Function | Cellular location | Glycosylation |
|---|---|---|---|
| Nidogen (Ndg) | Cell adhesion glycoprotein. Involved in cell-extracellular matrix (ECM) interactions. | Secreted | N-glycan<br><br>O-GalNAc in human orthologue NID1 and NID2 |
| Tropomyosin 1 (Tm1_1) – canonical sequence | Plays a central role in the calcium dependent regulation of muscle contraction in association with the troponin complex. | Cytoplasm, cytoskeleton | Predicted O-linked sites on Thr |
| Tropomyosin 1 (Tm1_2) – isoform 1 | Plays a central role in the calcium dependent regulation of muscle contraction in association with the troponin complex. | Cytoplasm, cytoskeleton | Predicted O-linked sites on Thr |
| Serpin 55B (Spn55B) | Enables serine-type endopeptidase inhibitor activity. | Extracellular space | Unknown |
| CG13321 | - | Cytoplasm, nucleus | - |
| Odorant-binding protein 44a (Obp44a) | Predicted to enable odorant binding activity. | Secreted | Unknown |
| CG5839 | - | - | - |
| Phosphatidylethanolamine-binding protein 1 (Pebp1) | Involved in defence response to bacterium | Cytosol | Human orthologue PEBP1 N-glycan |
| CG14419 | - | - | - |
| cRAP-LALBA | Common contaminant | | |
| cRAP-CSN1S1 | Common contaminant | | |
| cRAP-CSN1S2 | Common contaminant | | |
| cRAP-ALB | Common contaminant | | |

**Supplementary Table 4: List of enriched proteins from the peptide fraction of BH-PGANT9A over WT-PGANT9A fed with $Ac_4$GalN6yne.** Function was taken from FlyBase [1]; cellular location was taken from UniProt [2]; annotated glycosylation was taken from GlyGen [3].

| Gene name | Function | Cellular location | Glycosylation? |
|---|---|---|---|
| Juvenile hormone epoxide hydrolase 3 (Jheh3) | Enables serine hydrolase activity. | ER | Unknown |
| Straw (Stw) | Predicted to enable catechol oxidase activity and tyrosinase activity. Involved in chitin-based cuticle development. | Plasma membrane | Unknown |
| Nidogen (Ndg) | Cell adhesion glycoprotein. Involved in cell-extracellular matrix (ECM) interactions. | Secreted, extracellular space | N-glycan<br><br>O-GalNAc in human orthologue NID1 and NID2 |
| CG31266 | - | Secreted | - |
| Obscurin (Unc-89, Obsc) | Encodes a titin-like protein that is needed for assembly of a symmetrical sarcomere. It is involved in myogenesis and the Hippo signalling pathway. | Cytoplasm | Unknown |
| CG3683-RA | - | - | - |
| CG42376 | - | - | - |
| Juvenile hormone epoxide hydrolase 11 (Jheh1) | Expression changes are associated with increased tolerance to oxidative stress. | ER, microsome membrane | Unknown |
| Vermiform (Verm) | Encodes a chitin deacetylase-like protein involved in cuticle development and tracheal tube size control. | Cell surface | O-GlcNAc |
| Maltase A1 (Mal-A1) | Involved in carbohydrate metabolic process | Unknown | Predicted N-glycan |

| | | | |
|---|---|---|---|
| Maltase A3 (Mal-A3) | Involved in carbohydrate metabolic process | Unknown | Predicted N-glycan |
| α Spectrin (α-Spec) | It functions in a lipoprotein pathway that delivers dietary fat to the larval fat body for storage. | Cytoplasm, cytoskeleton, Golgi apparatus | Unknown |
| Lysozyme S (LysS) | May have a function in the digestion of bacteria in the food | Extracellular space | Unknown |
| Calmodulin (Cam) | Encodes a Calcium-binding messenger protein. | Cytoplasm, cell cortex | O-GlcNAc |
| Alkaline phosphatase (Alp4) | Important role in neural and renal epithelial function. | Cell membrane | N-glycan |
| Adaptor Protein complex 1/2, β subunit (AP-1-2β) | Encodes a clathrin adaptor involved in vesicle trafficking and autophagy regulation. | Endomembrane system | Unknown |
| CG15534 | Involved in ceramide biosynthetic process and sphingomyelin catabolic process. | Secreted | Unknown |
| Maltase A5 (Mal-A5) | Involved in carbohydrate metabolic process | Unknown | Unknown |
| Tetraspanin 42Ed (Tsp42Ed) | Orthologous to human CD63. | Membrane | Unknown |
| Juvenile hormone-inducible protein 26 (JhI-26) | Encodes a sperm protein. | Unknown | Unknown |
| Ferritin 1 heavy chain homologue (Fer1HCH) | Encodes one of two subunits of the major iron storage complex, the ferritin molecule. | Golgi, secreted | Unknown |
| Karyopherin β 3 (Kary β 3) | Involved in protein import into nucleus. | Cytoplasm, nucleus | unknown |
| msb1l | Unknown | Unknown | Unknown |
| CG11882 | - | - | - |
| CG8774 | Predicted to enable metalloaminopeptidase activity. Orthologous to human ENPEP (glutamyl aminopeptidase). | Cell membrane | Unknown |

| | | | |
|---|---|---|---|
| CG31198 | Predicted to enable metalloaminopeptidase activity. Predicted to be involved in peptide catabolic process and proteolysis. | Cell membrane | Unknown |
| CG11378 | Unknown | Unknown | Unknown |
| CG5839 | Predicted to enable metalloaminopeptidase activity. Predicted to be involved in peptide catabolic process and proteolysis. Orthologous to human ERAP1 (endoplasmic reticulum aminopeptidase 1). | Cell membrane | O-GlcNAc |
| CG18585 | Predicted to enable metallocarboxypeptidase activity. Predicted to be involved in proteolysis. Orthologous to human CPA1 (carboxypeptidase A1); CPA5 (carboxypeptidase A5); and CPB1 (carboxypeptidase B1). | Secreted | Unknown |
| Tetraspanin 66E (Tsp66E) | Encodes a tetraspanin, which spans the membrane four times. | Membrane | Unknown |
| Prophenoloxidase 2 (PPO2) | Encodes a protein stored in large crystals in the crystal cells (a type of hemocyte cell) that is involved in the melanization reaction. | Secreted | Predicted N-GlcNAc |
| CG3987 | - | - | - |
| Glutathione S transferase D4 (GstD4) | Conjugation of reduced glutathione to a wide number of exogenous and endogenous hydrophobic electrophiles. | Cytoplasm | Unknown |
| Proteasome β4 subunit (Pros β 4) | Encodes a protein involved in proteasomal degradation | Cytoplasm, nucleus | Unknown |
| Lysosomal α-mannosidase II (LManII) | Involved in the degradation of asparagine-linked carbohydrates of glycoproteins | Extracellular region, lysosome | Unknown |
| CG12171 | Involved in L-fucose catabolic process. | Unknown | Unknown |
| Jonah 65Aiii (Jon65Aiii) | Involved in innate immune response and proteolysis. | Extracellular space | Unknown |

| | | | |
|---|---|---|---|
| CG8664 | - | - | - |
| CG9119 | - | - | - |
| cRAP-LALBA | Common contaminant | | |
| cRAP-CSN1S1 | Common contaminant | | |
| cRAP-ALB | Common contaminant | | |

### Methods

#### Generation of Ubi-MOE and 10xUAS-MOE plasmids

The pJFRC81 vector (Addgene #36432) is called 10xUAS vector from now on, and was used for tissue- or cell-specific expression of the genes for NahK and AGX1$^{F383A}$ using the GAL4/UAS system. The pUbi-p63E-Casper vector [4] is called Ubi vector from here on. Both Ubi and 10xUAS vectors were linearised using NotI-HF and XbaI restriction enzymes with rCutSmart™ Buffer according to the manufacturer's specifications with incubation for 30 min at 37 °C. Amplified linearised products were purified using the NucleoSpin® Gel and PCR Clean-up kit. The pSBbi plasmid co-expressing AGX1$^{F383A}$ (FLAG-tagged) and NahK (HA-tagged) on the pSBbi plasmid (originally a gift from Eric Kowarz (Addgene #60514)) was previously made by us [5]. This plasmid was used as a template to amplify the genes for AGX1-FLAG and NahK-HA using CloneAmp HiFi PCR Premix according to the manufacturer's specifications with the primers Ubi_fwd, Ubi_rev and 10xUAS_fwd, 10xUAS_rev. Pre-infusion PCR was performed with 1% (v/v) DMSO. Initial denaturation at 98 °C for 1 min was followed by 25 cycles with the following conditions: 98 °C (10 s), 55-68 C (30 s), 72 °C (3 min). A final extension at 72 °C for 15 min was followed by a hold at 10 °C. Resulting pre-infusion products were purified using the NucleoSpin® Gel and PCR Clean-up kit. In-fusion cloning of the AGX1$^{F383A}$ and NahK into the Ubi and 10xUAS vectors was performed using 5X In-fusion HD enzyme premix according to the manufacturer's specifications with an incubation at 50 °C for 15 min. The resulting in-fusion products (5 µL) were transformed by mixing with Stellar™ Competent Cells (50 µL) and placing on ice for 30 min. Cells were heat shocked for 30 s at 42 °C and then placed on ice for 5 min. SOC medium was added to bring the total volume to 950 µL and cells were incubated by shaking at 225 rpm for 1 h at 37 °C. Colonies were inoculated into 100 mL cultures containing Ampicillin (100 µg/mL) overnight by shaking (225 rpm, 37 °C). Plasmids were purified using PureLink™ HiPure Plasmid Filter Maxiprep kit and sequenced again using Genewiz Sanger sequencing and Full Circle nanopore sequencing.

#### Generating plasmids for …

Plasmids encoding WT-PGANT9A and 35A in the backbone pUAST were previously made [6,7]. Genes for the corresponding BH-PGANTs in pUCIDT-Amp GolgenGate were synthesised by IDT. The 2-Ubi-INS vector, 2 Ubi from here on, was made by us.

The 2 Ubi vector was linearised by restriction enzyme digestion using BamHI and AvrII with rCutSmart$^{TM}$ Buffer according to the manufacturer's specifications with incubation for 15 min at 37 °C. Amplified linearised products were purified using the NucleoSpin® Gel and PCR Clean-up kit. AGX1$^{F383A}$ and NahK, were extracted from the Ubi-MOE plasmid described above using CloneAmp HiFi PCR Premix according to the manufacturer's specifications and the following primers: 2Ubi_pre-inf_fwd and 2Ubi_pre-inf_rev. Pre-infusion PCR was performed with 1% (v/v) DMSO in the reaction mixtures due to the high annealing temperatures of the in-fusion primers. Initial denaturation at 98 °C for 1 min was followed by 25 cycles with the following conditions: 98 °C (10 s), 55-68 C (30 s), 72 °C (3 min). A final extension at 72 °C for 15 min was followed by a hold at 10 °C. Resulting pre-infusion products were purified using the NucleoSpin® Gel and PCR Clean-up kit. In-fusion cloning of the AGX1$^{F383A}$ and NahK into the 2 Ubi vector was performed using 5X In-fusion HD enzyme premix according to the manufacturer's specifications with an incubation at 50 °C for 15 min. The resulting in-fusion products (5 µL) were transformed with Stellar$^{TM}$ Competent Cells (50 µL) (see above). Plasmids were purified using QIAprep Spin Miniprep Kit and sequenced. Ubi-MOE linearised using restriction enzyme digestion using NotI-HF and XbaI with rCutSmart$^{TM}$ Buffer according to the manufacturer's specifications with incubation for 45 min at 37 °C. WT- and BH-PGANT sequences were extracted by restriction enzyme digestion from the following previously generated plasmids: UAS-WT-PGANT-9A, UAS-BH-PGANT-9A, UAS-WT-PGANT-35A, UAS-BH-PGANT-35A. Restriction enzyme digestion was performed using NotI-HF and XbaI with rCutSmart$^{TM}$ Buffer according to the manufacturer's specifications with incubation for 45 min at 37 °C. Amplified linearised products were purified using the NucleoSpin® Gel and PCR Clean-up kit. WT- and BH-PGANTs were inserted into 2 Ubi-MOE by ligation using T4 DNA Ligase buffer and T4 DNA ligase according to the manufacturer's specifications with incubation for 1 h at RT. Ligation products (5 µL) were transformed with Stellar$^{TM}$ Competent Cells (50 µL) (see above). Colonies were inoculated into 100 mL cultures containing Ampicillin (100 µg/mL) overnight by shaking (225 rpm, 37 °C). Plasmids were purified using PureLink$^{TM}$ HiPure Plasmid Filter Maxiprep kit. Sequences were verified by sequencing.

**Generation of plasmids for K-562 cell transfection**

The pSBbi-AGX1$^{F383A}$-NahK plasmid from our previous study[8] was linearised by restriction enzyme digestion using SfI and rCutSmart® Buffer according to the manufacturer's specifications with incubation at 50 °C for 15 min. Amplified linearised products were purified using the NucleoSpin® Gel and PCR Clean-up kit. WT and BH PGANTs were extracted from the 2 Ubi plasmid (see above) using CloneAmp HiFi PCR Premix according to the manufacturer's specifications and the following primers: PGANT3_fwd, PGANT9_fwd, PGANT35A_fwd and PGANTs_rev. Pre-infusion PCR was performed with 1% (v/v) DMSO in the reaction mixtures due to the high annealing temperatures of the in-fusion primers. Initial denaturation at 98 °C for 1 min was followed by 25 cycles with the following conditions: 98 °C (10 s), 55-68 C (30 s), 72 °C (90 s). A final extension at 72 °C for 15 min was followed by a hold at 10 °C. Resulting pre-infusion products were purified using the NucleoSpin® Gel and PCR Clean-up kit. In-fusion cloning was performed using 5X In-fusion HD Enzyme Premix according to the manufacturer's specifications with an incubation at 50 °C for 15 min. In-fusion products were transformed using Stellar™ Competent Cells (see above). Plasmids were purified by miniprep then maxiprep (see above). Sequences were verified by sequencing.

**K-562 cell transfection**

K-562 cells (ATCC CCL-243) were grown in RPMI, 10 % Foetal Bovine Serum (FBS), penicillin (100 U/mL) and streptomycin (100 mg/uL). The following pSBbi plasmids co-expressing AGX1$^{F383A}$ and NahK were used in the stable transfection: WT-PGANT3, BH-PGANT3, WT-PGANT9A, BH-PGANT9A, WT-PGANT9B, BH-PGANT9B, WT-PGANT35A and BH-PGANT35A. Cells were transfected using Lipofectamine LTX according to the manufacturer's instructions with 2.4 µg pSBbi plasmid, 125 ng pCMV(CAT)T7-SB100 per well in a 6-well plate at 6.25x$10^6$ cells/mL. Cells were incubated overnight at 37°C after transfection and then selected with hygromycin B (100 µg/mL) for 14 days with splitting every 2-3 days.

**Cell surface labelling of K-562 cells**

Transfected K-562 cells were plated in a 6-well plate at 4x$10^5$ cells/mL in 1.6 mL growth medium and fed with either DMSO, $Ac_4$GalN6yne (2 µM) or $Ac_4$ManNAlk (10 µM) from a 10X stock in growth medium. Cells incubated overnight at 37 °C and were then pelleted at 500 g for 5 min at 4 °C. Pellets were resuspended in ice-cold 2 % FBS in PBS (100 µL) and transferred to a 96-well plate before centrifuging (500 g, 3 min, 4 °C). Pellets were washed with ice-cold 2 % (v/v) FBS in PBS (2x 200 µL) and the supernatant was discarded. Pellets were resuspended in 2 % FBS in PBS (35 µL) and 2X CuAAC solution (35 µL) made up of $CuSO_4{\cdot}5H_2O$ (200 µM), BTTAA (1 mM), sodium ascorbate (10 mM), aminoguanidine hydrochloride (10 mM) and CF680-picolyl azide (200 µM). Cells were incubated for 7 min on an orbital shaker. The click reaction was quenched with Bathocuproine disulfonate (BCS) (3 mM, 35 µL) and cells were pelleted at 500 g for 3 min at 4 °C. Pellets were washed with PBS (2x 200 µL) and then lysed with Lysis Buffer A (100 µL) containing Tris-HCl (pH 8, 50 mM), NaCl (150 mM), 1 % Triton-X-100, 0.5 % sodium deoxycholate, 0.1 % sodium dodecyl sulfate (SDS), $MgCl_2$ (1 mM), Benzonase® nuclease (100 mU/µL), 1X Halt$^{TM}$ Protease Inhibitor and $MgCl_2$ (1 mM). for 20 min on an orbital shaker at 4 °C. Lysates were centrifuged (1500 g, 20 min, 4 °C) and supernatant was collected for protein concentration analysis by Pierce$^{TM}$ BCA Protein Assay Kit. Where indicated, lysates (20 µg) were treated with PNGase F (5 U) in PBS and incubated overnight at 37 °C. The reaction was quenched by heating to 95 °C for 10 s with subsequent cooling to 4 °C. Lysates were mixed with 4X Loading Buffer and run on an SDS-PAGE gel for 1 h at 160 V. The gel was imaged for in-gel fluorescence and proteins were then transferred to a nitrocellulose membrane Bio-Rad Trans-Blot Turbo system at High MW setting (2.5 A, 10 min). Membrane was incubated with Revert$^{TM}$ 700 Total Protein Stain (LI-COR Bioscience, Lincoln, USA) (10 mL) and then washed twice with Wash Solution containing 6.7% (v/v) glacial acetic acid, 30% (v/v) methanol in water (2x 10 mL). Membrane was destained using Destaining Solution containing sodium hydroxide (0.1 mM) and 30% (v/v) methanol in water (10 mL) for 5 min and then blocked using Intercept® (TBS) Protein-Free Blocking Buffer (10 mL) (LI-COR) for 1 h at RT. The following antibodies were used: Rabbit anti-VSV-G (PA1-29903, 1:800 in Intercept Antibody diluent, overnight incubation at 4 °C), Mouse anti-HA (ab18181, 1:800 in Intercept Antibody diluent, overnight incubation at 4 °C), Rabbit anti-FLAG (PA1-984B, 1:1000 in Intercept Antibody diluent, overnight incubation at 4 °C).

**Microinjection of *Drosophila melanogaster* embryos for generation of transgenic flies**

*Drosophila* embryos with attP2 docking site were rinsed with water and then dechorinated using 50 % household bleach. Embryos were rinsed with water twice and then lined up on a cover slip with heptane glue and dehydrated for 7-9 min. Embryos were covered with Voltalef oil (VWR Chemicals, Radnor, USA) for the injection using the Eppendorf FemtoJet 5247 micromanipulator system (Eppendorf, Hamburg, Germany). Candidates were confirmed by red eyes (mini-white gene) and then balanced with TM3/TM6b flies to generate a stock of transgenic flies.

**Injection of embryos with modified sugars**

Embryos ($w^{1118}$ or Ubi-MOE) were washed from grape juice plates with water and transferred to a mesh basket. Embryos were rinsed with water to remove any remaining yeast. Embryos were dechorinated in 50 % bleach for 2 min and then washed with water to remove bleach. Embryos were lined up on a cover slip with heptane glue and dehydrated for 7-9 min. Embryos were covered with Voltalef oil for the injection. Embryos were injected with $Ac_4$GalNAz (5 mM, 5 % in water), $Ac_4$GalNAlk (15 mM, 15 % in water) or DMSO (5 % or 15 % in water) within 2 hours of egg laying and left to develop into first instar larvae for 24 h before collection.

**Modified sugar feeding**

Fly medium containing different peracetylated sugars was prepared by heating wheat food in the microwave until liquid. Liquid fly medium (10 mL) was mixed with the $Ac_4$GalNAz (100, 300 or 500 µM) or DMSO. Adult transgenic flies were placed on the food containing either the $Ac_4$GalNAz or DMSO and left to lay eggs overnight. Adults were removed and embryos left to develop at 25 °C until collection time. Third instar larvae were collected using the protocol below.

**Larval tissue collection and lysis**

Third instar wandering larvae were dissected in PBS. Tissues were transferred to Lysis Buffer A. Tissues were first crushed by hand using a pestle and then sonicated using a Bioruptor® Plus (Diagenode, Seraing, Belgium) at 4 °C (LOW, 30 s ON, 30 s OFF, 20 cycles for whole larvae, 15 cycles for other tissues). Lysates were centrifuged (20 min, 14000 g, 4 °C) and the supernatant was transferred to a new tube and centrifuged again (15 min, 14000 g, 4 °C). Concentrations were determined using the Pierce™ Rapid Gold BCA Protein Assay Kit (ThermoFischer Scientific, Waltham, USA).

**Embryo feeding and collection**

Adult flies were placed in cages containing $Ac_4$GalNAz (300 or 500 µM) or DMSO on a 6 cm grape juice plate and kept at 25 °C. Plates were changed every day and embryos were collected from day 3 at 22 h after egg laying. Embryos were detached from plates using water and transferred to a mesh basket. Embryos were rinsed with water (approx. 500 mL) and soaked in 50% household bleach for 2 min. Embryos were washed with water until the bleach smell was no longer detected (approx. 500 mL). Embryos were transferred to Lysis Buffer (approximately 1 mL Lysis Buffer per plate). Samples were sonicated using the Bioruptor® at 4 °C (LOW, 30 s ON, 30 s OFF, 15 cycles) and then centrifuged (10 min, 10000 g, 4 °C). The supernatant was centrifuged again (5 min, 10000 g, 4 °C). Concentrations were determined using the Pierce™ Rapid Gold BCA Protein Assay Kit.

**Labelling of lysates and analysis by streptavidin blot using CuAAC**

Lysates (30-60 µg) were incubated with 10X CuAAC solution containing $CuSO_4 \cdot 5H_2O$ (6 mM), BTTAA (12 mM), sodium ascorbate (100 mM), aminoguanidine hydrochloride (100 mM) and biotin-alkyne (1 mM) for 3 h at RT. When CuAAC was performed with biotin-picolyl-azide, the following 10X CuAAC was used: $CuSO_4 \cdot 5H_2O$ (3 mM), BTTAA (6 mM), sodium ascorbate (50 mM), aminoguanidine hydrochloride (50 mM) and biotin-picolyl-azide (1 mM). Clicked samples (15-30 µg) were mixed with 4X Loading Buffer and boiled for 3 min at 95 °C before running on an SDS-PAGE for 1 h at 160 V and Western blot. Membrane was incubated with Revert™ 700 Total Protein Stain (LI-COR Bioscience, Lincoln, USA) (10 mL) and then washed twice with Wash Solution containing 6.7% (v/v) glacial acetic acid, 30% (v/v) methanol in water (2x 10 mL). Membrane was destained using Destaining Solution containing sodium hydroxide (0.1 mM) and 30% (v/v) methanol in water (10 mL) for 5 min and then blocked using Intercept® (TBS) Protein-Free Blocking Buffer (10 mL) for 1 h at RT. Labelling was analysed using IRDye 800CW Streptavidin according to the manufacturer's specifications. The following primary antibodies were used: Mouse anti-HA (ab18181, (abcam, Cambridge, UK) 1:800 in Intercept Antibody diluent, overnight incubation at 4 °C), Rabbit anti-FLAG (PA1-984B (ThermoFischer Scientific, Waltham USA), 1:1000 in Intercept Antibody diluent, overnight incubation at 4 °C). The following secondary antibodies were used: IRDye® 800CW goat anti-rabbit (LI-COR) and IRDye® 800CW goat anti-mouse (LI-COR). All imaging was performed using the Odyssey CLx, images were processed on ImageStudioLite software and cropped using Adobe Illustrator.

**Enrichment of clicked lysates**

Lysates were enriched on neutravidin beads before running the SDS-PAGE. Amicon Ultra-0.5 (3 kDa) devices were washed with PBS (2x 300 µL) by centrifugation (14000 g, 10 min). Clicked lysates (300 µL) were added to the Amicon device and volume was filled to 500 µL for all samples before centrifugation (14000 g, 15 min at 4°C). The flow-through was discarded and PBS (300 µL) was added to each sample followed by another centrifugation (14000 g, 15 min at 4°C). Following sample recovery, sample volumes were adjusted to 300 µL with PBS and added to Neutravidin bead slurry (300 µL), previously washed with PBS (3x 300 µL). Samples were incubated with the beads on a rotator for 1 h at RT. Neutravidin beads were washed with 0.1 % SDS, 1 % Triton-X-100 in PBS (2x 300 µL), then with 6 M urea in PBS (2x 300 µL) and finally with PBS (2x 300 µL). Beads were then resuspended in a solution (40 µL) containing a 2:2:1 (v/v/v) mixture of Tris-Cl (pH 6.5), 80 % (v/v) glycerol in water and 20 % (w/v) SDS and boiled for 10 min at 95°C. Supernatants were collected and run on an SDS-PAGE for analysis by streptavidin blot as described above.

**Sample preparation for MS-glycoproteome analysis of larval tissues and embryos**

Wing imaginal discs (300 µg) from third instar wandering larvae and embryos (400 µg) fed with 300 µM $Ac_4GalNAz$ or DMSO were collected and lysed as described above. To remove endogenously biotinylated proteins, lysates were incubated for 30 min with Neutravidin beads slurry (300 µL), previously washed with PBS (3x 300 µL). For samples fed with $Ac_4GalNAz$: the supernatant was collected and treated with 10X CuAAC solution containing $CuSO_4{\cdot}5H_2O$ (6 mM), BTTAA (12 mM), sodium ascorbate (100 mM), aminoguanidine hydrochloride (100 mM) and Biotin-DADPS-alkyne (1 mM, BroadPharm, San Diego, USA) to a 1x solution and incubated for 3 h at RT. From here on, LC-MS grade reagents were used. Methanol/chloroform (1:0.25) precipitation was performed by mixing ice-cold methanol and chloroform with the lysates and centrifuging (18000 g, 10 min, 4 °C). Pellets were air-dried and resuspended in 0.1 % (v/v) SDS in PBS (250 µL) and sonicated for 25 min before centrifuging (3700 g, 5 min). Pellets were treated with 6 M urea in PBS (250 µL) and sonicated for 25 min and then centrifuged (3700 g, 5 min). Pellets were treated with PBS (250 µL) and sonicated for 25 min and then centrifuged (3700 g, 5 min). SDS, urea and PBS supernatants were combined and incubated for 1 h at RT with dimethylated Neutravidin beads slurry (300 µL), previously washed with PBS (3x 300 µL). Supernatants were discarded and beads were washed with 1 % (v/v) SDS in water (3x 350 µL), 6 M urea in PBS (6x 350 µL) and 50 mM ammonium bicarbonate (Ambic) in water (6x 350 µL). Beads were resuspended in 10 mM DTT in Ambic (100 µL) and incubated at 50 °C for 15 min. Beads were washed with Ambic (2x 100 µL) and then incubated with 20 mM iodoacetamide in Ambic (100 µL) for 30 min in the dark. The reaction was quenched with 10 mM DTT in Ambic (100 µL) and the supernatant was discarded. Beads were washed with 50 mM Ambic (3x 100 µL) and then resuspended in 50 mM Ambic (100 µL) before incubating with LysC (Promega, Madison, USA) (300 ng) overnight at 37 °C. The supernatant was transferred to a new tube and incubated with trypsin (300 ng) for 8 h at 37 °C before quenching with 1 % (v/v) formic acid. This forms the peptide fraction. Beads were incubated with 1 % (v/v) formic acid (150 µL) for 30 min at RT and the supernatant was collected. This process was repeated and the supernatants were combined. Beads were washed with acetonitrile (100 µL) and the supernatant was combined with previous supernatants. Solvents were removed by Speedvac (organic, 37 °C) and pellets were resuspended in 50 mM Ambic (100 µL). Samples were incubated with trypsin (300 ng) for 8 h at 37 °C before quenching with 1 % (v/v) formic acid. This forms the glycopeptide fraction. Glycopeptide and peptide fractions were desalted using the UltraMicroSpin™ columns (The Nest group, Inc., Ipswich USA) according to manufacturer's specifications and remaining solvent was removed by SpeedVac. Samples were reconstituted in 0.1 % (v/v) formic acid (16 µL for glycopeptide, 20 µL for peptide), sonicated for 25 min and then centrifuged (18000 g, 10 min).

Peptide samples were injected into an Orbitrap Eclipse™ Tribid™ Mass Spectrometer coupled to a Dionex UltiMate™ 3000 RSLCnano System. Peptides were separated on an EASY-spray PepMap rapid separation liquid chromatography C18 column (75 µm × 500 mm, 2 µm particle size; Thermo Scientific) using a gradient of 2–95% solvent B (5% DMSO, 0.1% formic acid, 75% acetonitrile and 20% water) over 120 min at 275 nL $min^{-1}$.

Full-scan MS1 spectra acquired in the Orbitrap were collected at a resolution of 120,000 at full width at half-maximum and a mass range from 300 to 1,500 *m*/*z* with a Standard AGC target and Auto maximum injection time. Dynamic exclusion was enabled with a repeat count of 1 and an exclusion duration of 20 s. Only charge states 2-6 with an intensity threshold greater than $1x10^4$ were selected for fragmentation. MS2 scans were generated at top speed for 3 s. HCD was performed on all selected precursor masses with the following parameters: 30 % normalised collision energy with a 1.2 *m/z* isolation window (Quadrupole), Standard AGC target and Dynamic maximum injection time. Each sample was injected in triplicate. Wing disc samples are two independent replicates with three technical replicates each for DMSO-fed and $Ac_4$GalNAz-fed samples. Embryo samples are one replicate for DMSO-fed with three technical replicates and three independent replicates for $Ac_4$galNAz-fed with three technical replicates each.

**MS data acquisition for glycopeptide analysis**

Samples were analyzed by online nanoflow LC–tandem MS using an Orbitrap Eclipse Tribrid MS instrument (Thermo Fisher Scientific) coupled to a Dionex UltiMate 3000 RSLCnano HPLC (Thermo Fisher Scientific). Glycopeptides were separated on an EASY-spray PepMap rapid separation liquid chromatography C18 column (75 µm × 500 mm, 2 µm particle size; Thermo Scientific) using a gradient of 2–95% solvent B (5% DMSO, 0.1% formic acid, 75% acetonitrile and 20% water) over 140 min at 275 nL $min^{-1}$.

Full-scan MS1 spectra acquired in the Orbitrap were collected at a resolution of 120,000 at full width at half-maximum and a mass range from 300 to 1,500 *m*/*z*.

Dynamic exclusion was enabled with a repeat count of three, repeat duration of 10 s, and exclusion duration of 10 s. Only charge states 2-6 were selected for fragmentation.

MS2 scans were generated at top speed for 3 s. HCD was performed on all selected precursor masses with the following parameters: isolation window of 2 *m/z*, 28% normalized collision energy, orbitrap detection (resolution of 30,000), maximum injection time of 54 s, and a 100 % normalized AGC target. An additional ETD fragmentation of the same precursor was triggered if (1) the precursor mass was between 300 and 1000 *m/z* and (2) fingerprint ions generated by the specific tags (329.1455 and 311.1348) were present at ±20 ppm and greater than 10 % relative intensity.

**MS-Glycoproteomics of whole larvae expressing BH PGANTs**

Third instar wandering larvae expressing NahK, AGX1$^{F383A}$ and WT- or BH-PGANT35A or 9A (350 µg) fed with 300 µM $Ac_4$GalN6yne were collected (whole larvae without fat body) and lysed as described above. To remove endogenously biotinylated proteins, lysates were incubated for 30 min with neutravidin beads slurry (300 µL), previously washed with PBS (3x 300 µL). The supernatant was collected and treated with PNGase F (5 U) in PBS with overnight incubation at 37 °C. The reaction was quenched by heating to 95 °C for 10 s with subsequent cooling to 4 °C. The solutions were incubated with 10X CuAAC solution containing $CuSO_4 \cdot 5H_2O$ (6 mM), BTTAA (12 mM), sodium ascorbate (100 mM), aminoguanidine hydrochloride (100 mM) and Biotin-DADPS-picolyl azide (1 mM) for 3 h at RT. From here on, LC-MS grade reagents were used. Treated lysates were added to Amicon Ultra-0.5 (10 kDa) device, previously washed with PBS (3x 300 µL), to remove excess biotin-DADPS-picolyl azide. Amicon devices were centrifuged (14000 g, 15 min at 4°C) and flow through was discarded. Buffer exchange was performed by addition of PBS (3x 400 µL) and centrifugation (14000 g, 15 min at 4°C). To recover the proteins, Amicon devices were placed upside down into a new microcentrifuge tube and centrifuged (1000 g, 2 min). Sample volumes were adjusted to 300 µL with PBS and added to dimethylated Neutravidin bead slurry (300 µL), previously washed with PBS (3x 300 µL). Samples were incubated with the beads on a rotator for 1 h at RT. Beads were washed with 1 % (v/v) SDS in water (2x 350 µL), 6 M urea in PBS (2x 350 µL) and 50 mM Ambic in water (2x 350 µL). Beads were resuspended in 10 mM DTT in Ambic (100 µL) and incubated at 50 °C for 15 min. Beads were washed with Ambic (2x 100 µL) and then incubated with 20 mM iodoacetamide in Ambic (100 µL) for 30 min in the dark. The reaction was quenched with 10 mM DTT in Ambic (100 µL) and the supernatant was discarded. Beads were washed with 50 mM Ambic (2x 100 µL) and then resuspended in 50 mM Ambic (100 µL) before incubating with LysC (300 ng) overnight at 37 °C. The supernatant was transferred to a new tube and incubated with trypsin (300 ng) for 8 h at 37 °C before quenching with 1 % (v/v) formic acid. This forms the peptide fraction. Beads were incubated with 1 % (v/v) formic acid (150 µL) for 30 min at RT and the supernatant was collected. This process was repeated and the supernatants were combined. Solvents were removed by SpeedVac (organic, 37 °C) and pellets were resuspended in 50 mM Ambic (100 µL). Samples were incubated with trypsin (300 ng) for 8 h at 37 °C before quenching with 1 % (v/v) formic acid. This forms the glycopeptide fraction. Glyco- and peptide fractions were desalted using EvoTips according to manufacturer's specifications and remaining solvent was removed by SpeedVac. Samples were reconstituted in 0.1 % (v/v) formic acid (16 µL for glycopeptide, 20 µL for peptide).

Analysis was performed on an Evosep One coupled online to a hybrid (trapped ion mobility spectrometry) TIMS quadrupole TOF (time of flight) mass spectrometer (Bruker TimsTOF Pro 2, Bruker, Billerica, United States) via a captive spray nano-electrospray ion source. Peptides were separated on an EvoSep 60 SPD performance column (80 cm x 150 µm x 1.5 µm, Bruker). Samples were analysed with an ion mobility range from $1/K_0$ = 0.60 to 1.60 $V.s.cm^{-2}$. Equal ion accumulation time and ramp times were applied in the dual TIMS analyser of 100 ms each. Mass spectra were recorded from 100–1700 *m/z*. The ion mobility dimension was calibrated regularly using the Bruker ES-TOF tuning mix (*m/z*, $1/K_0$: 622.0280, 0.9915 $V.s.cm^{-2}$; 922.0098, 1.1986 $V.s.cm^{-2}$; and 1221.9906, 1.3934 $V.s.cm^{-2}$). When operating the mass spectrometer in diaPASEF mode, isolation windows covered a mass range of 282.2 – 1199.6 *m/z* with a mobility range of 0.60 to 1.60 $V.s.cm^{-2}$ and a cycle time of 1.37 s. Collision energy was ramped linearly as a function of ion mobility from 20 eV at $1/K_0$ = 0.60 $V.s.cm^{-2}$ to 59 eV at $1/K_0$ = 1.60 $V.s.cm^{-2}$. Reconstituted samples were run as three biological replicates with three technical replicates each except for replicate 1 of WT-PGANT9A run with two technical replicates and replicate 3 of BH-PGANT9A run with two technical replicates.

**Analysis of glycopeptides using Byonic™**

The protocol used for this section is based on the STAR Protocols by Calle *et al*., 2023 [9]. A focused parameter FASTA search was performed against the *Drosophila melanogaster* species FASTA obtained from UniProt. Parameters for the search in Byonic™ (v5.8.24) were as follows: 10 ppm mass tolerance for precursor mass ions with 20 ppm fragment mass tolerance for HCD fragmentation and 0.1 Da mass tolerance for ETD fragmentation; up to two missed cleavages per peptide with semi-specific, C-terminal tryptic digestion (R, K cleavage sites); 1 % false discovery rate (FDR) with standard reverse-decoy techniques. Carbamidomethyl was set as a fixed modification and Deamidated and Oxidation were set as variable modifications.. Under the 'Advanced' tab, 'Create a focused database' was ticked to generate focused FASTA files for individual raw data files.

The focused FASTA file for each sample was used to search for glycan modifications using the following parameters in the 'Glycans' tab:
HexNAc(1)Hex(1)NeuAc(2) 125.059 @ OGlycan | common2
HexNAc(1) 125.059 @ OGlycan | common2
HexNAc(1)Hex(1) 125.059 @ OGlycan | common2
HexNAc(1)NeuAc(1) 125.059 @ OGlycan | common2
HexNAc(1)Hex(1)NeuAc(1) 125.059 @ OGlycan | common2
HexNAc(2)Hex(1)NeuAc(1) 125.059 @ OGlycan | common2
HexNAc(2)Hex(2) 125.059 @ OGlycan | common2
HexNAc(2)Hex(2)NeuAc(1) 125.059 @ OGlycan | common2

The excel files from the output of Byonic™ were used to filter peptides containing glycan modifications and specific cleavage. If any hits satisfied this criteria, MS-spectra were manually evaluated in ExCalibur.

**Analysis using Dia-NN and Perseus**

Dia-NN (1.9.1) was used to process MS files from TimsTOF. A spectral library using *Drosophila melanogaster* FASTA file from UniProt was generated for consistency and to reduce subsequent search time. The following parameters were used: 'FASTA digest for library-free search/library generation' and 'Deep learning-based spectra, RTs and IMs prediction' were ticked; Trypsin/P cleavage with 1 missed cleavage was set; oxidation, N-terminal acetylation and carbamidomethylation were selected. Raw data MS files and the spectral library generated were uploaded to Dia-NN. The following parameters were used for Dia-NN search: Trypsin/P cleavage with 1 missed cleavage; oxidation, N-terminal acetylation and carbamidomethylation were selected; 1 % FDR; 15 ppm mass accuracy; 15 ppm MS1 accuracy; peptidoforms, match between runs (MBR), no shared spectra and heuristic protein interference were selected; for protein inference, 'Genes' was selected; for neural network classifier, 'Single-pass mode' was selected; for quantification strategy, 'QuantUMS (high accuracy)' was selected; for cross-run normalisation, 'RT-dependent' was selected; for library generation, 'IDs, RT &IM profiling' was selected; for speed and RAM usage, 'Optimal results' was selected.

Perseus (2.0.11.0) was used to analyse the pg_matrix.tsv and pr_matrix.tsv files in the following order: columns were annotated with protein name, gene name, UniProt identifier and FlyBase identifier. Rows were annotated by categorical annotation to rename replicates with the same name for subsequent statistical analysis. Data was transformed to a logarithmic base using log2(x). Rows were filtered based on valid values (70 % cut-off) and missing values were replaced. Data was normalised using column-wise median subtraction. Two-sided two-sample Welch's t-test with Benjamini-Hochberg FDR-adjusted p-value was performed. Statistically enriched proteins were chosen with a cut-off of p-value 0.05 and 2-fold enrichment.

**Analysis of signal peptides from proteomics**

UniProt ID mapping tool was used to download fasta files of protein hits for each experiment. Fasta files were uploaded to SignalP 6.0 (DTU Health Tech) [10] to predict signal peptides using eukarya with default parameters. Proteins classified as 'SP' were considered to have a classical signal peptide.

**Lectin blots**

The following strains of third instar wandering larvae were collected: $w^{1118}$; Ubi AGX1$^{F383A}$, NahK; Ubi AGX1$^{F383A}$, NahK, Ubi WT PGANT 35A; Ubi AGX1$^{F383A}$, NahK, Ubi BH PGANT 35A; Ubi AGX1$^{F383A}$, NahK, Ubi WT PGANT 9A; Ubi AGX1$^{F383A}$, NahK, Ubi BH PGANT 9A. The fat body was removed by dissection and remaining larval tissues were lysed as described above. Samples (30 μg) were mixed with 4X Loading

Buffer and boiled for 3 min at 95 °C before running on an SDS-PAGE for 1 h at 160 V. Proteins were transferred to a nitrocellulose membrane. Membrane was stained for total protein as described above and then blocked using Intercept® (TBS) Blocking Buffer (10 mL) for 30 min at RT. Membrane was incubated with avidin block solution (10 mL, abcam kit 64212) for 15 min at RT and then washed with TBS (3x 10 mL) before incubation with biotin block solution (10 mL, abcam kit 64212) for 15 min at RT followed by washing with TBS (3x10 mL) to remove background from endogenously biotinylated proteins. Membrane was incubated for 1 h at RT with the following biotinylated lectins from vector laboratories (Newark, USA): ConA (40 µg/mL in PBST, 0.1 mM $CaCl_2$, 0.1 mM Mn $Cl_2$), SBA (20 µg/mL in PBST, 0.1 mM $CaCl_2$, 0.1 mM Mn $Cl_2$), DBA (20 µg/mL in PBST, 0.1 mM $CaCl_2$,), VVL (20 µg/mL in PBST, 0.1 mM $CaCl_2$), MALII (20 µg/mL in PBST), AAL (20 µg/mL in PBST), SNA (20 µg/mL in PBST). Membranes were washed with PBS (3x 10 mL) and PBST (3x 10 mL) before incubation with IRDye 800CW Streptavidin (LI-COR) according to the manufacturer's specifications.

**HPA blotting to check Ndg labelling by PGANT9A, 9B and 35A in S2R+ cells**

Nidogen cDNA (FBgn0026403) was subcloned from pUC57-Nidogen (Genscript) into the HindIII and NotI sites of pIB-V5/His vector (ThermoFisher Scientific), fused with V5/His tag. The plasmid pIB-nidogen-V5/His was co-transfected with pIB-pgant9a-V5/His (or pIB-pgant9a-FLAG), pIB-pgant9b-V5/His (or pIB-pgant9b-FLAG), pIB-pgant35a-V5/His and pIB-V5/His into S2R+ cells using Effectene transfection reagent (QIagen) according to manufacturer's instructions. After 72hrs, the cells and media were collected. The expressed proteins were purified using anti-V5-agarose (ThermoFisher Scientific) from cell lysates and media according to manufacturer's instructions. Purified proteins or cell lysates were electrophoresed in 4-12% SDS-PAGE gels and then the proteins were transferred to nitrocellulose membranes. After blocking with blocking buffer (LI-COR), the membranes were incubated with anti-V5 antibody (ThermoFisher Scientifc) and IRDye 680LT-conjugated HPA. The membranes were then washed and incubated with 800CW-conjugated anti-mouse IgG (LI-COR). Finally, the membranes were washed with PBST (0.1% Tween-20), rinsed in PBS, and imaged using a Li-COR Odyssey Infrared Imaging System.

**Primer and plasmid sequences**

| Primer name | Sequence |
|---|---|
| Ubi_fwd | CCGGGTACCgcggccgcGCCACCatgaacattaatgacctcaaactcacgttg |
| Ubi_rev | cgccaggccttctagactaGGCGTAGTCGGGCACGTCGTA |
| 10xUAS_fwd | TTACTTCAGGCGGCCGCGCCACCatgaacattaatgacctcaaactcacgttg |
| 10xUAS_rev | AACGATTCATTCTAGActaGGCGTAGTCGGGCACGTCG |
| 2Ubi_pre-inf_fwd | ccacgGGTCACCactGGATCCAACTTAAAAAAAAAAAATCAAAatgaacattaatgacctcaaactcacgttg |
| 2Ubi_pre-inf_rev | ttaaaaacgattcatCCTAGGCTAGGCGTAGTCGGGCACGtcgtagg |
| PGANT3_fwd | aaactaccccaagctggcctctgaggccAACTTAAAAAAAAAAAATCAAAatgggtctccgg |
| PGANT9_fwd | aaactaccccaagctggcctctgaggccAACTTAAAAAAAAAAAATCAAAatggccttcatctgg |
| PGANT35A_fwd | aaactaccccaagctggcctctgaggccAACTTAAAAAAAAAAAATCAAAatgatgcaaatcaagcg |
| PGANTs_rev | ttgatccccaagcttggcctgacaggccCTACTTACCCAGGCGGTTCATT |
| 2-Ubi-INS | GACTCAGGTGGCACTTTTCGGGGAAATGTGCGCGGAACCCCTATTTGTTTATTTTTCTAAATACATTCAAATATGTATCCGCTCATGAGACAATAACCCT<br>GATAAATGCTTCAATAATATTGAAAAAGGAAGAGTATGAGTATTCAACATTTCCGTGTCGCCCTTATTCCCTTTTTTGCGGCATTTTGCCTTCCTGTTTTTG<br>CTCACCCAGAAACGCTGGTGAAAGTAAAAGATGCTGAAGATCAGTTGGGTGCACGAGTGGGTTACATCGAACTGGATCTCAACAGCGGTAAGATCCTT<br>GAGAGTTTTCGCCCCGAAGAACGTTTTCCAATGATGAGCACTTTTAAAGTTCTGCTATGTGGCGCGGTATTATCCCGTATTGACGCCGGGCAAGAGCAA<br>CTCGGTCGCCGCATACACTATTCTCAGAATGACTTGGTTGAGTACTCACCAGTCACAGAAAAGCATCTTACGGATGGCATGACAGTAAGAGAATTATGC<br>AGTGCTGCCATAACCATGAGTGATAACACTGCGGCCAACTTACTTCTGACAACGATCGGAGGACCGAAGGAGCTAACCGCTTTTTTGCACAACATGGG<br>GGATCATGTAACTCGCCTTGATCGTTGGGAACCGGAGCTGAATGAAGCCATACCAAACGACGAGCGTGACACCACGATGCCTGTAGCAATGGCAACAA<br>CGTTGCGCAAACTATTAACTGGCGAACTACTTACTCTAGCTTCCCGGCAACAATTAATAGACTGGATGGAGGCGGATAAAGTTGCAGGACCACTTCTGC<br>GCTCGGCCCTTCCGGCTGGCTGGTTTATTGCTGATAAATCTGGAGCCGGTGAGCGTGGGTCTCGCGGTATCATTGCAGCACTGGGGCCAGATGGTAA<br>GCCCTCCCGTATCGTAGTTATCTACACGACGGGGAGTCAGGCAACTATGGATGAACGAAATAGACAGATCGCTGAGATAGGTGCCTCACTGATTAAGC<br>ATTGGTAACTGTCAGACCAAGTTTACTCATATATACTTTAGATTGATTTAAAACTTCATTTTTAATTTAAAAGGATCTAGGTGAAGATCCTTTTTGATAATCT<br>CATGACCAAAATCCCTTAACGTGAGTTTTCGTTCCACTGAGCGTCAGACCCCGTAGAAAAGATCAAAGGATCTTCTTGAGATCCTTTTTTTCTGCGCGTA<br>ATCTGCTGCTTGCAAACAAAAAAACCACCGCTACCAGCGGTGGTTTGTTTGCCGGATCAAGAGCTACCAACTCTTTTTCCGAAGGTAACTGGCTTCAGC<br>AGAGCGCAGATACCAAATACTGTTCTTCTAGTGTAGCCGTAGTTAGGCCACCACTTCAAGAACTCTGTAGCACCGCCTACATACCTCGCTCTGCTAATC<br>CTGTTACCAGTGGCTGCTGCCAGTGGCGATAAGTCGTGTCTTACCGGGTTGGACTCAAGACGATAGTTACCGGATAAGGCGCAGCGGTCGGGCTGAA<br>CGGGGGGTTCGTGCACACAGCCCAGCTTGGAGCGAACGACCTACACCGAACTGAGATACCTACAGCGTGAGCTATGAGAAAGCGCCACGCTTCCCGA<br>AGGGAGAAAGGCGGACAGGTATCCGGTAAGCGGCAGGGTCGGAACAGGAGAGCGCACGAGGGAGCTTCCAGGGGGAAACGCCTGGTATCTTTATAG<br>TCCTGTCGGGTTTCGCCACCTCTGACTTGAGCGTCGATTTTTGTGATGCTCGTCAGGGGGGCGGAGCCTATGGAAAAACGCCAGCAACGCGGCCTTTT<br>TACGGTTCCTGGCCTTTTGCTGGCCTTTTGCTCACATGTTACCGTCGACGATGTAGGTCACGGTCTCGAAGCCGCGGTGCGGGTGCCAGGGCGTGCC<br>CTTGGGCTCCCCGGGCGCGTACTCCACCTCACCCATCTGGTCCATCATGATGAACGGGTCGAGGTGGCGGTAGTTGATCCCGGCGAACGCGCGGCG<br>CACCGGGAAGCCCTCGCCCTCGAAACCGCTGGGCGCGGTGGTCACGGTGAGCACGGGACGTGCGACGGCGTCGGCGGGTGCGGATACGCGGGGC<br>AGCGTCAGCGGGTTCTCGACGGTCACGGCGGGCATGTCGACAAGCCGAACATATGGGCGCGCCTAGTATGTATGTAAGTTAATAAAACCCATTTTTGC<br>GGAAAGTAGATAAAAAAAACATTTTTTTTTTTTTACTGCACTGGATATCATTGAACTTATCTGATCAGTTTTAAATTTACTTCGATCCAAGGGTATTTGATGTA<br>CCAGGTTCTTTCGATTACCTCTCACTCAAAATGACATTCCACTCAAAGTCAGCGCTGTTTGCCTCCTTCTCTGTCCACAGAAATATCGCCGTCTCTTTCG<br>CCGCTGCGTCCGCTATCTCTTTCGCCACCGTTTGTAGCGTTACGTAGCGTCAATGTCCGCCTTCAGTTGCATTTTGTCAGCGGTTTCGTGACGAAGCTC<br>CAAGCGGTTTACGCCATCAATTAAACACAAAGTGCTGTGCCAAAACTCCTCTCGCTTCTTATTTTTGTTTGTTTTTTGAGTGATTGGGGTGGTGATTGGTT<br>TTGGGTGGGTAAGCAGGGGAAAGTGTGAAAAATCCCGGCAATGGGCCAAGAGGATCAGGAGCTATTAATTCGCGGAGGCAGCAAACACCCATCTGCC<br>GAGCATCTGAACAATGTGAGTAGTACATGTGCATACATCTTAAGTTCACTTGATCTATAGGAACTGCGATTGCAACATCAAATTGTATGCGGCGTGAGAA<br>CTGCGACCCACAAAAATCCCAAACCGCAATTGCACAAACAAATAGTGACACGAAACAGATTATTCTGGTAGCTGTTCTCGCTATATAAGACAATTTTTGA<br>GATCATATCATGATCAAGACATCTAAAGGCATTCATTTTCGACTATATTCTTTTTTACAAAAAATATAACAACCAGATATTTTAAGCTGATCCTAGATGCAC<br>AAAAAATAAATAAAAGTATAAACCTACTTCGTAGGATACTTCGGGGTACTTTTTGTTCGGGGTTAGATGAGCATAACGCTAGTAGTTGATATTTGAGATCC<br>CCTATCATTGCAGGGTGACAGCGGAGCGGCTTCGCAGAGCTGCATTAACCAGGGCTTCGGGCAGGCCAAAAACTACGGCACGCTCCGGCCACCCAGT<br>CCGCCGGAGGACTCCGGTTCAGGGAGCGGCCAACTAGCCGAGAACCTCACCTATGCCTGGCACAATATGGACATCTTTGGGGCGGTCAATCAGCCGG<br>GCTCCGGATGGCGGCAGCTGGTCAACCGGACACGCGGACTATTCTGCAACGAGCGACACATACCGGCGCCCAGGAAACATTTGCTCAAGAACGGTGA<br>GTTTCTATTCGCAGTCGGCTGATCTGTGTGAAATCTTAATAAAGGGTCCAATTACCAATTTGAAACTCAGTTTGCGGCGTGGCCTATCCGGGCGAACTTT<br>TGGCCGTGATGGGCAGTTCCGGTGCCGGAAAGACGACCCTGCTGAATGCCCTTGCCTTTCGATCGCCGCAGGGCATCCAAGTATCGCCATCCGGGAT<br>GCGACTGCTCAATGGCCAACCTGTGGACGCCAAGGAGATGCAGGCCAGGTGCGCCTATGTCCAGCAGGATGACCTCTTTATCGGCTCCCTAACGGCC<br>AGGGAACACCTGATTTTCCAAGCCATGGTGCGGATGCCACGACATCTGACCTATCGGCAGCGAGTGGCCCGCGTGGATCAGGTGATCCAGGAGCTTT<br>CGCTCAGCAAATGTCAGCACACGATCATCGGTGTGCCCGGCAGGGTGAAAGGTCTGTCCGGCGGAGAAAGGAAGCGTCTGGCATTCGCCTCCGAGG<br>CTCTAACCGATCCGCCGCTTCTGATCTGCGATGAGCCCACCTCCGGACTGGACTCCTTTACCGCCCACAGCGTCGTCCAGGTGCTGAAGAAGCTGTCG<br>CAGAAGGGCAAGACCGTCATCCTGACCATTCATCAGCCGTCTTCCGAGCTGTTTGAGCTCTTTGACAAGATCCTTCTGATGGCCGAGGGCAGGGTAGC<br>TTTCTTGGGCACTCCCAGCGAAGCCGTCGACTTCTTTTCCTAGTGAGTTCGATGTGTTTATTAAGGGTATCTAGTATTACATAACATCTCAACTCCTATCC<br>AGCGTGGGTGCCCAGTGTCCTACCAACTACAATCCGGCGGACTTTTACGTACAGGTGTTGGCCGTTGTGCCCGGACGGGAGATCGAGTCCCGTGATC<br>GGATCGCCAAGATATGCGACAATTTTGCCATTAGCAAAGTAGCCCGGGATATGGAGCAGTTGTTGGCCACCAAAAATCTGGAGAAGCCACTGGAGCAG<br>CCGGAGAATGGGTACACCTACAAGGCCACCTGGTTCATGCAGTTCCGGGCGGTCCTGTGGCGATCCTGGCTGTCGGTGCTCAAGGAACCACTCCTCG<br>TAAAAGTGCGACTTATTCAGACAACGGTGAGTGGTTCCAGTGGAAACAAATGATATAACGCTTACAATTCTTGGAAACAAATTCGCTAGATTTTAGATAG<br>AATTGCCTGATTCCACACCCTTCTTAGTTTTTTTCAATGAGATGTATAGTTTATAGTTTTGCAGAAGATAAATAAATTTCATTTAACTCGCGAATATTAATGA<br>GATGCGAGTAACATTTTAATTTGCAGATGGTTGCCATCTTGATTGGCCTCATCTTTTTGGGCCAACAACTCACGCAAGTGGGTGTGATGAATATCAACGG<br>AGCCATCTTCCTCTTCCTGACCAACATGACCTTTCAAAACGTCTTTGCCACGATAAATGTAAGTCATGTTTAGAATACATTTGCATTTCAATAATTTACTAA<br>CTTTCTAATGAATCGATTCGATTTAGGTGTTCACCTCAGAGCTGCCAGTTTTTATGAGGGAGGCCCGAAGTCGACTTTATCGCTGTGACACATACTTTCT<br>GGGCAAAACGATTGCCGAATTGCCGCTTTTTCTCACAGTGCCACTGGTCTTCACGGCGATTGCCTATCCGATGATCGGACTGCGGGCCGGAGTGCTGC<br>ACTTCTTCAACTGCCTGGCGCTGGTCACTCTGGTGGCCAATGTGTCAACGTCCTTCGGATATCTAATATCCTGCGCCAGCTCCTCGACCTCGATGGCGC<br>TGTCTGTGGGTCCGCCGGTTATCATACCATTCCTGCTCTTTGGCGGCTTCTTCTTGAACTCGGGCTCGGTGCCAGTATACCTCAAATGGTTGTCGTACC<br>TCTCATGGTTCCGTTACGCCAACGAGGGTCTGCTGATTAACCAATGGGCGGACGTGGAGCCGGGCGAAATTAGCTGCACATCGTCGAACACCACGTG<br>CCCCAGTTCGGGCAAGGTCATCCTGGAGACGCTTAACTTCTCCGCCGCCGATCTGCCGCTGGACTACGTGGGTCTGGCCATTCTCATCGTGAGCTTCC<br>GGGTGCTCGCATATCTGGCTCTAAGACTTCGGGCCCGACGCAAGGAGTAGCCGACATATATCCGAAATAACTGCTTGTTTTTTTTTTTTACCATTATTAC |

| | |
|---|---|
| | CATCGTGTTTACTGTTTATTGCCCCCTCAAAAAGCTAATGTAATTATATTTGTGCCAATAAAAACAAGATATGACCTATAGAATACAAGTATTTCCCCTTCG<br>AACATCCCCACAAGTAGACTTTGGATTTGTCTTCTAACCAAAAGACTTACACACCTGCATACCTTACATCAAAAACTCGTTTATCGCTACATAAAACACCG<br>GGATATATTTTTTATATACATACTTTTCAAATCGCGCGCCCTCTTCATAATTCACCTCCACCACACCACGTTTCGTAGTTGCTCTTTCGCTGTCTCCCACC<br>CGCTCTCCGCAACACATTCACCTTTTGTTCGACGACCTTGGAGCGACTGTCGTTAGTTCCGCGCGATTCGGTTCGCTCAAATGGTTCCGAGTGGTTCAT<br>TTCGTCTCAATAGAAATTAGTAATAAATATTTGTATGTACAATTTATTTGCTCCAATATATTTGTATATATTTCCCTCACAGCTATATTTATTCTAATTTAATA<br>TTATGACTTTTTAAGGTAATTTTTTGTGACCTGTTCGGAGTGATTAGCGTTACAATTTGAACTGAAAGTGACATCCAGTGTTTGTTCCTTGTGTAGATGCA<br>TCTCAAAAAAATGGTGGGCATAATAGTGTTGTTTATATATATCAAAAATAACAACTATAATAATAAGAATACATTTAATTTAGAAAATGCTTGGATTTCACTG<br>GAACTAGGGCGCGCCTCCGGAACATAATGGTGCAGGGCGCTGACTTCCGCGTTTCCAGACTTTACGAAACACGGAAACCGAAGACCATTCATGTTGTT<br>GCTCAGGTCGCAGACGTTTTGCAGCAGCAGTCGCTTCACGTTCGCTCGCGTATCGGTGATTCATTCTGCTAACCAGTAAGGCAACCCCGCCAGCCTAG<br>CCGGGTCCTCAACGACAGGAGCACGATCATGCGCACCCGTGGCCAGGGCCGCAAGCTTGCATGCCTGCAGACATCTTGTCGCCGGAACGCAGCGAC<br>AGGGATTCCAATGTGTCCGTATCCTTTAGGCTTTTGCCCTTCAGTTCCAGACGAAGCGACTGGCGATTCGCGTGTGGGGTCTGCTTCAGGGTCTTGTGA<br>ATTAGGGCGCGCAGATCGCCGATGGGCGTGGCGCCGGAGGGCACCTTCACCTTGCCGTACGGCTTGCTGTTCTTCGCGTTCAAAATCTCCAGCTCCA<br>TTTTGCTTTCGGTGCGCTTGCAATCAGTACTGTCCAAAATCGAAAATCGCCGAACCGTAGTGTGACCGTGCGGGGCTCTGCGAAAATAAACTTTTTTAG<br>GTATATGGCCACACACGGGGAAAGCACAGTGGATTATATGTATTAATATATTATCCAGGTTTTCATTACTTATCCAGATGTAAGCCCACTTAAAGCGATTT<br>AACAATTATTTGCCGAAAGAGTATAAACAAATTTTACTTAAAAATGGATTAAGAAAAGCTTGTGTAAGATTATGCGCAGCGTTGCCAGATAGCTCCATTTA<br>AAACACTTCAAAAACAATAAGTTTTGAAAATATATACATAAATAGCAGTCGTTGCCGCAACGCTCAACACATCACACTTTTAAAACACCCTTTACCTACAC<br>AGAATTACTTTTTAAATTTCCAGTCAAGCTGCGAGTTTCAAAATTATAGCCGGTAGAGAAGACAGTGCTATTTCAAAAGCAAACTAACAAGGGTCTTAAAT<br>TCCAAAACACCAATCCTAACAAGCCTTGGACTTTTGTAAGTTTAGATCAAAGGTGGCATTGCATTCAATGTCATGGTAAGAAGTAGGTCGTCTAGGTAGA<br>AATCCTCATTCAGCCGGTCAAGTCAGTACGAGAAAGGTCTCAATTTGAAATTGTCTTAAAAATATTTTATTGTTTTGTACTGTGGTGAGTTTAAACGAAAA<br>ACACAAAAAAAAAGTGATACATAGAAATCATAAAAAATTTTAATACAAGGTATTCGTACGTATCAAAAACATTTCGGCACAATTTTTTTTCTCTGTACTAAA<br>GTGTTACGAACACTACGGTATTTTTTAGTGATTTTCAACGGACACCGAAGGTATATAAACAGCGTTCGCGAACGGTCGCCTTCAAAACCAATTGACATTT<br>GCAGCAGCAAGTACAAGCAGAAAGTAAAGCGCAATCAGCGAAAAATTTATACTTAATTGTTGGTGATTAAAGTACAATTAAAAGAACATTCTCGAAAGTC<br>ACAAGAAACGTAAGTTTTTAACTCGCTGTTACCAATTAGTAATAAGAGCAACAAGACGTTGAGTAATTTCAAGAAAAACTGCATTTCAAGGTCTTTGTTCG<br>GCCATTTTTTTTTATTCAACGCTCTACGTAATTACAAAATAAGAAATTGGCAGCCACGCATCTCGTTTTCCCAATGAATTGGCATCAAAACGCAAACAAAT<br>CTATAAATAAAACTTACGTGTTGATTTTCGCCAAGATTTATTGGCAAATTGTGAAATTCGCAGTGACGTATTTGAAAATTCGAGAAATCACATACGCACTC<br>GAGCATTTGTGTGCATGTTATTAGTTAGTTGTTTAGTTAATTGAAGTATTTTACCAACGAAATCCACTTATTTTTAGCTGAAATAGAGTAGGTTGCTTAAAC<br>AAAGCCACGTCTGAAAATTTCTTATTGCTTGTAGTTGTGACGTCACCATATACACACAAAATAATGTGTATGCATGCATTTCAGCTGTGTATATATACATG<br>CACACACTCGCAACACGAAAACGATGACGAAGCAACGGAACAAAGGTTTCTCAACTACCCTTTGTTCCCTGTTTCTTCGCTTTCCTTTGTTCCAATATTC<br>GTAGAGGGTTAATAGGGGTTTCTCAACAAAGTTGGCGTCGATAAATAAGTTTCCCATTTTTATTCCCCAGCCAGGAAGTTAGTTTCAATAGTTTTGTAATT<br>TCAACGAAACTCATTTGATTTCGTACTAATTTTCCACATCTCTATTTTCTGCCCGCAGAATAATCCACGGGTCACCACTGGATCCGTCACCGGTAGCGCG<br>ATCGCAGTCCTAGGATGAATCGTTTTTAAAATAACAAATCAATTGTTTTATAATATTCGTACGATTCTTTGATTATGTAATAAAATGTGATCATTAGGAAGAT<br>TACGAAAAATATAAAAAATATGAGTTCTGTGTGTATAACAAATGCTGTAAACGCCACAATTGTGTTTGTTGCAAATAAACCCATGATTATTTGATTAAAATT<br>GTTGTTTTCTTTGTTCATAGACAATAGTGTGTTTTGCCTAAACGTGTACTGCATAAACTCCATGCGAGTGTATAGCGAGCTAGTGGCTAACGCTTGCCCC<br>ACCAAAGTAGATTCGTCAAAATCCTCAATTTCATCACCCTCCTCCAAGTTTAACATTTGGCCGTCGGAATTAACTTCTAAAGATGCCACATAATCTAATAA<br>ATGAAATAGAGATTCAAACGTGGCGTCATCGTCCGTTTCGACCATTTCCGAAAAGAACTCGGGCATAAACTCTATGATTTCTCTGGACGTGGTGTTGTCG<br>AAACTCTCAAAGTACGCAGTCAGGAACGTGCGCGACATGTCGTCGGGAAACTCGCGCGGAAACATGTTGTTGTAACCGAACGGGTCCCATAGCGCCAA<br>AACCAAATCTGCCAGCGTCAATAGAATGAGCACGATGCCGACAATGGAGCTGGCTTGGATAGCGATTCGAGTTAACTTAATTAATGGCCACGTAATAAG<br>TGTGCGTTGAATTTATTCGCAAAAACATTGCATATTTTCGGCAAAGTAAAATTTTGTTGCATACCTTATCAAAAAATAAGTGCTGCATACTTTTTAGAGAAA<br>CCAAATAATTTTTTATTGCATACCCGTTTTTAATAAAATACATTGCATACCCTCTTTTAATAAAAAAATATTGCATACTTTGACGAAACAAATTTTCGTTGCAT<br>ACCCAATAAAAGATTATTATATTGCATACCCGTTTTTAATAAAATACATTGCATACCCTCTTTTAATAAAAAAATATTGCATACGTTGACGAAACAAATTTTCG<br>TTGCATACCCAATAAAAGATTATTATATTGCATACCTTTTCTTGCCATACCATTTAGCCGATCAATTGTGCTCGGCAACAGTATATTTGTGGTGTGCCAAC<br>TAGAGGTTTATCGGATGTTTACAGCAGTATTGGGTTTTTATTCCCTTTGGGTCTTAGGGGGCTTGGACAAGCTCTGGATTCTGGTCAGATGTTTGAATTG<br>TGAATATAAGCTCTCATTCACCGCTCGTTCAATCCGACGGGATAGGCGCACTCTACCAATAGAATTTAGCTCGTGTGTGGGCCCTTATATTTTTCCCCAA<br>ACAGACATAAGCGCAGCAGAGCCAAATCGCTAATATCGGACTATATTGGCATTGCGCTGGTTAAGCAAGGAGTGTACCTGCCGGTCTTATAGAATCTTT<br>AGTGTAATAAGTCACTGGGCGCATTAACAATACTGTATTGGGAGGATGTGCATGAGTTATGCTCATTCTGCAGAAAATAGATGCTGGTCAAATCCTTACT<br>GGAGATGGCGCAGGATTTCATTTTGCAATTGTGGCATGGCACCTTAGGATCGCTTGCTATACGCTAAGACCGCAGCTACATCAGGAGTCCACAAATGCA<br>CATTAGGAGCGAATTTTTACTTCTTTAGATTTTTAGTTCTCCGATGAAACCCCGAAAGTCGAGTGGACTTTGCTTTCTTATGCTTTGAGTGTGAAAGTAAT<br>ACCTCACAAATTCGAGGTTTGATCAAGGACTCATAAGGTGAGATCCTGACTGTTTACGCCTCTAGCACGGCAGCTTGCAGTGTCAATATGTTGCCCGAT<br>CCCTATTATAGAATATGAGAGGAATACATATCGAGTCGGACATACTCTTGTCTCTGGCGCCTGAAGTCGGTGGGCATCAAATACCAGTAGAAGTCCGAG<br>GTTATAATATATATGATCTACGTTTCTAGGTGTTTCGAAGGTCGGAGGGTTGGGTCGCAGGTCGAGCATCAGTTACCCCGTTGTCCCCTTATACGTTTCG<br>CAATACGTGGAAAGTGCTGGGCTCAGCGCAGGTCCACCAGCGCAGGCATGCTTGGGCGCGGAGAGCAGCATTGGAGACCGCATCCATCCCAAGGAC<br>AGCCGCGATCGACTGGAGGCCGCGCTTTATGATTAGGATATTATGCACATAGCGAGTCCGACTTCGCGCGCCGTGCCGATATTGAATAGTTTTCGTTTG<br>TGGCAGATAGAGGGTGTCCTGTCAACTGTTGGGCCGGATTAGTGATGACGGTATAGTGTCGTTGTACACCTTTCGTATCTAGGTAGTGACACCGACTTC<br>ATAGATTCGCAAGCGAGGGAGTTGCGATCCCCCTCTCTGCCCTTCAATACTGGAAACATCTCGATATGCAAACCGCAACCGTGGATCATCAGGTTCGCC<br>GACACAGACTAAGGACACTACCAAGCAGAACTCTGGAAGTTGACAGTGGGTTGGACTACCCGGTACACGGGGGCCGTGGGCGGAAATGTCGAAAAGT<br>TGGTGCACCGTATTTCTTGTATACGCGAACTAGCACTCCCCTTGTCGCGTACCGTACACACAGCCCCCATTCGGAGCCGACCTTATGAGCTACGTGAG<br>GCGGCGGCGACTAACGCTAACGCGCCAACGCCTTAGTCTCGCACACCTCCGTGTTGGAGCGCAAATGTAAGCGTTAGGCCTGTCAAAGCGTCACCGA<br>AGGGGCCCCAAAACTGACATGGGCTTGGCCGAACCGGCCGGTGGGTTGGAGAGATGGAACAGGGCCGGACTGCTTCAGGGTTACAATCCCTCAAAGT<br>GATTACCTGAAACCGCAGGCCCTCTAGCGGTTTGTACTCTCTACGTCCGCAGCACCTCTGTGATGGGTTCAGAGGAATCGTGGAGTCGACCTCCGCTG<br>ACGGCAAAGAGGAATCGTGCTCCAGCACACCTCGCAGATACCGGCGCTCTGCGCGACCCTGATCCATACAGGAACGGTCGGTGCATGCCCGTGCCGA<br>GGTCTACGAGTTAGTACGGACCATTAGAAAAAAGCGTGCCGCGCGGGTCCTCCGGCAGATAGTGTTGGGGATGCTGGACTGTGGAAACCGATTCAGG<br>GGCTACGCGTCGGCCCCTATCCCGAGGTGCCGCCGCGCTTCTGGCCACGTAATAAGTGTGCGTTGAATTTATTCGCAAAAACATTGCATATTTTCGGCA<br>AAGTAAAATTTTGTTGCATACCTTATCAAAAAATAAGTGCTGCATACTTTTTAGAGAAACCAAATAATTTTTTATTGCATACCCGTTTTTAATAAAATACATT<br>GCATACCCTCTTTTAATAAAAAAATATTGCATACTTTGACGAAACAAATTTTCGTTGCATACCCAATAAAAGATTATTATATTGCATACCCGTTTTTAATAAAA<br>TACATTGCATACCCTCTTTTAATAAAAAAATATTGCATACGTTGACGAAACAAATTTTCGTTGCATACCCAATAAAAGATTATTATATTGCATACCTTTTCTTG<br>CCATACCATTTAGCCGATCAATTGTGCTCGGCAACAGTATATTTGTGGTGTGCCAACATTTAAATGGCCGGCCGTTTGCCGTTAACTCGAATCGCTATCC<br>AAGCCAGCTCCATTGTCGGCATCGTGCTCATTCTATTGACGCTGGCAGATTTGGTTTTGGCGCTATGGGACCCGTTCGGTTACAACAACATGTTTCCGC<br>GCGAGTTTCCCGACGACATGTCGCGCACGTTCCTGACTGCGTACTTTGAGAGTTTCGACAACACCACGTCCAGAGAAATCATAGAGTTTATGCCCGAGT<br>TCTTTTCGGAAATGGTCGAAACGGACGATGACGCCACGTTTGAATCTCTATTTCATTTATTAGATTATGTGGCATCTTTAGAAGTTAATTCCGACGGCCAA<br>ATGTTAAACTTGGAGGAGGGTGATGAAATTGAGGATTTTGACGAATCTACTTTGGTGGGGCAAGCGTTAGCCACTAGCTCGCTATACACTCGCATGGAG<br>TTTATGCAGTACACGTTTAGGCAAAACACACTATTGTCTATGAACAAAGAAAACAACAATTTTAATCAAATAATCATGGGTTTATTTGCAACAAACACAATT<br>GTGGCGTTTACAGCATTTGTTATACACACAGAACTCATATTTTTTATATTTTTCGTAATCTTCCTAATGATCACATTTTATTACATAATCAAAGAATCGTACG<br>AATATTATAAAACAATTGATTTGTTATTTTAAAAACGATTCATAGGCCTTCTAGACTGAGCTCATGATATCATGCTAGCATGCGGCCGCATAGATCTCTGA<br>ATTCATGGTACCCGTGGATTATTCTGCGGGCAGAAAATAGAGATGTGGAAAATTAGTACGAAATCAAATGAGTTTCGTTGAAATTACAAAACTATTGAAA<br>CTAACTTCCTGGCTGGGGAATAAAAATGGGAAACTTATTTATCGACGCCAACTTTGTTGAGAAACCCCTATTAACCCTCTACGAATATTGGAACAAAGGA<br>AAGCGAAGAAACAGGGAACAAAGGGTAGTTGAGAAACCTTTGTTCCGTTGCTTCGTCATCGTTTTCGTGTTGCGAGTGTGTGCATGTATATATACACAG<br>CTGAAATGCATGCATACACATTATTTTGTGTGTATATGGTGACGTCACAACTACAAGCAATAAGAAATTTTCAGACGTGGCTTTGTTTAAGCAACCTACTC<br>TATTTCAGCTAAAAATAAGTGGATTTCGTTGGTAAAATACTTCAATTAACTAAACAACTAACTAATAACATGCACACAAATGCTCGAGTGCGTATGTGATTT<br>CTCGAATTTTCAAATACGTCACTGCGAATTTCACAATTTGCCAATAAATCTTGGCGAAAATCAACACGTAAGTTTTATTTATAGATTTGTTTGCGTTTTGAT<br>GCCAATTCATTGGGAAAACGAGATGCGTGGCTGCCAATTTCTTATTTTGTAATTACGTAGAGCGTTGAATAAAAAAAAATGGCCGAACAAAGACCTTGAA<br>ATGCAGTTTTTCTTGAAATTACTCAACGTCTTGTTGCTCTTATTACTAATTGGTAACAGCGAGTTAAAAACTTACGTTTCTTGTGACTTTCGAGAATGTTCT<br>TTTAATTGTACTTTAATCACCAACAATTAAGTATAAATTTTTCGCTGATTGCGCTTTACTTTCTGCTTGTACTTGCTGCTGCAAATGTCAATTGGTTTTGAA<br>GGCGACCGTTCGCGAACGCTGTTTATATACCTTCGGTGTCCGTTGAAAATCACTAAAAAATACCGTAGTGTTCGTAACACTTTAGTACAGAGAAAAAAAA<br>TTGTGCCGAAATGTTTTTGATACGTACGAATACCTTGTATTAAAATTTTTTATGATTTCTATGTATCACTTTTTTTTTGTGTTTTTCGTTTAAACTCACCACA<br>GTACAAAACAATAAAATATTTTTAAGACAATTTCAAATTGAGACCTTTCTCGTACTGACTTGACCGGCTGAATGAGGATTTCTACCTAGACGACCTACTTC<br>TTACCATGACATTGAATGCAATGCCACCTTTGATCTAAACTTACAAAAGTCCAAGGCTTGTTAGGATTGGTGTTTTGGAATTTAAGACCCTTGTTAGTTTG<br>CTTTTGAAATAGCACTGTCTTCTCTACCGGCTATAATTTTGAAACTCGCAGCTTGACTGGAAATTTAAAAAGTAATTCTGTGTAGGTAAAGGGTGTTTTAA<br>AAGTGTGATGTGTTGAGCGTTGCGGCAACGACTGCTATTTATGTATATATTTTCAAAACTTATTGTTTTTGAAGTGTTTTAAATGGAGCTATCTGGCAACG<br>CTGCGCATAATCTTACACAAGCTTTTCTTAATCCATTTTTAAGTAAAATTTGTTTATACTCTTTCGGCAAATAATTGTTAAATCGCTTTAAGTGGGCTTACAT<br>CTGGATAAGTAATGAAAACCTGGATAATATATTAATACATATAATCCACTGTGCTTTCCCCGTGTGTGGCCATATACCTAAAAAAGTTTATTTTCGCAGAG<br>CCCCGCACGGTCACACTACGGTTCGGCGATTTTCGATTTTGGACAGTACTGATTGCAAGCGCACCGAAAGCAAAATGGAGCTGGAGATTTTGAACGCG<br>AAGAACAGCAAGCCGTACGGCAAGGTGAAGGTGCCCTCCGGCGCCACGCCCATCGGCGATCTGCGCGCCCTAATTCACAAGACCCTGAAGCAGACCC<br>CACACGCGAATCGCCAGTCGCTTCGTCTGGAACTGAAGGGCAAAAGCCTAAAGGATACGGACACATTGGAATCCCTGTCGCTGCGTTCCGGCGACAA<br>GATGTGCCAAACGAATTCTTGAAGACGAAAGGGCCTCGTGATACGCCTATTTTTATAGGTTAATGTCATGATAATAATGGTTTCTTA |
| WT PGANT3 | atgggtctccggttccagcagctgaagaagctctggctgctctaccttttcctgctcttcttcgcgttcttcatgttcgccatcagcatcaacctgtacgtggccagcatccagggcggggacgcggagatgcgtcatccgaaaccg<br>ccgcccaagcgccgctccctctggccgcacaaaaacatcgtggcccactacatcggcaagggagacatctttggcaacatgacagcagatgattacaacatcaacctgttccagcccatcaacggagaaggtgccgatg<br>ggcgaccagttgtggtgccgcctcgcgatcgctttcgcatgcagcgctttttccggctaaacagcttcaacctgctggccagcgaccgcatcccactgaaccgcaccctcaaggactaccgcacacctgaatgccgcgacaa<br>gaagtacgccagcggtctacccagcacgtcggttataattgttttccacaacgaggcctggtccgtgctcctgcgaaccatcaccagcgtcatcaaccggtcgccgcgccacctgcttaaggaaattatcctggtggacgatgc<br>cagcgatcgctcatatttaaaacgtcaactggagagctatgtgaaagtgctggcggtacccacgaggatctttcgcatgaagaagcgaagtggcttagtgcctgcccgccttttgggagccgaaaatgctcgcggcgatgtgc<br>tgaccttcctcgacgcccactgcgagtgctcgcggggatggctggaaccacttctttcccgcattaaggaatcacgcaaggtggtaatctgcccggtaatcgacataatatccgatgacaactttagctacacaaaaaccttcg<br>agaaccactggggcgctttcaactggcagctgagcttccgttggttctcctcggatcgtaagcggcagacagcgggcaacagctccaaggatagcactgacccgattgccactcccggaatggccggcggattgtttgctatc<br>gatcgcaagtacttttacgaaatgggatcctacgacagcaacatgcgcgtttggggcggcgagaatgtggaaatgtccttccgcatttggcagtgcggcggtcgtgtggagatcagtccctgctcgcacgtgggtcacgtgttc<br>cgcagcagcacgccctacaccttccccggcggcatgagcgaggttctcacggataatttggccagggccgccaccgtctggatggacgattggcagtactttataatgctgtacacctctggactgacgctgggagccaagg<br>acaaggtgaatgttacggagcgtgtggcgctcagggagaggctccagtgcaaaccgttctcctggtatctggagaacatatggccagagcacttctttccagctcccgaccgcttttcggcaagatcatatggctggacgggg<br>agacagaatgtgcccaggcctacagcaagcacatgaagaacctaccaggtcgtgctttgtcacgagagtggaagcgagcgttcgaggaaatcgacagcaaggcggaggagctgatggcgctgatcgatcgatctggagcgg<br>gacaagtgccttcgccccctcaaagaggatgtgccgcgctcatcattgtctgcagtaaccgtaggcgattgcacatctcacgcccagtcgatggatatgtttgtaataactccgaaggccagatcatgactaacgacaacgtc<br>tgcctgacctatcgacagcaaaagctgggtgtgatcaagatgctaaagaaccgcaacgccaccacttcgaatgtaatgctggcgcagtgcgcttccgactcgagtcagttgtggacctacgacatggatactcagcaaatat<br>ctcatagggacacaaagctctgccttactctgaaagcggcgactaactcgcgtctccaaaaggtcgagaaggttgtgctcagtatggaatgcgacttcaaagacatcacacagaagtggggcttcataccactgccatggcg<br>catg |

| | |
|---|---|
| BH PGANT3 | atgggtctccggttccagcagctgaagaagctctggctgctctaccttttcctgctcttcttcgcgttcttcatgttcgccatcagcatcaacctgtacgtggccagcatccagggcggggacgcggagatgcgtcatccgaaaccgccgcccaagcgccgctccctctggccgcacaaaaacatcgtggcccactacatcggcaagggagacatctttggcaacatgacagcagatgattacaacatcaacctgttccagcccatcaacggagaaggtgccgatgggcgaccagttgtggtgccgcctcgcgatcgctttcgcatgcagcgctttttccggctaaacagcttcaacctgctggccagcgaccgcatcccactgaaccgcaccctcaaggactaccgcacacctgaatgccgcgacaagaagtacgccagcggtctacccagcacgtcggttataattgttttccacaacgaggcctggtccgtgctcctgcgaaccatcaccagcgtcatcaaccggtcgccgcgccacctgcttaaggaaattatcctggtggacgatgccagcgatcgctcatatttaaaacgtcaactggagagctatgtgaaagtgctggcggtacccacgaggatctttcgcatgaagaagcgaagtggcttagtgcctgcccgccttttgggagccgaaaatgctcgcggcgatgtgctgaccttcctcgacgcccactgcgagtgctcgcggggatggctggaaccacttctttcccgcattaaggaatcacgcaaggtggtaatctgcccggtagccgacataatatccgatgacaactttagctacacaaaaaccttcgagaaccactggggcgctttcaactggcagctgagcttccgttggttctcctcggatcgtaagcggcagacagcgggcaacagctccaaggatagcactgacccgattgccactcccggaatggccggcggagcctttgctatcgatcgcaagtacttttacgaaatgggatcctacgacagcaacatgcgcgtttggggcggcgagaatgtggaaatgtccttccgcatttggcagtgcggcggtcgtgtggagatcagtccctgctcgcacgtgggtcacgtgttccgcagcagcacgccctacaccttccccggcggcatgagcgaggttctcacggataatttggccagggccgccaccgtctggatggacgattggcagtactttataatgctgtacacctctggactgacgctgggagccaaggacaaggtgaatgttacggagcgtgtggcgctcagggagaggctccagtgcaaaccgttctcctggtatctggagaacatatggccagagcacttctttccagctcccgaccgctttttcggcaagatcatatggctggacggggagacagaatgtgcccaggcctacagcaagcacatgaagaacctaccaggtcgtgctttgtcacgagagtggaagcgagcgttcgaggaaatcgacagcaaggcggaggagctgatggcgctgatcgatctggagcgggacaagtgccttcgcccccctcaaagaggatgtgccgcgctcatcattgtctgcagtaaccgtaggcgattgcacatctcacgcccagtcgatggatatgtttgtaataactccgaagggccagatcatgactaacgacaacgtctgcctgacctatcgacagcaaaagctgggtgtgatcaagatgctaaagaaccgcaacgccaccacttcgaatgtaatgctggcgcagtgcgcttccgactcgagtcagttgtggacctacgacatggatactcagcaaatatctcatagggacacaaagctctgccttactctgaaagcggcgactaactcgcgtctccaaaaggtcgagaaggttgtgctcagtatggaatgcgacttcaaagacatcacacagaagtggggcttcataccactgccatggcgcatg |
| WT PGANT9A | atggccttcatctggcggcgacgttcgacgacgattgtgaagctggtggccttcgccctggccatttggttttgcattgccttcctggtatacacggatgacacccggcgtagagccgcccaggagggggctggtgccagtggggcaggcagtgctcccggagtgggcggcggagccggaggactgggtgatcccattgccttggctctgagaaacgagcctgctggcgaggactttggcatcaatgggaatgtcatcggcggaggaggacagaagcaggcgcacgatgaggccgacatcccacccactgtgggaaaacataaggctgacctacaggcggaaaggatgcggaagaaggcggccgagcagccaaagaaaaagccacaagaagatagcaaaaaagtgattgatccgccggcgaactttgaggagaatcctggcgagctgggaaaacccgtgcgcctgcccaaggaaatgtccgacgaaatgaagaaggccgttgacgatggctggaccaaaaacgccttcaatcaatatgtatcggatctcatatccgttcatcgaactctgcccgatcctcgagatgcctggtgcaaggacgaggcgcgctatctaaccaatctgccaaagaccgatgtcatcatctgcttccacaacgaagcctggaccgtgctgttgagaacagttcactcagtgctggacaggtcgccggaacatctaatagggaagataatcctcgtcgatgactacagcgatatgcctcatttgaagaggcagctggaggactactttgctgcctatcccaaagtccagataattcgtggccaaaagcgagagggtctgatccgagcccgaatttgggtgccaaccatgccaagtcccctgttctcacctatttggattcccactgcgagtgcaccgaaggatggcttgaaccacttttggaccgaattgcacgtaattccaccactgtcgtgtgtcccgttattgatgtcattagcgatgagaccctggagtaccactatcgcgattcgggcggagttaatgtcggaggattcgattggaatcttcagttcagctggcatccggttcccgaacgtgaaaggaagcgccacaatagcaccgccgaacccgtctactcacccacaatggccggcggacttttctccatcgacagagagttcttcgatcgactgggcacctatgactctggatttgatatttggggcggtgagaatctggagctatccttcaagacctggatgtgcggcggcaccctggagatagttccctgctctcacgtgggacacatattccggaaacgttcgccctacaagtggcgttctggcgtgaatgtgctgaagaagaactccgtgcggttggccgaggtttggatggatgagtactcgcagtactactatcatcgcatcggcaacgacaagggcgactggggcgatgttagcgatcgcaggaagctccgtaacgatctcaagtgcaaaagcttcaaatggtatctggacaacatttatcccgagctcttcattcccggcgattctgtggcccacggagagataagaaatctaggttatggcggtcgcacctgcttggacgcaccggctggcaagaagcaccagaagaaggcggtgggcacgtatccctgccatcggcagggtggaaaccagtactggatgctcagcaaggcgggcgaaatccgtcgcgacgactcctgtctcgattatgccggcaaggatgtgacgcttttcggctgccacgggggcaaaggaaatcagttctggacgtaccgcgagaacacaaagcagctgcaccacggcacctccggcaagtgtctggccatcagcgagagcaaggacaagctgctgatggaggagtgcagcgcgagcctcagtcgccaacagtggaccctcgagaactacgacagctcgaagttg |
| BH PGANT9A | atggccttcatctggcggcgacgttcgacgacgattgtgaagctggtggccttcgccctggccatttggttttgcattgccttcctggtatacacggatgacacccggcgtagagccgcccaggagggggctggtgccagtggggcaggcagtgctcccggagtgggcggcggagccggaggactgggtgatcccattgccttggctctgagaaacgagcctgctggcgaggactttggcatcaatgggaatgtcatcggcggaggaggacagaagcaggcgcacgatgaggccgacatcccacccactgtgggaaaacataaggctgacctacaggcggaaaggatgcggaagaaggcggccgagcagccaaagaaaaagccacaagaagatagcaaaaaagtgattgatccgccggcgaactttgaggagaatcctggcgagctgggaaaacccgtgcgcctgcccaaggaaatgtccgacgaaatgaagaaggccgttgacgatggctggaccaaaaacgccttcaatcaatatgtatcggatctcatatccgttcatcgaactctgcccgatcctcgagatgcctggtgcaaggacgaggcgcgctatctaaccaatctgccaaagaccgatgtcatcatctgcttccacaacgaagcctggaccgtgctgttgagaacagttcactcagtgctggacaggtcgccggaacatctaatagggaagataatcctcgtcgatgactacagcgatatgcctcatttgaagaggcagctggaggactactttgctgcctatcccaaagtccagataattcgtggccaaaagcgagagggtctgatccgagcccgaatttgggtgccaaccatgccaagtcccctgttctcacctatttggattcccactgcgagtgcaccgaaggatggcttgaaccacttttggaccgaattgcacgtaattccaccactgtcgtgtgtcccgttgccgatgtcattagcgatgagaccctggagtaccactatcgcgattcgggcggagttaatgtcggaggattcgattggaatcttcagttcagctggcatccggttcccgaacgtgaaaggaagcgccacaatagcaccgccgaacccgtctactcacccacaatggccggcggagccttctccatcgacagagagttcttcgatcgactgggcacctatgactctggatttgatatttggggcggtgagaatctggagctatccttcaagacctggatgtgcggcggcaccctggagatagttccctgctctcacgtgggacacatattccggaaacgttcgccctacaagtggcgttctggcgtgaatgtgctgaagaagaactccgtgcggttggccgaggtttggatggatgagtactcgcagtactactatcatcgcatcggcaacgacaagggcgactggggcgatgttagcgatcgcaggaagctccgtaacgatctcaagtgcaaaagcttcaaatggtatctggacaacatttatcccgagctcttcattcccggcgattctgtggcccacggagagataagaaatctaggttatggcggtcgcacctgcttggacgcaccggctggcaagaagcaccagaagaaggcggtgggcacgtatccctgccatcggcagggtggaaaccagtactggatgctcagcaaggcgggcgaaatccgtcgcgacgactcctgtctcgattatgccggcaaggatgtgacgcttttcggctgccacgggggcaaaggaaatcagttctggacgtaccgcgagaacacaaagcagctgcaccacggcacctccggcaagtgtctggccatcagcgagagcaaggacaagctgctgatggaggagtgcagcgcgagcctcagtcgccaacagtggaccctcgagaactacgacagctcgaagttg |
| WT PGANT9B | atggccttcatctggcggcgacgttcgacgacgattgtgaagctggtggccttcgccctggccatttggttttgcattgccttcctggtatacacggatgacacccggcgtagagccgcccaggagggggctggtgccagtggggcaggcagtgctcccggagtgggcggcggagccggaggactgggtgatcccattgccttggctctgagaaacgagcctgctggcgaggactttggcatcaatgggaatgtcatcggcggaggaggacagaagcaggcgcacgatgaggccgacatcccacccactgtgggaaaacataaggctgacctacaggcggaaaggatgcggaagaaggcggccgagcagccaaagaaaaagccacaagaagatagcaaaaaagtgattgatccgccggcgaactttgaggagaatcctggcgagctgggaaaacccgtgcgcctgcccaaggaaatgtccgacgaaatgaagaaggccgttgacgatggctggaccaaaaacgccttcaatcaatatgtatcggatctcatatccgttcatcgaactctgcccgatcctcgagatgcctggtgcaaggacgaggcgcgctatctaaccaatctgccaaagaccgatgtcatcatctgcttccacaacgaagcctggaccgtgctgttgagaacagttcactcagtgctggacaggtcgccggaacatctaatagggaagataatcctcgtcgatgactacagcgatatgcctcatttgaagaggcagctggaggactactttgctgcctatcccaaagtccagataattcgtggccaaaagcgagagggtctgatccgagcccgaatttgggtgccaaccatgccaagtcccctgttctcacctatttggattcccactgcgagtgcaccgaaggatggcttgaaccacttttggaccgaattgcacgtaattccaccactgtcgtgtgtcccgttattgatgtcattagcgatgagaccctggagtaccactatcgcgattcgggcggagttaatgtcggaggattcgattggaatcttcagttcagctggcatccggttcccgaacgtgaaaggaagcgccacaatagcaccgccgaacccgtctactcacccacaatggccggcggacttttctccatcgacagagagttcttcgatcgactgggcacctatgactctggatttgatatttggggcggtgagaatctggagctatccttcaagacctggatgtgcggcggcaccctggagatagttccctgctctcacgtgggacacatattccggaaacgttcgccctacaagtggcgttctggcgtgaatgtgctgaagaagaactccgtgcggttggccgaggtttggatggatgagtactcgcagtactactatcatcgcatcggcaacgacaagggcgactggggcgatgttagcgatcgcaggaagctccgtaacgatctcaagtgcaaaagcttcaaatggtatctggacaacatttatcccgagctcttcattcccggcgattctgtggcccacggagagatcgcgaatgtgcccaacggcatgtgtctggatgcaaaagaaaaatccgaagaggaaacaccggtttccatttatgagtgccatggccaaggaggaaatcagtactggatgctcagcaaggcgggcgaaatccgtcgcgacgactcctgtctcgattatgccggcaaggatgtgacgcttttcggctgccacgggggcaaaggaaatcagttctggacgtaccgcgagaacacaaagcagctgcaccacggcacctccggcaagtgtctggccatcagcgagagcaaggacaagctgctgatggaggagtgcagcgcgagcctcagtcgccaacagtggaccctcgagaactacgacagctcgaagttg |
| BH PGANT9B | atggccttcatctggcggcgacgttcgacgacgattgtgaagctggtggccttcgccctggccatttggttttgcattgccttcctggtatacacggatgacacccggcgtagagccgcccaggagggggctggtgccagtggggcaggcagtgctcccggagtgggcggcggagccggaggactgggtgatcccattgccttggctctgagaaacgagcctgctggcgaggactttggcatcaatgggaatgtcatcggcggaggaggacagaagcaggcgcacgatgaggccgacatcccacccactgtgggaaaacataaggctgacctacaggcggaaaggatgcggaagaaggcggccgagcagccaaagaaaaagccacaagaagatagcaaaaaagtgattgatccgccggcgaactttgaggagaatcctggcgagctgggaaaacccgtgcgcctgcccaaggaaatgtccgacgaaatgaagaaggccgttgacgatggctggaccaaaaacgccttcaatcaatatgtatcggatctcatatccgttcatcgaactctgcccgatcctcgagatgcctggtgcaaggacgaggcgcgctatctaaccaatctgccaaagaccgatgtcatcatctgcttccacaacgaagcctggaccgtgctgttgagaacagttcactcagtgctggacaggtcgccggaacatctaatagggaagataatcctcgtcgatgactacagcgatatgcctcatttgaagaggcagctggaggactactttgctgcctatcccaaagtccagataattcgtggccaaaagcgagagggtctgatccgagcccgaatttgggtgccaaccatgccaagtcccctgttctcacctatttggattcccactgcgagtgcaccgaaggatggcttgaaccacttttggaccgaattgcacgtaattccaccactgtcgtgtgtcccgttgccgatgtcattagcgatgagaccctggagtaccactatcgcgattcgggcggagttaatgtcggaggattcgattggaatcttcagttcagctggcatccggttcccgaacgtgaaaggaagcgccacaatagcaccgccgaacccgtctactcacccacaatggccggcggagccttctccatcgacagagagttcttcgatcgactgggcacctatgactctggatttgatatttggggcggtgagaatctggagctatccttcaagacctggatgtgcggcggcaccctggagatagttccctgctctcacgtgggacacatattccggaaacgttcgccctacaagtggcgttctggcgtgaatgtgctgaagaagaactccgtgcggttggccgaggtttggatggatgagtactcgcagtactactatcatcgcatcggcaacgacaagggcgactggggcgatgttagcgatcgcaggaagctccgtaacgatctcaagtgcaaaagcttcaaatggtatctggacaacatttatcccgagctcttcattcccggcgattctgtggcccacggagagatcgcgaatgtgcccaacggcatgtgtctggatgcaaaagaaaaatccgaagaggaaacaccggtttccatttatgagtgccatggccaaggaggaaatcagtactggatgctcagcaaggcgggcgaaatccgtcgcgacgactcctgtctcgattatgccggcaaggatgtgacgcttttcggctgccacgggggcaaaggaaatcagttctggacgtaccgcgagaacacaaagcagctgcaccacggcacctccggcaagtgtctggccatcagcgagagcaaggacaagctgctgatggaggagtgcagcgcgagcctcagtcgccaacagtggaccctcgagaactacgacagctcgaagttg |
| WT PGANT35A | atgatgcaaatcaagcgcctcctgtgcaaatcctgcggcctgggcacccttctggtggcggtcgtctggctgctggcgctccttttctactcgcacagcctgcgcagctccatccgctcggcaggatggaggatcgacgaagggaacgcaacgccacgggccgagctgagctaccaggccagggtcaccgtcggatgcactccaaatgccagcactaccactggcgaatcgccggcggcacccaagccgcccagtgatccggagcaactggaactgttgggtgtggtgcgcaacaagcaggacaagtacatacgcgacattggctacaagcatcacgccttcaacgcgctggtgtccaataacataggactcttccgggcgatcccagacactcggcacaaggtgtgcgatcgccaggagaccacggaagcggaaaacctgccccaggcctccatagtcatgtgcttctacaacgagcacaagatgacgctgatgcgttcgattaagacggtgctggaacgcacgcccagctacctgctgcgcgagattattctggtggatgatcacagcgatcttccagaattggaattccatttgcacggggatctgcgtgcccgcctgaagtacgacaacctgcggtacatcaaaaacgagcagcgcgagggtctcattcgttcccgggtcattggtgcacgcgaggccgtgggcgatgtgctcgtcttcctcgactcgcatatcgaggtcaatcagcagtggctggagccgctgctgcggctcatcaagtcggagaacgccactctggccgtgcccgtcatcgacctgatcaacgctgatacctttgagtacacgcccagtccactagtgcgcggaggattcaactggggcctgcacttccgctgggagaatctgcccgagggcacgctcaaggtgccggaggattttcgcggacccttccgttcgcccaccatggccggcggattgtttgccgtgaacaggaagtacttccagcatctgggcgagtacgacatggccatggatatatggggtggcgagaatatagagatctccttcagagcctggcagtgtggtggcgccattaagatcgttccctgctcccgtgtcggtcacatattccgcaaacgacgtccctacacttcgccagatggagccaacaccatgttgaagaactcactgcgtttggcgcacgtctggatggatcagtacaaggattactatctcaagcacgaaaaggtgcccaagacctatgattatggagacattagcgatcgcctcaagcttagggagcgtctgcagtgcagggactttgcctggtatctaaagaatgtctatccagagctgcatgtgccgggcgaggagtccaagaagtcggcggcagcgcccatctttcagccttggcactcgcgaaaacgtaactatgtcgacacattccagctgcgactgacgggaacggagctctgtgccgccgtggtggctcccaaggtgaagggtttctggaagaagggcagctccctgcaattgcaaacctgccgaaggacacccaatcagctgtggtacgaaacggagaaggcggaaattgtgctggacaagttgctgtgcctggaggcatccggggatgcgcaggtcacggtcaacaagtgccacgagatgctcggcgatcagcagtggcggcacactcgcaacgccaacagtcccgtctacaacatggccaaaggcacctgcctccgggcagctgctcccacgacgggcgccctcatcagtctggatctgtgctccaagtccaacggagcgggcggctcctgggacattgtgcagctgaagaagccaacggaggcggagggcagggccaaggaggcgaggaactcggacaaggcgctg |
| BH PGANT35A | atgatgcaaatcaagcgcctcctgtgcaaatcctgcggcctgggcacccttctggtggcggtcgtctggctgctggcgctccttttctactcgcacagcctgcgcagctccatccgctcggcaggatggaggatcgacgaagggaacgcaacgccacgggccgagctgagctaccaggccagggtcaccgtcggatgcactccaaatgccagcactaccactggcgaatcgccggcggcacccaagccgcccagtgatccggagcaactggaactgttgggtgtggtgcgcaacaagcaggacaagtacatacgcgacattggctacaagcatcacgccttcaacgcgctggtgtccaataacataggactcttccgggcgatcccagacactcggcacaaggtgtgcgatcgccaggagaccacggaagcggaaaacctgccccaggcctccatagtcatgtgcttctacaacgagcacaagatgacgctgatgcgttcgattaagacggtgctggaacgcacgcccagctacctgctgcgcgagattattctggtggatgatcacagcgatcttccagaattggaattccatttgcacggggatctgcgtgcccgcctgaagtacgacaacctgcggtacatcaaaaacgagcagcgcgagggtctcattcgttcccgggtcattggtgcacgcgaggccgtgggcgatgtgctcgtcttcctcgactcgcatatcgaggtcaatcagcagtggctggagccgctgctgcggctcatcaagtcggagaacgccactctggccgtgcccgtcgccgacctgatcaacgctgatacctttgagtacacgcccagtccactagtgcgcggaggattcaactggggcctgcacttccgctgggagaatctgcccgagggcacgctcaaggtgccggaggattttcgcggacccttccgttcgcccaccatggccggcggagcctttgccgtgaacaggaagtacttccagcatctgggcgagtacgacatggccatggatatatggggtggcgagaatatagagatctccttcagagcctggcagtgtggtggcgccattaagatcgttccctgctcccgtgtcggtcacatattccgcaaacgacgtccctacacttcgccagatggagccaacaccatgttgaagaactcactgcgtttggcgcacgtctggatggatcagtacaaggattactatctcaagcacgaaaaggtgcccaagacctatgattatggagacattagcgatcgcctcaagcttagggagcgtctgcagtgcagggactttgcctggtatctaaagaatgtctatccagagctgcatgtgccgggcgaggagtccaagaagtcggcggcagcgcccatctttcagccttggcactcgcgaaaacgtaactatgtcgacacattccagctgcgactgacgggaacggagctctgtgccgccgtggtggctcccaaggtgaagggtttctggaagaagggcagctccctgcaattgcaaacctgccgaaggacacccaatcagctgtggtacgaaacggagaaggcggaaattgtgctggacaagttgctgtgcctggaggcatccggggatgcgcaggtcacggtcaacaagtgccacgagatgctcggcgatcagcagtggcggcacactcgcaacgccaacagtcccgtctacaacatggccaaaggcacctgcctccgggcagctgctcccacgacgggcgccctcatcagtctggatctgtgctccaagtccaacggagcgggcggctcctgggacattgtgcagctgaagaagccaacggaggcggagggcagggccaaggaggcgaggaactcggacaaggcgctg |
| UAS-WT-PGANT-9A | tggatttcactggaactagggcgcgcctccggaacataatggtgcagggcgctgacttccgcgtttccagactttacgaaacacggaaaccgaagaccattcatgttgttgctcaggtcgcagacgttttgcagcagcagtcgcttcacgttcgctcgcgtatcggtgattcattctgctaaccagtaaggcaaccccgccagcctagccgggtcctcaacgacaggagcacgatcatgcgcacccgtggccagggccgcaagcttgcatgcctgcaggtcggagtactgtcctccgagcggagtactgtcctccgagcggagtactgtcctccgagcggagtactgtcctccgagcggagtactgtcctccgagcggagactctagccctagggcatgcctgcaggtcggagtactgtcctccgagcggagtactgtcctccgagcggagtactgtcctccgagcggagtactgtcctccgagcggagtactgtcctccgagcggagactctagcgctagcgacgtcgagcgccggagtataaatagaggcgcttcgtctacggagcgacaattcaattcaaacaagcaaagtgaacacgtcgctaagcgaaagctaagcaaataaacaagcgcagctgaacaagctaaacaatctgcagtaaagtgcaagttaaagtgaatcaattaaaagtaaccagcaaccaagtaaatcaactgcaactactgaaatctgccaagaagtaattattgaatacaagaagagaactctgaatagatctaaaaggtaggttcaaccactgatgcctaggcacaccgaaacgactaaccctaattcttatcctttacttcaggcggccgcAACTTAAAAAAAAAAAATCAAAatggccttcatctggcggcgacgttcgacgacgattgtgaagctggtggccttcgccctggccatttggttttgcattgccttcctggtatacacggatgacacccggcgtagagccgcccaggagggggctggtgccagtggggcaggcagtgctcccggagtgggcggcggagccggaggactgggtgatcccattgccttggctctgagaaacgagcctgctggcgaggactttggcatcaatgggaatgtcatcggcggaggaggacagaagcaggcgcacgatgaggccgacatcccacccactgtgggaaaacataaggctgacctacaggcggaaaggatgcggaagaaggcggccgagcagccaaagaaaaagccacaagaagatagcaaaaaagtgattgatccgccggcgaactttgaggagaatcctggcgagctgggaaaacccgtgcgcctgcccaaggaaatgtccgacgaaatgaagaaggccgttgacgatggctggac |

| | |
|---|---|
| | caaaaacgccttcaatcaatatgtatcggatctcatatccgttcatcgaactctgcccgatcctcgagatgcctggtgcaaggacgaggcgcgctatctaaccaatctgccaaagaccgatgtcatcatctgcttccacaacga<br>agcctggaccgtgctgttgagaacagttcactcagtgctggacaggtcgccggaacatctaatagggaagataatcctcgtcgatgactacagcgatatgcctcatttgaagaggcagctggaggactactttgctgcctatcc<br>caaagtccagataattcgtggccaaaagcgagagggtctgatccgagcccgaattttgggtgccaaccatgccaagtcccctgttctcacctatttggattcccactgcgagtgcaccgaaggatggcttgaaccacttttgga<br>ccgaattgcacgtaattccaccactgtcgtgtgtcccgttattgatgtcattagcgatgagaccctggagtaccactatcgcgattcgggcggagttaatgtcggaggattcgattggaatcttcagttcagctggcatccggttccc<br>gaacgtgaaaggaagcgccacaatagcaccgccgaacccgtctactcacccacaatggccggcggacttttctccatcgacagagagttcttcgatcgactgggcacctatgactctggatttgatatttggggcggtgaga<br>atctggagctatccttcaagacctggatgtgcggcggcaccctggagatagttccctgctctcacgtgggacacatattccggaaacgttcgccctacaagtggcgttctggcgtgaatgtgctgaagaagaactccgtgcggtt<br>ggccgaggtttggatggatgagtactcgcagtactactatcatcgcatcggcaacgacaagggcgactggggcgatgttagcgatcgcaggaagctccgtaacgatctcaagtgcaaaagcttcaaatggtatctggacaa<br>catttatcccgagctcttcattcccggcgattctgtggcccacggagagataagaaatctaggttatggcggtcgcacctgcttggacgcaccggctggcaagaagcaccagaagaaggcggtgggcacgtatccctgccat<br>cggcagggtggaaaccagtactggatgctcagcaaggcgggcgaaatccgtcgcgacgactcctgtctcgattatgccggcaaggatgtgacgcttttcggctgccacgggggcaaaggaaatcagttctggacgtaccg<br>cgagaacacaaagcagctgcaccacggcacctccggcaagtgtctggccatcagcgagagcaaggacaagctgctgatggaggagtgcagcgcgagcctcagtcgccaacagtggaccctcgagaactacgacag<br>ctcgaagttgTACACTGATATCGAAATGAACCGCCTGGGTAAGTAGtctagaatgaatcgtttttaaaataacaaatcaattgttttataatattcgtacgattctttgattatgtaataaaatgtgatcattagg<br>aagattacgaaaaatataaaaaatatgagttctgtgtgtataacaaatgctgtaaacgccacaattgtgtttgttgcaaataaacccatgattatttgattaaaattgttgttttctttgttcatagacaatagtgtgttttgcctaaacgtgt<br>actgcataaactccatgcgagtgtatagcgagctagtggctaacgcttgccccaccaaagtagattcgtcaaaatcctcaatttcatcaccctcctccaagtttaacatttggccgtcggaattaacttctaaagatgccacataat<br>ctaataaatgaaatagagattcaaacgtggcgtcatcgtccgtttcgaccatttccgaaaagaactcgggcataaactctatgatttctctggacgtggtgttgtcgaaactctcaaagtacgcagtcaggaacgtgcgcgacat<br>gtcgtcgggaaactcgcgcggaaacatgttgttgtaaccgaacgggtcccatagcgccaaaaccaaatctgccagcgtcaatagaatgagcacgatgccgacaatggagctggcttggatagcgattcgagttaacggcc<br>ggccgtttaaacgaattcttgaagacgaaagggcctcgtgatacgcctatttttataggttaatgtcatgataataatggtttcttagactcaggtggcacttttcggggaaatgtgcgcggaacccctatttgtttatttttctaaatacat<br>tcaaatatgtatccgctcatgagacaataaccctgataaatgcttcaataatattgaaaaaggaagagtatgagtattcaacatttccgtgtcgcccttattcccttttttgcggcattttgccttcctgtttttgctcacccagaaacgct<br>ggtgaaagtaaaagatgctgaagatcagttgggtgcacgagtgggttacatcgaactggatctcaacagcggtaagatccttgagagttttcgccccgaagaacgttttccaatgatgagcacttttaaagttctgctatgtggcg<br>cggtattatcccgtattgacgccgggcaagagcaactcggtcgccgcatacactattctcagaatgacttggttgagtactcaccagtcacagaaaagcatcttacggatggcatgacagtaagagaattatgcagtgctgcc<br>ataaccatgagtgataacactgcggccaacttacttctgacaacgatcggaggaccgaaggagctaaccgcttttttgcacaacatgggggatcatgtaactcgccttgatcgttgggaaccggagctgaatgaagccatac<br>caaacgacgagcgtgacaccacgatgcctgtagcaatggcaacaacgttgcgcaaactattaactggcgaactacttactctagcttcccggcaacaattaatagactggatggaggcggataaagttgcaggaccacttct<br>gcgctcggcccttccggctggctggtttattgctgataaatctggagccggtgagcgtgggtctcgcggtatcattgcagcactggggccagatggtaagccctcccgtatcgtagttatctacacgacggggagtcaggcaact<br>atggatgaacgaaatagacagatcgctgagataggtgcctcactgattaagcattggtaactgtcagaccaagtttactcatatatactttagattgatttaaaacttcatttttaatttaaaaggatctaggtgaagatcctttttgata<br>atctcatgaccaaaatcccttaacgtgagttttcgttccactgagcgtcagaccccgtagaaaagatcaaaggatcttcttgagatcctttttttctgcgcgtaatctgctgcttgcaaacaaaaaaaccaccgctaccagcggtgg<br>tttgtttgccggatcaagagctaccaactctttttccgaaggtaactggcttcagcagagcgcagataccaaatactgttcttctagtgtagccgtagttaggccaccacttcaagaactctgtagcaccgcctacatacctcgctct<br>gctaatcctgttaccagtggctgctgccagtggcgataagtcgtgtcttaccgggttggactcaagacgatagttaccggataaggcgcagcggtcgggctgaacggggggttcgtgcacacagcccagcttggagcgaac<br>gacctacaccgaactgagatacctacagcgtgagctatgagaaagcgccacgcttcccgaagggagaaaggcggacaggtatccggtaagcggcagggtcggaacaggagagcgcacgagggagcttccagggg<br>gaaacgcctggtatctttatagtcctgtcgggtttcgccacctctgacttgagcgtcgatttttgtgatgctcgtcaggggggcggagcctatggaaaaacgccagcaacgcggcctttttacggttcctggccttttgctggccttttg<br>ctcacatgttaccgtcgacgatgtaggtcacggtctcgaagccgcggtgcgggtgccagggcgtgcccttgggctccccgggcgcgtactccacctcacccatctggtccatcatgatgaacgggtcgaggtggcggtagttg<br>atcccggcgaacgcgcggcgcaccgggaagccctcgccctcgaaaccgctgggcgcggtggtcacggtgagcacgggacgtgcgacggcgtcggcgggtgcggatacgcggggcagcgtcagcgggttctcgacg<br>gtcacggcgggcatgtcgacaagccgaacatatgggcgcgcctagtatgtatgtaagttaataaaacccatttttgcggaaagtagataaaaaaaacatttttttttactgcactggatatcattgaacttatctgatcagttttaa<br>atttacttcgatccaagggtatttgatgtaccaggttctttcgattacctctcactcaaaatgacattccactcaaagtcagcgctgtttgcctccttctctgtccacagaaatatcgccgtctctttcgccgctgcgtccgctatctctttcg<br>ccaccgtttgtagcgttacgtagcgtcaatgtccgccttcagttgcattttgtcagcggtttcgtgacgaagctccaagcggtttacgccatcaattaaacacaaagtgctgtgccaaaactcctctcgcttcttatttttgtttgtttttga<br>gtgattggggtggtgattggttttgggtgggtaagcaggggaaagtgtgaaaaatcccggcaatgggccaagaggatcaggagctattaattcgcggaggcagcaaacacccatctgccgagcatctgaacaatgtgagta<br>gtacatgtgcatacatcttaaggaagttatgcccggatctaattcaattagagactaattcaattagagctaattcaattagcgccggagtataaatagaggcgcttcgtctacggagcgacaattcaattcaaacaagcaaagt<br>gaacacgtcgctaagcgaaagctaagcaaataaacaagcgcagctgaacaagctaaacaatcggcccatggtgagcaagggcgaggaggataacatggccatcatcaaggagttcatgcgcttcaaggtgcacatgg<br>agggctccgtgaacggccacgagttcgagatcgagggcgagggcgagggccgcccctacgagggcacccagaccgccaagctgaaggtgaccaagggtggcccccctgcccttcgcctgggacatcctgtcccctcag<br>ttcatgtacggctccaaggcctacgtgaagcaccccgccgacatccccgactacttgaagctgtccttccccgagggcttcaagtgggagcgcgtgatgaacttcgaggacggcggcgtggtgaccgtgacccaggactcct<br>ccctgcaggacggcgagttcatctacaaggtgaagctgcgcggcaccaacttcccctccgacggccccgtaatgcagaagaagaccatgggctgggaggcctcctccgagcggatgtaccccgaggacggcgccctga<br>agggcgagatcaagcagaggctgaagctgaaggacggcggccactacgacgctgaggtcaagaccacctacaaggccaagaagcccgtgcagctgcccggcgcctacaacgtcaacatcaagttggacatcacctc<br>ccacaacgaggactacaccatcgtggaacagtacgaacgcgccgagggccgccactccaccggcggcatggacgagctgtacaagtaagcgccgcgactctgatcataatcagccataccacatttgtagaggttttact<br>tgctttaaaaaacctcccacacctccccctgaacctgaaacataaaatgaatgcaattgttgttgttaacttgtttattgcagcttataatggttacaaataaagcaatagcatcacaaatttcacaaataaagcatttttttcactgcat<br>tctagttgtggtttgtccaaactcatcaatgtatcttagggcccgacgcaaggagtagccgacatatatccgaaataactgcttgtttttttttaccattattaccatcgtgtttactgtttattgccccctcaaaaagctaatgtaattatat<br>ttgtgccaataaaaacaagatatgacctatagaatacaagtatttccccttcgaacatccccacaagtagactttggatttgtcttctaaccaaaagacttacacacctgcataccttacatcaaaaactcgtttatcgctacataaa<br>acaccgggatatatttttatatacatacttttcaaatcgcgcgccctcttcataattcacctccaccacaccacgtttcgtagttgctctttcgctgtctcccacccgctctccgcaacacattcaccttttgttcgacgaccttggagcga<br>ctgtcgttagttccgcgcgattcggttcgctcaaatggttccgagtggttcatttcgtctcaatagaaattagtaataaatatttgtatgtacaatttatttgctccaatatatttgtatatatttccctcacagctatatttattctaatttaatatta<br>tgactttttaaggtaatttttttgtgacctgttcggagtgattagcgttacaatttgaactgaaagtgacatccagtgtttgttccttgtgtagatgcatctcaaaaaaatggtgggcataatagtgttgtttatatatatcaaaaataacaact<br>ataataataagaatacatttaatttagaaaatgct |
| UAS-BH-PGANT-9A | tggatttcactggaactagggcgcgcctccggaacataatggtgcagggcgctgacttccgcgtttccagactttacgaaacacggaaaccgaagaccattcatgttgttgctcaggtcgcagacgttttgcagcagcagtcgc<br>ttcacgttcgctcgcgtatcggtgattcattctgctaaccagtaaggcaaccccgccagcctagccgggtcctcaacgacaggagcacgatcatgcgcacccgtggccagggccgcaagcttgcatgcctgcaggtcggagt<br>actgtcctccgagcggagtactgtcctccgagcggagtactgtcctccgagcggagtactgtcctccgagcggagtactgtcctccgagcggagactctagccctagggcatgcctgcaggtcggagtactgtcctccgagcg<br>gagtactgtcctccgagcggagtactgtcctccgagcggagtactgtcctccgagcggagtactgtcctccgagcggagactctagcgctagcgacgtcgagcgccggagtataaatagaggcgcttcgtctacggagcga<br>caattcaattcaaacaagcaaagtgaacacgtcgctaagcgaaagctaagcaaataaacaagcgcagctgaacaagctaaacaatctgcagtaaagtgcaagttaaagtgaatcaattaaaagtaaccagcaaccaa<br>gtaaatcaactgcaactactgaaatctgccaagaagtaattattgaatacaagaagagaactctgaatagatctaaaaggtaggttcaaccactgatgcctaggcacaccgaaacgactaaccctaattcttatcctttacttca<br>ggcggccgcAACTTAAAAAAAAAAATCAAAatggccttcatctggcggcgacgttcgacgacgattgtgaagctggtggccttcgccctggccatttggttttgcattgccttcctggtatacacggatgacacccgg<br>cgtagagccgcccaggagggggctggtgccagtggggcaggcagtgctcccggagtgggcggcggagccggaggactgggtgatcccattgccttggctctgagaaacgagcctgctggcgaggactttggcatcaat<br>gggaatgtcatcggcggaggaggacagaagcaggcgcacgatgaggccgacatcccacccactgtgggaaaacataaggctgacctacaggcggaaaggatgcggaagaaggcggccgagcagccaaagaaa<br>aagccacaagaagatagcaaaaaagtgattgatccgccggcgaactttgaggagaatcctggcgagctgggaaaacccgtgcgcctgcccaaggaaatgtccgacgaaatgaagaaggccgttgacgatggctggac<br>caaaaacgccttcaatcaatatgtatcggatctcatatccgttcatcgaactctgcccgatcctcgagatgcctggtgcaaggacgaggcgcgctatctaaccaatctgccaaagaccgatgtcatcatctgcttccacaacga<br>agcctggaccgtgctgttgagaacagttcactcagtgctggacaggtcgccggaacatctaatagggaagataatcctcgtcgatgactacagcgatatgcctcatttgaagaggcagctggaggactactttgctgcctatcc<br>caaagtccagataattcgtggccaaaagcgagagggtctgatccgagcccgaattttgggtgccaaccatgccaagtcccctgttctcacctatttggattcccactgcgagtgcaccgaaggatggcttgaaccacttttgga<br>ccgaattgcacgtaattccaccactgtcgtgtgtcccgttgccgatgtcattagcgatgagaccctggagtaccactatcgcgattcgggcggagttaatgtcggaggattcgattggaatcttcagttcagctggcatccggttcc<br>cgaacgtgaaaggaagcgccacaatagcaccgccgaacccgtctactcacccacaatggccggcggagccttctccatcgacagagagttcttcgatcgactgggcacctatgactctggatttgatatttggggcggtgag<br>aatctggagctatccttcaagacctggatgtgcggcggcaccctggagatagttccctgctctcacgtgggacacatattccggaaacgttcgccctacaagtggcgttctggcgtgaatgtgctgaagaagaactccgtgcgg<br>ttggccgaggtttggatggatgagtactcgcagtactactatcatcgcatcggcaacgacaagggcgactggggcgatgttagcgatcgcaggaagctccgtaacgatctcaagtgcaaaagcttcaaatggtatctggaca<br>acatttatcccgagctcttcattcccggcgattctgtggcccacggagagataagaaatctaggttatggcggtcgcacctgcttggacgcaccggctggcaagaagcaccagaagaaggcggtgggcacgtatccctgcc<br>atcggcagggtggaaaccagtactggatgctcagcaaggcgggcgaaatccgtcgcgacgactcctgtctcgattatgccggcaaggatgtgacgcttttcggctgccacgggggcaaaggaaatcagttctggacgtacc<br>gcgagaacacaaagcagctgcaccacggcacctccggcaagtgtctggccatcagcgagagcaaggacaagctgctgatggaggagtgcagcgcgagcctcagtcgccaacagtggaccctcgagaactacgaca<br>gctcgaagttgTACACTGATATCGAAATGAACCGCCTGGGTAAGTAGtctagaatgaatcgtttttaaaataacaaatcaattgttttataatattcgtacgattctttgattatgtaataaaatgtgatcattag<br>gaagattacgaaaaatataaaaaatatgagttctgtgtgtataacaaatgctgtaaacgccacaattgtgtttgttgcaaataaacccatgattatttgattaaaattgttgttttctttgttcatagacaatagtgtgttttgcctaaacgt<br>gtactgcataaactccatgcgagtgtatagcgagctagtggctaacgcttgccccaccaaagtagattcgtcaaaatcctcaatttcatcaccctcctccaagtttaacatttggccgtcggaattaacttctaaagatgccacata<br>atctaataaatgaaatagagattcaaacgtggcgtcatcgtccgtttcgaccatttccgaaaagaactcgggcataaactctatgatttctctggacgtggtgttgtcgaaactctcaaagtacgcagtcaggaacgtgcgcgac<br>atgtcgtcgggaaactcgcgcggaaacatgttgttgtaaccgaacgggtcccatagcgccaaaaccaaatctgccagcgtcaatagaatgagcacgatgccgacaatggagctggcttggatagcgattcgagttaacgg<br>ccggccgtttaaacgaattcttgaagacgaaagggcctcgtgatacgcctatttttataggttaatgtcatgataataatggtttcttagactcaggtggcacttttcggggaaatgtgcgcggaacccctatttgtttatttttctaaata<br>cattcaaatatgtatccgctcatgagacaataaccctgataaatgcttcaataatattgaaaaaggaagagtatgagtattcaacatttccgtgtcgcccttattcccttttttgcggcattttgccttcctgtttttgctcacccagaaac<br>gctggtgaaagtaaaagatgctgaagatcagttgggtgcacgagtgggttacatcgaactggatctcaacagcggtaagatccttgagagttttcgccccgaagaacgttttccaatgatgagcacttttaaagttctgctatgtg<br>gcgcggtattatcccgtattgacgccgggcaagagcaactcggtcgccgcatacactattctcagaatgacttggttgagtactcaccagtcacagaaaagcatcttacggatggcatgacagtaagagaattatgcagtgct<br>gccataaccatgagtgataacactgcggccaacttacttctgacaacgatcggaggaccgaaggagctaaccgcttttttgcacaacatgggggatcatgtaactcgccttgatcgttgggaaccggagctgaatgaagcca<br>taccaaacgacgagcgtgacaccacgatgcctgtagcaatggcaacaacgttgcgcaaactattaactggcgaactacttactctagcttcccggcaacaattaatagactggatggaggcggataaagttgcaggaccac<br>ttctgcgctcggcccttccggctggctggtttattgctgataaatctggagccggtgagcgtgggtctcgcggtatcattgcagcactggggccagatggtaagccctcccgtatcgtagttatctacacgacggggagtcaggca<br>actatggatgaacgaaatagacagatcgctgagataggtgcctcactgattaagcattggtaactgtcagaccaagtttactcatatatactttagattgatttaaaacttcatttttaatttaaaaggatctaggtgaagatcctttttg<br>ataatctcatgaccaaaatcccttaacgtgagttttcgttccactgagcgtcagaccccgtagaaaagatcaaaggatcttcttgagatcctttttttctgcgcgtaatctgctgcttgcaaacaaaaaaaccaccgctaccagcgg<br>tggtttgtttgccggatcaagagctaccaactctttttccgaaggtaactggcttcagcagagcgcagataccaaatactgttcttctagtgtagccgtagttaggccaccacttcaagaactctgtagcaccgcctacatacctcgc<br>tctgctaatcctgttaccagtggctgctgccagtggcgataagtcgtgtcttaccgggttggactcaagacgatagttaccggataaggcgcagcggtcgggctgaacggggggttcgtgcacacagcccagcttggagcga<br>acgacctacaccgaactgagatacctacagcgtgagctatgagaaagcgccacgcttcccgaagggagaaaggcggacaggtatccggtaagcggcagggtcggaacaggagagcgcacgagggagcttccagg<br>gggaaacgcctggtatctttatagtcctgtcgggtttcgccacctctgacttgagcgtcgatttttgtgatgctcgtcaggggggcggagcctatggaaaaacgccagcaacgcggcctttttacggttcctggccttttgctggccttt<br>tgctcacatgttaccgtcgacgatgtaggtcacggtctcgaagccgcggtgcgggtgccagggcgtgcccttgggctccccgggcgcgtactccacctcacccatctggtccatcatgatgaacgggtcgaggtggcggtagt<br>tgatcccggcgaacgcgcggcgcaccgggaagccctcgccctcgaaaccgctgggcgcggtggtcacggtgagcacgggacgtgcgacggcgtcggcgggtgcggatacgcggggcagcgtcagcgggttctcgac<br>ggtcacggcgggcatgtcgacaagccgaacatatgggcgcgcctagtatgtatgtaagttaataaaacccatttttgcggaaagtagataaaaaaaacatttttttttactgcactggatatcattgaacttatctgatcagtttta<br>aatttacttcgatccaagggtatttgatgtaccaggttctttcgattacctctcactcaaaatgacattccactcaaagtcagcgctgtttgcctccttctctgtccacagaaatatcgccgtctctttcgccgctgcgtccgctatctctttc<br>gccaccgtttgtagcgttacgtagcgtcaatgtccgccttcagttgcattttgtcagcggtttcgtgacgaagctccaagcggtttacgccatcaattaaacacaaagtgctgtgccaaaactcctctcgcttcttatttttgtttgtttttg<br>agtgattggggtggtgattggttttgggtgggtaagcaggggaaagtgtgaaaaatcccggcaatgggccaagaggatcaggagctattaattcgcggaggcagcaaacacccatctgccgagcatctgaacaatgtgagt<br>agtacatgtgcatacatcttaaggaagttatgcccggatctaattcaattagagactaattcaattagagctaattcaattagcgccggagtataaatagaggcgcttcgtctacggagcgacaattcaattcaaacaagcaaag<br>tgaacacgtcgctaagcgaaagctaagcaaataaacaagcgcagctgaacaagctaaacaatcggcccatggtgagcaagggcgaggaggataacatggccatcatcaaggagttcatgcgcttcaaggtgcacatg<br>gagggctccgtgaacggccacgagttcgagatcgagggcgagggcgagggccgcccctacgagggcacccagaccgccaagctgaaggtgaccaagggtggcccccctgcccttcgcctgggacatcctgtcccctca<br>gttcatgtacggctccaaggcctacgtgaagcaccccgccgacatccccgactacttgaagctgtccttccccgagggcttcaagtgggagcgcgtgatgaacttcgaggacggcggcgtggtgaccgtgacccaggactc<br>ctccctgcaggacggcgagttcatctacaaggtgaagctgcgcggcaccaacttcccctccgacggccccgtaatgcagaagaagaccatgggctgggaggcctcctccgagcggatgtaccccgaggacggcgccct<br>gaagggcgagatcaagcagaggctgaagctgaaggacggcggccactacgacgctgaggtcaagaccacctacaaggccaagaagcccgtgcagctgcccggcgcctacaacgtcaacatcaagttggacatcac<br>ctcccacaacgaggactacaccatcgtggaacagtacgaacgcgccgagggccgccactccaccggcggcatggacgagctgtacaagtaagcgccgcgactctgatcataatcagccataccacatttgtagaggtttt<br>acttgctttaaaaaacctcccacacctccccctgaacctgaaacataaaatgaatgcaattgttgttgttaacttgtttattgcagcttataatggttacaaataaagcaatagcatcacaaatttcacaaataaagcatttttttcactg<br>cattctagttgtggtttgtccaaactcatcaatgtatcttagggcccgacgcaaggagtagccgacatatatccgaaataactgcttgtttttttttaccattattaccatcgtgtttactgtttattgccccctcaaaaagctaatgtaatt<br>atatttgtgccaataaaaacaagatatgacctatagaatacaagtatttccccttcgaacatccccacaagtagactttggatttgtcttctaaccaaaagacttacacacctgcataccttacatcaaaaactcgtttatcgctacat<br>aaaacaccgggatatatttttatatacatacttttcaaatcgcgcgccctcttcataattcacctccaccacaccacgtttcgtagttgctctttcgctgtctcccacccgctctccgcaacacattcaccttttgttcgacgaccttggag<br>cgactgtcgttagttccgcgcgattcggttcgctcaaatggttccgagtggttcatttcgtctcaatagaaattagtaataaatatttgtatgtacaatttatttgctccaatatatttgtatatatttccctcacagctatatttattctaatttaat<br>attatgactttttaaggtaatttttttgtgacctgttcggagtgattagcgttacaatttgaactgaaagtgacatccagtgtttgttccttgtgtagatgcatctcaaaaaaatggtgggcataatagtgttgtttatatatatcaaaaataac<br>aactataataataagaatacatttaatttagaaaatgct |

| | |
|---|---|
| UAS-WT-PGANT-35A | tggatttcactggaactagggcgcgcctccggaacataatggtgcagggcgctgacttccgcgtttccagactttacgaaacacggaaaccgaagaccattcatgttgttgctcaggtcgcagacgttttgcagcagcagtcgc<br>ttcacgttcgctcgcgtatcggtgattcattctgctaaccagtaaggcaaccccgccagcctagccgggtcctcaacgacaggagcacgatcatgcgcacccgtggccagggccgcaagcttgcatgcctgcaggtcggagt<br>actgtcctccgagcggagtactgtcctccgagcggagtactgtcctccgagcggagtactgtcctccgagcggagtactgtcctccgagcggagactctagccctagggcatgcctgcaggtcggagtactgtcctccgagcg<br>gagtactgtcctccgagcggagtactgtcctccgagcggagtactgtcctccgagcggagtactgtcctccgagcggagactctagcgctagcgacgtcgagcgccggagtataaatagaggcgcttcgtctacggagcga<br>caattcaattcaaacaagcaaagtgaacacgtcgctaagcgaaagctaagcaaataaacaagcgcagctgaacaagctaaacaatctgcagtaaagtgcaagttaaagtgaatcaattaaaagtaaccagcaaccaa<br>gtaaatcaactgcaactactgaaatctgccaagaagtaattattgaatacaagaagagaactctgaatagatctaaaaggtaggttcaaccactgatgcctaggcacaccgaaacgactaaccctaattcttatcctttacttca<br>ggcggccgcAACTTAAAAAAAAAAATCAAAatgatgcaatcaagcgcctcctgtgcaaatcctgcggcctgggcacccttctggtggcggtcgtctggctgctggcgctccttttctactcgcacagcctgcgca<br>gctccatccgctcggcaggatggaggatcgacgaagggaacgcaacgccacgggccgagctgagctaccaggccagggtcaccgtcggatgcactccaaatgccagcactaccactggcgaatcgccggcggcacc<br>caagccgcccagtgatccggagcaactggaactgttgggtgtggtgcgcaacaagcaggacaagtacatacgcgacattggctacaagcatcacgccttcaacgcgctggtgtccaataacataggactcttccgggcga<br>tcccagacactcggcacaaggtgtgcgatcgccaggagaccacggaagcggaaaacctgccccaggcctccatagtcatgtgcttctacaacgagcacaagatgacgctgatgcgttcgattaagacggtgctggaacg<br>cacgcccagctacctgctgcgcgagattattctggtggatgatcacagcgatcttccagaattggaattccatttgcacggggatctgcgtgcccgcctgaagtacgacaacctgcggtacatcaaaaacgagcagcgcgag<br>ggtctcattcgttcccgggtcattggtgcacgcgaggccgtgggcgatgtgctcgtcttcctcgactcgcatatcgaggtcaatcagcagtggctggagccgctgctgcggctcatcaagtcggagaacgccactctggccgtg<br>cccgtcatcgacctgatcaacgctgatacctttgagtacacgcccagtccactagtgcgcggaggattcaactggggcctgcacttccgctgggagaatctgcccgagggcacgctcaaggtgccggaggattttcgcggac<br>ccttccgttcgcccaccatggccggcggattgtttgccgtgaacaggaagtacttccagcatctgggcgagtacgacatggccatggatatatggggtggcgagaatatagagatctccttcagagcctggcagtgtggtggc<br>gccattaagatcgttccctgctcccgtgtcggtcacatattccgcaaacgacgtccctacacttcgccagatggagccaacaccatgttgaagaactcactgcgtttggcgcacgtctggatggatcagtacaaggattactatct<br>caagcacgaaaaggtgcccaagacctatgattatggagacattagcgatcgcctcaagcttagggagcgtctgcagtgcagggactttgcctggtatctaaagaatgtctatccagagctgcatgtgccgggcgaggagtcc<br>aagaagtcggcggcagcgcccatctttcagccttggcactcgcgaaaacgtaactatgtcgacacattccagctgcgactgacgggaacggagctctgtgccgccgtggtggctcccaaggtgaagggtttctggaagaag<br>ggcagctccctgcaattgcaaacctgccgaaggacacccaatcagctgtggtacgaaacggagaaggcggaaattgtgctggacaagttgctgtgcctggaggcatccggggatgcgcaggtcacggtcaacaagtgcc<br>acgagatgctcggcgatcagcagtggcggcacactcgcaacgccaacagtcccgtctacaacatggccaaaggcacctgcctccgggcagctgctcccacgacgggcgccctcatcagtctggatctgtgctccaagtcc<br>aacggagcgggcggctcctgggacattgtgcagctgaagaagccaacggaggcggagggcagggccaaggaggcgaggaactcggacaaggcgctgTACACTGATATCGAAATGAACCGCCTG<br>GGTAAGTAGtctagaatgaatcgttttaaaataacaaatcaattgttttatatattcgtacgattctttgattatgtaataaaatgtgatcattaggaagattacgaaaaatataaaaaatatgagttctgtgtgtataacaaatg<br>ctgtaaacgccacaattgtgtttgttgcaaataaacccatgattatttgattaaaattgttgttttcttgttcatagacaatagtgtgttttgcctaaacgtgtactgcataaactccatgcgagtgtatagcgagctagtggctaacgcttg<br>ccccaccaaagtagattcgtcaaaatcctcaatttcatcaccctcctccaagtttaacatttggccgtcggaattaacttctaaagatgccacataatctaataaatgaaatagagattcaaacgtggcgtcatcgtccgtttcgac<br>catttccgaaaagaactcgggcataaactctatgatttctctggacgtggtgttgtcgaaactctcaaagtacgcagtcaggaacgtgcgcgacatgtcgtcgggaaactcgcgcggaaacatgttgttgtaaccgaacgggtc<br>ccatagcgccaaaaccaaatctgccagcgtcaatagaatgagcacgatgccgacaatggagctggcttggatagcgattcgagttaacggccggccgtttaaacgaattcttgaagacgaaaggcctcgtgatacgcct<br>atttttataggttaatgtcatgataataatggtttcttagactcaggtggcacttttcggggaaatgtgcgcggaacccctatttgtttatttttctaaatacattcaaatatgtatccgctcatgagacaataaccctgataaatgcttcaat<br>aatattgaaaaaggaagagtatgagtattcaacatttccgtgtcgcccttattcccttttttgcggcattttgccttcctgtttttgctcacccagaaacgctggtgaaagtaaaagatgctgaagatcagttgggtgcacgagtgggtt<br>acatcgaactggatctcaacagcggtaagatccttgagagttttcgccccgaagaacgttttccaatgatgagcacttttaaagttctgctatgtggcgcggtattatcccgtattgacgccgggcaagagcaactcggtcgccgc<br>atacactattctcagaatgacttggttgagtactcaccagtcacagaaaagcatcttacggatggcatgacagtaagagaattatgcagtgctgccataaccatgagtgataacactgcggccaacttacttctgacaacgatc<br>ggaggaccgaaggagctaaccgcttttttgcacaacatgggggatcatgtaactcgccttgatcgttgggaaccggagctgaatgaagccataccaaacgacgagcgtgacaccacgatgcctgtagcaatggcaacaa<br>cgttgcgcaaactattaactggcgaactacttactctagcttcccggcaacaattaatagactggatggaggcggataaagttgcaggaccacttctgcgctcggcccttccggctggctggtttattgctgataaatctggagcc<br>ggtgagcgtgggtctcgcggtatcattgcagcactggggccagatggtaagccctcccgtatcgtagttatctacacgacggggagtcaggcaactatggatgaacgaaatagacagatcgctgagataggtgcctcactga<br>ttaagcattggtaactgtcagaccaagtttactcatatatactttagattgatttaaaacttcatttttaatttaaaaggatctaggtgaagatcctttttgataatctcatgaccaaaatcccttaacgtgagttttcgttccactgagcgtca<br>gaccccgtagaaaagatcaaaggatcttcttgagatcctttttttctgcgcgtaatctgctgcttgcaaacaaaaaaaccaccgctaccagcggtggtttgtttgccggatcaagagctaccaactctttttccgaaggtaactggct<br>tcagcagagcgcagataccaaatactgttcttctagtgtagccgtagttaggccaccacttcaagaactctgtagcaccgcctacatacctcgctctgctaatcctgttaccagtggctgctgccagtggcgataagtcgtgtctta<br>ccgggttggactcaagacgatagttaccggataaggcgcagcggtcgggctgaacggggggttcgtgcacacagcccagcttggagcgaacgacctacaccgaactgagatacctacagcgtgagctatgagaaagc<br>gccacgcttcccgaagggagaaaggcggacaggtatccggtaagcggcagggtcggaacaggagagcgcacgagggagcttccagggggaaacgcctggtatctttatagtcctgtcgggtttcgccacctctgacttg<br>agcgtcgatttttgtgatgctcgtcaggggggcggagcctatggaaaaacgccagcaacgcggcctttttacggttcctggccttttgctggccttttgctcacatgttaccgtcgacgatgtaggtcacggtctcgaagccgcggt<br>gcgggtgccagggcgtgcccttgggctccccgggcgcgtactccacctcacccatctggtccatcatgatgaacgggtcgaggtggcggtagttgatcccggcgaacgcgcggcgcaccgggaagccctcgccctcgaa<br>accgctgggcgcggtggtcacggtgagcacgggacgtgcgacggcgtcggcgggtgcggatacgcggggcagcgtcagcgggttctcgacggtcacggcgggcatgtcgacaagccgaacatatggcgcgcctagt<br>atgtatgtaagttaataaaacccattttgcggaaagtagataaaaaaaacatttttttttactgcactggatatcattgaacttatctgatcagttttaaatttacttcgatccaagggtatttgatgtaccaggttctttcgattacctctc<br>actcaaaatgacattccactcaaagtcagcgctgtttgcctccttctctgtccacagaaatatcgccgtctctttcgccgctgcgtccgctatctctttcgccaccgtttgtagcgttacgtagcgtcaatgtccgccttcagttgcatttg<br>tcagcggtttcgtgacgaagctccaagcggtttacgccatcaattaaacacaaagtgctgtgccaaaactcctctcgcttcttatttttgtttgtttttgagtgattggggtggtgattggttttgggtgggtaagcaggggaaagtgtg<br>aaaaatcccggcaatgggccaagaggatcaggagctattaattcgcggaggcagcaaacacccatctgccgagcatctgaacaatgtgagtagtacatgtgcatacatcttaaggaagttatgcccggatctaattcaatta<br>gagactaattcaattagagctaattcaattagcgccggagtataaatagaggcgcttcgtctacggagcgacaattcaattcaaacaagcaaagtgaacacgtcgctaagcgaaagctaagcaaataaacaagcgcagct<br>gaacaagctaaacaatcggcccatggtgagcaagggcgaggaggataacatggccatcatcaaggagttcatgcgcttcaaggtgcacatggagggctccgtgaacggccacgagttcgagatcgagggcgagggcg<br>agggccgcccctacgagggcacccagaccgccaagctgaaggtgaccaagggtggcccctgcccttcgcctgggacatcctgtcccctcagttcatgtacggctccaaggcctacgtgaagcaccccgccgacatccc<br>cgactacttgaagctgtccttccccgagggcttcaagtgggagcgcgtgatgaacttcgaggacggcggcgtggtgaccgtgacccaggactcctccctgcaggacggcgagttcatctacaaggtgaagctgcgcggcac<br>caacttcccctccgacggccccgtaatgcagaagaagaccatgggctgggaggcctcctccgagcggatgtaccccgaggacggcgccctgaagggcgagatcaagcagaggctgaagctgaaggacggcggcca<br>ctacgacgctgaggtcaagaccacctacaaggccaagaagcccgtgcagctgcccggcgcctacaacgtcaacatcaagttggacatcacctcccacaacgaggactacaccatcgtggaacagtacgaacgcgccg<br>agggccgccactccaccggcggcatggacgagctgtacaagtaagcgccgcgactctgatcataatcagccataccacatttgtagaggttttacttgctttaaaaaacctcccacacctccccctgaacctgaaacataaa<br>atgaatgcaattgttgttgttaacttgtttattgcagcttataatggttacaaataaagcaatagcatcacaaatttcacaaataaagcatttttttcactgcattctagttgtggtttgtccaaactcatcaatgtatcttagggcccgacgc<br>aaggagtagccgacatatatccgaaataactgcttgtttttttttaccattattaccatcgtgtttactgtttattgccccctcaaaaagctaatgtaattatatttgtgccaataaaaacaagatatgacctatagaatacaagtatttcc<br>ccttcgaacatccccacaagtagactttggatttgtcttctaaccaaaagacttacacacctgcataccttacatcaaaaactcgtttatcgctacataaaacaccgggatatatttttatatacatacttttcaaatcgcgcgccctct<br>tcataattcacctccaccacaccacgtttcgtagttgctctttcgctgtctcccacccgctctccgcaacacattcaccttttgttcgacgaccttggagcgactgtcgttagttccgcgcgattcggttcgctcaaatggttccgagtggt<br>tcatttcgtctcaatagaaattagtaataaatatttgtatgtacaatttatttgctccaatatatttgtatatatttccctcacagctatatttattctaatttaatattatgactttttaaggtaatttttttgtgacctgttcggagtgattagcgttaca<br>atttgaactgaaagtgacatccagtgtttgttccttgtgtagatgcatctcaaaaaaatggtgggcataatagtgttgtttatatatatcaaaaataacaactataataataagaatacatttaatttagaaaatgct |
| UAS-BH-PGANT-35A | tggatttcactggaactagggcgcgcctccggaacataatggtgcagggcgctgacttccgcgtttccagactttacgaaacacggaaaccgaagaccattcatgttgttgctcaggtcgcagacgttttgcagcagcagtcgc<br>ttcacgttcgctcgcgtatcggtgattcattctgctaaccagtaaggcaaccccgccagcctagccgggtcctcaacgacaggagcacgatcatgcgcacccgtggccagggccgcaagcttgcatgcctgcaggtcggagt<br>actgtcctccgagcggagtactgtcctccgagcggagtactgtcctccgagcggagtactgtcctccgagcggagtactgtcctccgagcggagactctagccctagggcatgcctgcaggtcggagtactgtcctccgagcg<br>gagtactgtcctccgagcggagtactgtcctccgagcggagtactgtcctccgagcggagtactgtcctccgagcggagactctagcgctagcgacgtcgagcgccggagtataaatagaggcgcttcgtctacggagcga<br>caattcaattcaaacaagcaaagtgaacacgtcgctaagcgaaagctaagcaaataaacaagcgcagctgaacaagctaaacaatctgcagtaaagtgcaagttaaagtgaatcaattaaaagtaaccagcaaccaa<br>gtaaatcaactgcaactactgaaatctgccaagaagtaattattgaatacaagaagagaactctgaatagatctaaaaggtaggttcaaccactgatgcctaggcacaccgaaacgactaaccctaattcttatcctttacttca<br>ggcggccgcAACTTAAAAAAAAAAATCAAAatgatgcaatcaagcgcctcctgtgcaaatcctgcggcctgggcacccttctggtggcggtcgtctggctgctggcgctccttttctactcgcacagcctgcgca<br>gctccatccgctcggcaggatggaggatcgacgaagggaacgcaacgccacgggccgagctgagctaccaggccagggtcaccgtcggatgcactccaaatgccagcactaccactggcgaatcgccggcggcacc<br>caagccgcccagtgatccggagcaactggaactgttgggtgtggtgcgcaacaagcaggacaagtacatacgcgacattggctacaagcatcacgccttcaacgcgctggtgtccaataacataggactcttccgggcga<br>tcccagacactcggcacaaggtgtgcgatcgccaggagaccacggaagcggaaaacctgccccaggcctccatagtcatgtgcttctacaacgagcacaagatgacgctgatgcgttcgattaagacggtgctggaacg<br>cacgcccagctacctgctgcgcgagattattctggtggatgatcacagcgatcttccagaattggaattccatttgcacggggatctgcgtgcccgcctgaagtacgacaacctgcggtacatcaaaaacgagcagcgcgag<br>ggtctcattcgttcccgggtcattggtgcacgcgaggccgtgggcgatgtgctcgtcttcctcgactcgcatatcgaggtcaatcagcagtggctggagccgctgctgcggctcatcaagtcggagaacgccactctggccgtg<br>cccgtcgccgacctgatcaacgctgatacctttgagtacacgcccagtccactagtgcgcggaggattcaactggggcctgcacttccgctgggagaatctgcccgagggcacgctcaaggtgccggaggattttcgcgga<br>cccttccgttcgcccaccatggccggcggagcctttgccgtgaacaggaagtacttccagcatctgggcgagtacgacatggccatggatatatggggtggcgagaatatagagatctccttcagagcctggcagtgtggtgg<br>cgccattaagatcgttccctgctcccgtgtcggtcacatattccgcaaacgacgtccctacacttcgccagatggagccaacaccatgttgaagaactcactgcgtttggcgcacgtctggatggatcagtacaaggattactat<br>ctcaagcacgaaaaggtgcccaagacctatgattatggagacattagcgatcgcctcaagcttagggagcgtctgcagtgcagggactttgcctggtatctaaagaatgtctatccagagctgcatgtgccgggcgaggagt<br>ccaagaagtcggcggcagcgcccatctttcagccttggcactcgcgaaaacgtaactatgtcgacacattccagctgcgactgacgggaacggagctctgtgccgccgtggtggctcccaaggtgaagggtttctggaaga<br>agggcagctccctgcaattgcaaacctgccgaaggacacccaatcagctgtggtacgaaacggagaaggcggaaattgtgctggacaagttgctgtgcctggaggcatccggggatgcgcaggtcacggtcaacaagt<br>gccacgagatgctcggcgatcagcagtggcggcacactcgcaacgccaacagtcccgtctacaacatggccaaaggcacctgcctccgggcagctgctcccacgacgggcgccctcatcagtctggatctgtgctccaa<br>gtccaacggagcgggcggctcctgggacattgtgcagctgaagaagccaacggaggcggagggcagggccaaggaggcgaggaactcggacaaggcgctgTACACTGATATCGAAATGAACCGC<br>CTGGGTAAGTAGtctagaatgaatcgttttaaaataacaaatcaattgttttatatattcgtacgattctttgattatgtaataaaatgtgatcattaggaagattacgaaaaatataaaaaatatgagttctgtgtgtataac<br>aaatgctgtaaacgccacaattgtgtttgttgcaaataaacccatgattatttgattaaaattgttgttttcttgttcatagacaatagtgtgttttgcctaaacgtgtactgcataaactccatgcgagtgtatagcgagctagtggctaa<br>cgcttgccccaccaaagtagattcgtcaaaatcctcaatttcatcaccctcctccaagtttaacatttggccgtcggaattaacttctaaagatgccacataatctaataaatgaaatagagattcaaacgtggcgtcatcgtccgtt<br>tcgaccatttccgaaaagaactcgggcataaactctatgatttctctggacgtggtgttgtcgaaactctcaaagtacgcagtcaggaacgtgcgcgacatgtcgtcgggaaactcgcgcggaaacatgttgttgtaaccgaac<br>gggtcccatagcgccaaaaccaaatctgccagcgtcaatagaatgagcacgatgccgacaatggagctggcttggatagcgattcgagttaacggccggccgtttaaacgaattcttgaagacgaaaggcctcgtgata<br>cgcctatttttataggttaatgtcatgataataatggtttcttagactcaggtggcacttttcggggaaatgtgcgcggaacccctatttgtttatttttctaaatacattcaaatatgtatccgctcatgagacaataaccctgataaatgct<br>tcaataatattgaaaaaggaagagtatgagtattcaacatttccgtgtcgcccttattcccttttttgcggcattttgccttcctgtttttgctcacccagaaacgctggtgaaagtaaaagatgctgaagatcagttgggtgcacgagt<br>gggttacatcgaactggatctcaacagcggtaagatccttgagagttttcgccccgaagaacgttttccaatgatgagcacttttaaagttctgctatgtggcgcggtattatcccgtattgacgccgggcaagagcaactcggtc<br>gccgcatacactattctcagaatgacttggttgagtactcaccagtcacagaaaagcatcttacggatggcatgacagtaagagaattatgcagtgctgccataaccatgagtgataacactgcggccaacttacttctgacaa<br>cgatcggaggaccgaaggagctaaccgcttttttgcacaacatgggggatcatgtaactcgccttgatcgttgggaaccggagctgaatgaagccataccaaacgacgagcgtgacaccacgatgcctgtagcaatggca<br>acaacgttgcgcaaactattaactggcgaactacttactctagcttcccggcaacaattaatagactggatggaggcggataaagttgcaggaccacttctgcgctcggcccttccggctggctggtttattgctgataaatctgg<br>agccggtgagcgtgggtctcgcggtatcattgcagcactggggccagatggtaagccctcccgtatcgtagttatctacacgacggggagtcaggcaactatggatgaacgaaatagacagatcgctgagataggtgcctc<br>actgattaagcattggtaactgtcagaccaagtttactcatatatactttagattgatttaaaacttcatttttaatttaaaaggatctaggtgaagatcctttttgataatctcatgaccaaaatcccttaacgtgagttttcgttccactgag<br>cgtcagaccccgtagaaaagatcaaaggatcttcttgagatcctttttttctgcgcgtaatctgctgcttgcaaacaaaaaaaccaccgctaccagcggtggtttgtttgccggatcaagagctaccaactctttttccgaaggtaa<br>ctggcttcagcagagcgcagataccaaatactgttcttctagtgtagccgtagttaggccaccacttcaagaactctgtagcaccgcctacatacctcgctctgctaatcctgttaccagtggctgctgccagtggcgataagtcgt<br>gtcttaccgggttggactcaagacgatagttaccggataaggcgcagcggtcgggctgaacggggggttcgtgcacacagcccagcttggagcgaacgacctacaccgaactgagatacctacagcgtgagctatgaga<br>aagcgccacgcttcccgaagggagaaaggcggacaggtatccggtaagcggcagggtcggaacaggagagcgcacgagggagcttccagggggaaacgcctggtatctttatagtcctgtcgggtttcgccacctctg<br>acttgagcgtcgatttttgtgatgctcgtcaggggggcggagcctatggaaaaacgccagcaacgcggcctttttacggttcctggccttttgctggccttttgctcacatgttaccgtcgacgatgtaggtcacggtctcgaagccg<br>cggtgcgggtgccagggcgtgcccttgggctccccgggcgcgtactccacctcacccatctggtccatcatgatgaacgggtcgaggtggcggtagttgatcccggcgaacgcgcggcgcaccgggaagccctcgccctc<br>gaaaccgctgggcgcggtggtcacggtgagcacgggacgtgcgacggcgtcggcgggtgcggatacgcggggcagcgtcagcgggttctcgacggtcacggcgggcatgtcgacaagccgaacatatggcgcgcc<br>tagtatgtatgtaagttaataaaacccattttgcggaaagtagataaaaaaaacatttttttttactgcactggatatcattgaacttatctgatcagttttaaatttacttcgatccaagggtatttgatgtaccaggttctttcgattacct<br>ctcactcaaaatgacattccactcaaagtcagcgctgtttgcctccttctctgtccacagaaatatcgccgtctctttcgccgctgcgtccgctatctctttcgccaccgtttgtagcgttacgtagcgtcaatgtccgccttcagttgcatt<br>ttgtcagcggtttcgtgacgaagctccaagcggtttacgccatcaattaaacacaaagtgctgtgccaaaactcctctcgcttcttatttttgtttgtttttgagtgattggggtggtgattggttttgggtgggtaagcaggggaaagtg<br>tgaaaaatcccggcaatgggccaagaggatcaggagctattaattcgcggaggcagcaaacacccatctgccgagcatctgaacaatgtgagtagtacatgtgcatacatcttaaggaagttatgcccggatctaattcaatt<br>agagactaattcaattagagctaattcaattagcgccggagtataaatagaggcgcttcgtctacggagcgacaattcaattcaaacaagcaaagtgaacacgtcgctaagcgaaagctaagcaaataaacaagcgcag<br>ctgaacaagctaaacaatcggcccatggtgagcaagggcgaggaggataacatggccatcatcaaggagttcatgcgcttcaaggtgcacatggagggctccgtgaacggccacgagttcgagatcgagggcgaggg<br>cgagggccgcccctacgagggcacccagaccgccaagctgaaggtgaccaagggtggcccctgcccttcgcctgggacatcctgtcccctcagttcatgtacggctccaaggcctacgtgaagcaccccgccgacatc<br>cccgactacttgaagctgtccttccccgagggcttcaagtgggagcgcgtgatgaacttcgaggacggcggcgtggtgaccgtgacccaggactcctccctgcaggacggcgagttcatctacaaggtgaagctgcgcggc<br>accaacttcccctccgacggccccgtaatgcagaagaagaccatgggctgggaggcctcctccgagcggatgtaccccgaggacggcgccctgaagggcgagatcaagcagaggctgaagctgaaggacggcggc<br>cactacgacgctgaggtcaagaccacctacaaggccaagaagcccgtgcagctgcccggcgcctacaacgtcaacatcaagttggacatcacctcccacaacgaggactacaccatcgtggaacagtacgaacgcgc<br>cgagggccgccactccaccggcggcatggacgagctgtacaagtaagcgccgcgactctgatcataatcagccataccacatttgtagaggttttacttgctttaaaaaacctcccacacctccccctgaacctgaaacataa<br>aatgaatgcaattgttgttgttaacttgtttattgcagcttataatggttacaaataaagcaatagcatcacaaatttcacaaataaagcatttttttcactgcattctagttgtggtttgtccaaactcatcaatgtatcttagggcccgacg<br>caaggagtagccgacatatatccgaaataactgcttgtttttttttaccattattaccatcgtgtttactgtttattgccccctcaaaaagctaatgtaattatatttgtgccaataaaaacaagatatgacctatagaatacaagtatttc |

| | |
|---|---|
| | cccttcgaacatccccacaagtagactttggatttgtcttctaaccaaaagacttacacacctgcataccttacatcaaaaactcgtttatcgctacataaaacaccgggatatattttttatatacatacttttcaaatcgcgcgccct cttcataattcacctccaccacaccacgtttcgtagttgctctttcgctgtctcccacccgctctccgcaacacattcaccttttgttcgacgaccttggagcgactgtcgttagttccgcgcgattcggttcgctcaaatggttccgagtg gttcatttcgtctcaatagaaattagtaataaatatttgtatgtacaatttatttgctccaatatatttgtatatatttccctcacagctatatttattctaatttaatattatgactttttaaggtaatttttgtgacctgttcggagtgattagcgttac aatttgaactgaaagtgacatccagtgtttgttccttgtgtagatgcatctcaaaaaaatggtgggcataatagtgttgtttatatatatcaaaaataacaactataataataagaatacatttaatttagaaaatgct |
